## Supplementary figures and images for "A population modification gene drive targeting both *Saglin* and *Lipophorin* impairs *Plasmodium* transmission in *Anopheles* mosquitoes"

### Supplemental Figure 1

#### Suppl. Figure 1

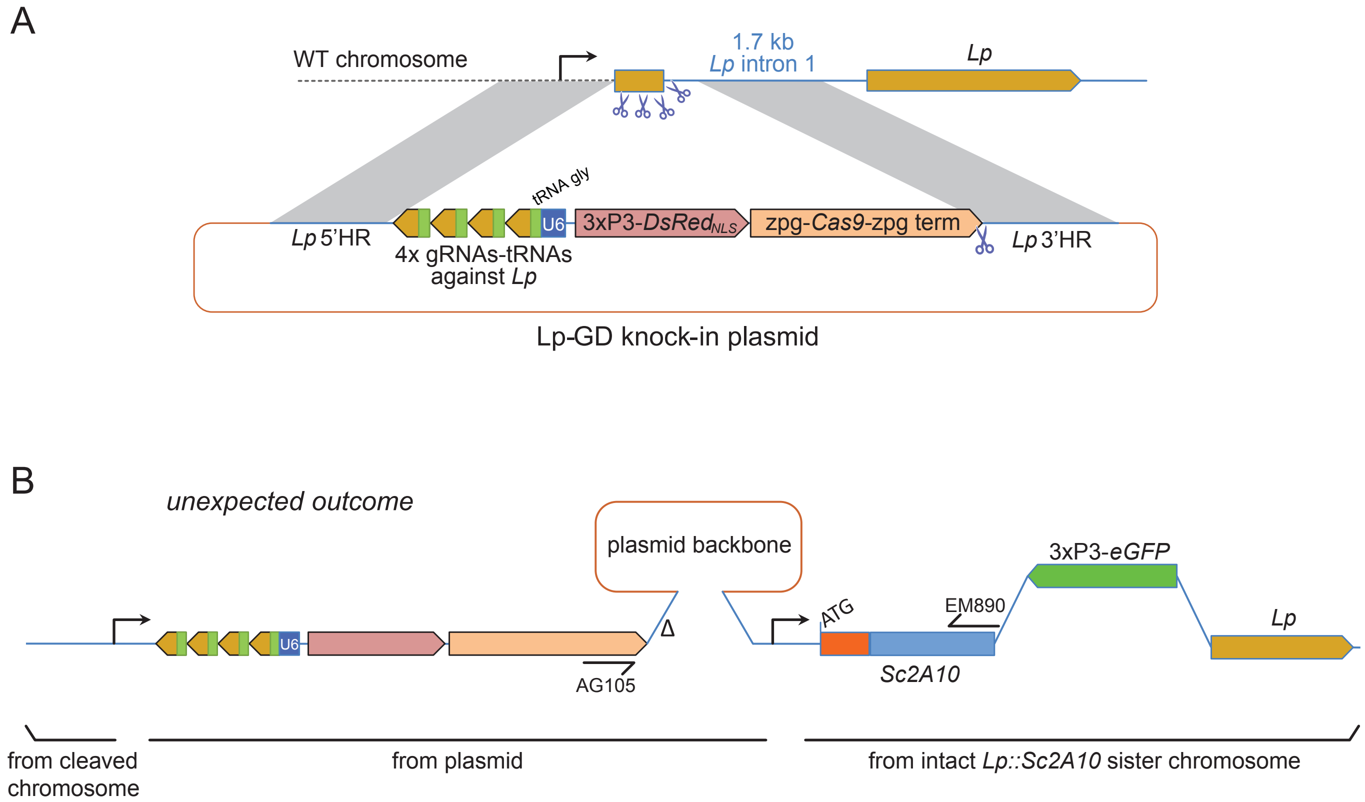

### Supplemental Figure 2

Suppl. Figure 2

A

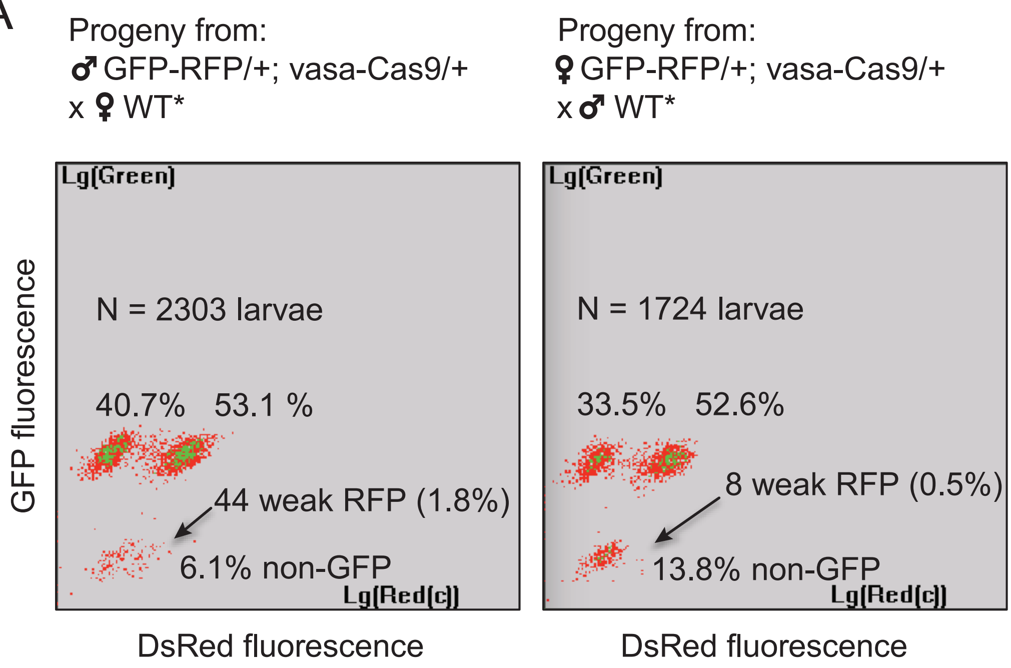


B

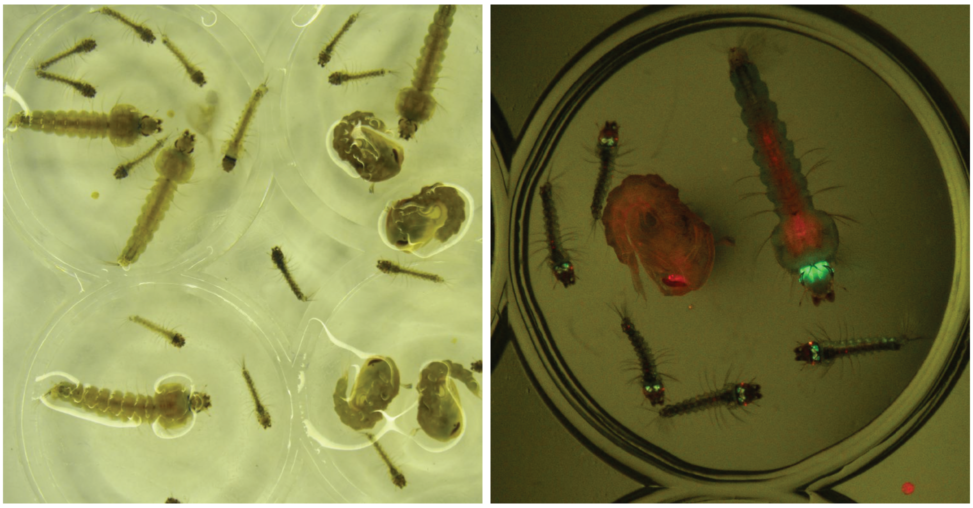
