## Supplemental File 1 for "A population modification gene drive targeting both *Saglin* and *Lipophorin* impairs *Plasmodium* transmission in *Anopheles* mosquitoes"

```
Supplemental File 1: DNA sequences

Supplemental file 1A : Lp::Sc2A10 knock-in plasmid

LOCUS       pENTR3xgRNALp-Sc2A10  8489 bp    DNA     circular       FEATURES             Location/Qualifiers
     misc_feature    537..552
                     /note="M13F"
     misc_feature    complement(6750..6768)
                     /note="M13R"
     misc_feature    3637..3645
                     /note=""
     misc_feature    3659..3736
                     /note="re-coded signal peptide"
                     /note=""
     misc_feature    3737..4090
                     /note="heavy chain_2A10%E2%80%9D misc_feature
                     3737..4465 /note=%222A10"
     misc_feature    4502..4507
                     /note="perfect intron splice acceptor site"
     misc_feature    4475..4486
                     /note="cleavage site"
     misc_feature    4466..4498
                     /note="encompasses cleavage site"
     misc_feature    complement(4520..5668)
                     /note="3xP3-GFP-pAtub56D"
     misc_feature    4499..4519
                     /note="PRESERVE"
     misc_feature    4097..4141
                     /note="(G4S)3-linker"
     misc_feature    4142..4465
                     /note="light chain_2A10%E2%80%9D misc_feature
                     4487..4507 /note=%22peptide added to ApoII"
     misc_feature    complement(4520..4688)
                     /note="Dm beta-Tub56D terminator"
     misc_feature    4469..4687
                     /note="gBlock"
     misc_feature    4689..5407
                     /note="GFP"
     misc_feature    complement(4575..4580)
                     /note="polyA site"
     misc_feature    5619..5668
                     /note="3xPax6 binding sites"
     misc_feature    5572..5579
                     /note="TATA box"
     misc_feature    6688..6688
                     /note="mutated to T"
     misc_feature    2133..3652
                     /note="Lp promoter 5%82%C4%F4flk HA"
     misc_feature    complement(906..1226)
                     /note="AGAP013557 U6 prom"
     misc_feature    complement(810..881)
                     /note="pX330 chimeric guide RNA scaffold"
     misc_feature    complement(875..881)
                     /note="CRISPR  end"
     misc_feature    complement(807..870)
                     /note="tracer"
     misc_feature    774..809
                     /note="from Ag U6"
     misc_feature    complement(1359..1679)
                     /note="AGAP013557 U6 prom"
     misc_feature    complement(1263..1334)
                     /note="pX330 chimeric guide RNA scaffold"
     misc_feature    complement(1328..1334)
                     /note="CRISPR  end"
     misc_feature    complement(1260..1323)
                     /note="tracer"
     misc_feature    1231..1262
                     /note="from Ag U6"
     misc_feature    complement(1812..2132)
                     /note="AGAP013557 U6 prom"
     misc_feature    complement(1716..1787)
                     /note="pX330 chimeric guide RNA scaffold"
     misc_feature    complement(1781..1787)
                     /note="CRISPR  end"
     misc_feature    complement(1713..1776)
                     /note="tracer"
     misc_feature    1684..1715
                     /note="from Ag U6"
     misc_feature    complement(882..905)
                     /note="sgRNA1 EM1096"
     misc_feature    1335..1358
                     /note="sgRNA2 EM1098"
     misc_feature    1788..1811
                     /note="sgRNA3 EM1100"
     misc_feature    complement(6701..6726)
                     /note="EM1103 gRNA2 target"
     misc_feature    5671..6697
                     /note="Lp 3%82%C4%F4flk HA"
ORIGIN
        1 CTTTCCTGCG TTATCCCCTG ATTCTGTGGA TAACCGTATT ACCGCCTTTG AGTGAGCTGA
       61 TACCGCTCGC CGCAGCCGAA CGACCGAGCG CAGCGAGTCA GTGAGCGAGG AAGCGGAAGA
      121 GCGCCCAATA CGCAAACCGC CTCTCCCCGC GCGTTGGCCG ATTCATTAAT GCAGCTGGCA
      181 CGACAGGTTT CCCGACTGGA AAGCGGGCAG TGAGCGCAAC GCAATTAATA CGCGTACCGC
      241 TAGCCAGGAA GAGTTTGTAG AAACGCAAAA AGGCCATCCG TCAGGATGGC CTTCTGCTTA
      301 GTTTGATGCC TGGCAGTTTA TGGCGGGCGT CCTGCCCGCC ACCCTCCGGG CCGTTGCTTC
      361 ACAACGTTCA AATCCGCTCC CGGCGGATTT GTCCTACTCA GGAGAGCGTT CACCGACAAA
      421 CAACAGATAA AACGAAAGGC CCAGTCTTCC GACTGAGCCT TTCGTTTTAT TTGATGCCTG
      481 GCAGTTCCCT ACTCTCGCGT TAACGCTAGC ATGGATGTTT TCCCAGTCAC GACGTTGTAA
      541 AACGACGGCC AGTCTTAAGC TCGGGCCCCT ACAGGTCACT AATACCATCT AAGTAGTTGA
      601 TTCATAGTGA CTGGATATGT TGTGTTTTAC AGTATTATGT AGTCTGTTTT TTATGCAAAA
      661 TCTAATTTAA TATATTGATA TTTATATCAT TTTACGTTTC TCGTTCAACT TTTCTATACA
      721 AAGTTGGTAC CGGGCCCCCC GCTAGCGTCG ACGGTATCGA TAAGCTTGAT cggatccGAT
      781 GAAAATAAGA AAAACATTTG ACAAAAAAAg caccgactcg gtgccacttt ttcaagttga
      841 taacggacta gccttatttt aacttgctaT TTCtagctct aaaacGGTGT CGCTAGTGCT
      901 GATTCAAGGA CGAGGGGAAA AAAGGTTGTA TATATACTTT GCGCTTTCAA TCCTTGCTCT
      961 AGCGATGCAT AAAGGATATT CAAGAAGGAT TTTTTGCCTG AGTGTGACTG TTATAAGGTT
     1021 TTGATCCTGC CGTAACTCAC GCTAATGTGA TGTTTCATAA ACTTTTCAAC ATTTCTTGTA
     1081 TTGCTTCATC TCTTTTTTAA TCAAAAGTTA CTATTCATTG TAATACGTTG AAATATCAAA
     1141 AAAGAATAAA CAGTTTTCCT ATGATTAATT CAAAAACAGA GTTTTCTAGA ATAACCAAGA
     1201 GCCAAACCGC TTCAGTGTAT ATGTGGggaa GATGAAAATA AGAAAAACAT TTGACAAAAA
     1261 AAgcaccgac tcggtgccac tttttcaagt tgataacgga ctagccttat tttaacttgc
     1321 taTTTCtagc tctaaaacTC CCTGACGATA TTTCACGCAA GGACGAGGGG AAAAAAGGTT
     1381 GTATATATAC TTTGCGCTTT CAATCCTTGC TCTAGCGATG CATAAAGGAT ATTCAAGAAG
     1441 GATTTTTTGC CTGAGTGTGA CTGTTATAAG GTTTTGATCC TGCCGTAACT CACGCTAATG
     1501 TGATGTTTCA TAAACTTTTC AACATTTCTT GTATTGCTTC ATCTCTTTTT TAATCAAAAG
     1561 TTACTATTCA TTGTAATACG TTGAAATATC AAAAAAGAAT AAACAGTTTT CCTATGATTA
     1621 ATTCAAAAAC AGAGTTTTCT AGAATAACCA AGAGCCAAAC CGCTTCAGTG TATATGTGGa
     1681 gagGATGAAA ATAAGAAAAA CATTTGACAA AAAAAgcacc gactcggtgc cactttttca
     1741 agttgataac ggactagcct tattttaact tgctaTTTCt agctctaaaa cCCGAGGACC
     1801 CACATCGTTC CAAGGACGAG GGGAAAAAAG GTTGTATATA TACTTTGCGC TTTCAATCCT
     1861 TGCTCTAGCG ATGCATAAAG GATATTCAAG AAGGATTTTT TGCCTGAGTG TGACTGTTAT
     1921 AAGGTTTTGA TCCTGCCGTA ACTCACGCTA ATGTGATGTT TCATAAACTT TTCAACATTT
     1981 CTTGTATTGC TTCATCTCTT TTTTAATCAA AAGTTACTAT TCATTGTAAT ACGTTGAAAT
     2041 ATCAAAAAAG AATAAACAGT TTTCCTATGA TTAATTCAAA AACAGAGTTT TCTAGAATAA
     2101 CCAAGAGCCA AACCGCTTCA GTGTATATGT GGAACAGTTC ATTCCCGATT GAGGGATTTT
     2161 ATTCCCCGGG GGCCTTTTCA AACGGCTTAA TATAAGCAAT TAATAGTATT TTTTCTTTCA
     2221 GGTTAGTTTA CTGTAATGGT GTAATTGTCA TCTTACACCT CCGTCTGATA AGAGATTACG
     2281 AAGCTCAGTA TGATGAAATA AATAAGATAA ATTTATTTAA AAAAGAACAA TTGCTATGAG
     2341 AGTGAAATAC AACAGTGGCG TTCACAATAT TCGAAAAACA ATAAAATTAA AAAAAAAACA
     2401 AGAAAACATT CACAAACATA TCAATCTGCT TTCATCGACA CCGAACTGCT AGCCTCCCCA
     2461 GTCTAACCGC GGTGGGGACG TTTAATTGCC TTTGTTCTCG CACCCGGTCA AACATACACT
     2521 TCGGACCTTG CTCCGAACCC CACTGTAATC CCTAGCTCGT CATCATCATT GCCGGCATCA
     2581 TGCTAAGCGT GCATTATTTT CACAACTTAG CGTAATGCTA GCGTGCGCTA GCAACAAACT
     2641 CGCCCGCAGA CTCGTCACAG CACCGGTACG ATCGATCGTT TACCGTTCCC TTTCCCGATC
     2701 GGGTTGGCTG CGATATCCGT GTCCGGGTAG AAAACTTCCC CTTTTACACA CACACACTCA
     2761 CATACACACA GAGCTGAATA GCAACTTACC TTATCTGTTC GTCATCGCTC GGCCGGATCT
     2821 GGACGAATCT TCGCACCGAT AACCATGTGG ATCTACGACC TCCGCTTGGC TGTCTCTCTG
     2881 CTCATGTGTA TGTCTGTGTG TGTGTGTATG TGAGCTTCTT CCCTCAAATC CCTCGATCTC
     2941 GCTGTGCCAA CAATCAAACG TGCAAGTGCA AACATTGCAC CCCATTGATT ATACACCAAC
     3001 ACCAACCAAT TCCCCTTGCG GAGGCATCTC TGTGCTCGGC AGCATGTTTA CCGCAGATCT
     3061 ACAGAGAACT TCAATTGAGG TCCTTTCCAC CCCCAGCCCT CAACCGGCAA TCCGGCAGCC
     3121 ACTGGATCAT CAGCGAAAGA GAGAGAGAGC AGAGCAGAAC AGAGGTGACC AACTGTGGTA
     3181 TCGCTTCCCG CGCGCCGGTG TGTTGGTGTC CATTTCGGTG ATCGCGATCC CGGCCGCTTC
     3241 CAGCACCGTC CACCGATCAG TCACAAAAAC GCTCTCCAAA CCCCTTATCA GCACCGTTCG
     3301 CTGATGTGAA CCCCCGTTCA AACCCCAAAT GCAGTGTTTG TATTGCTGTG TGTATGTGTA
     3361 CGTGCGTGTG TGTGGAAATT CTATAAAAGT AGGCACCCGT GGCCGGGATC CGTTATTCCC
     3421 GTCCTGAGGC CCGCCCGGGA TCGCTGGTGA CGACAGACGA GCCGCTGTGT GACGTACGTA
     3481 GTGCCCGATC GGTAAAGAGT GAACCGTCTT CTCTGCAGTG TAGGAGAGAA CGGTTTCATC
     3541 TTTTTCGCCC ACACCCCCCC GGTTACATTC CATGTTGAAC TGTAAGGTCT AGTGAACATT
     3601 TCCTGAGTGT GGAAAGTGTG GTTTAGTGCG TGAGAGTGCA CGGACACGAC ACGGAACGAT
     3661 GTGGGTgCTg GGaGGcAGaA GaCTcCTgTG GtcgTTtCTc GTcagctTgG TcCTcATcCA
     3721 gtccGTcagC gcagccCAGA TCCAGCTGGT GCAGTCGGGC CCCGAGCTGA AGAAGCCCGG
     3781 CGAAACGGTG AAGATCAGCT GTAAGGCGAG CGGTTACACC TTCACCAACT ACGGCATCAA
     3841 CTGGGTGAAG CAGGCGCCGG GCAAGGGTCT GAAGTGGATG GGATGGATCA ACACGATCAC
     3901 CGAGGAGCCC ACCTTCGCCG AAGAGTTCAC GGGACGCTTC GCCTTCAGCC TGGAAACCTC
     3961 GGCGAGCACC GCCTACCTCC AGATCAACAA CCTGAAGAAC GAGGACACGG CCACGTACTT
     4021 CTGTGCGCGC GGCAGCGAGT TCGGACGCCT GGTCTACTGG GGCCAGGGAG CGAGCGTGAC
     4081 CGTCTCGAGC TCGACCGGTG GTGGTGGCAG CGGCGGAGGA GGTAGCGGAG GTGGCGGCAG
     4141 CGACATCCAG ATGACGCAGA CGACGAGCTC GCTGTCGGCC TCGCTGGGCG ATCGTGTGAC
     4201 GATCTCGTGC AGCGCCTCGC AGGGCATCAG CAACTACCTG AACTGGTACC AGCAGAAGCC
     4261 GGACGGTACC GTCAAGCTGC TGATCTTCTA CACCTCGACC CTGTACTCGG GTGTGCCGTC
     4321 GCGTTTCTCG GGTTCGGGCT CGGGCACCGA TTACTCGCTG ACCATCTCGA ACCTGGAGCC
     4381 CGAGGACATC GCGACGTACT ACTGCCAGCA GTACTCGCGT TTCCCGTACG TGTTCGGCGG
     4441 CGGCACGAAG CTGGAGATCA AGCGCGCcAA GGAaCGtTTC CGtCGCGGaA TtCGtGAaTC
     4501 CGCAGGTATG TTCCTATACg aaaccccaac aaaaaccata attgtttAGA CTTGTGAACA
     4561 AAATTGGATC CGACTTTATT GATTACGTTG TTAAGAGAAC AAATCTTTTA CAACTGAATT
     4621 CATTTGTTCT CGTTTCATTT TTTTTCGCAA AACATTGATC GAGAATTCGA TTGATTTCCG
     4681 ATTCGAATTT ACTTGTACAG CTCGTCCATG CCGAGAGTGA TCCCGGCGGC GGTCACGAAC
     4741 TCCAGCAGGA CCATGTGATC GCGCTTCTCG TTGGGGTCTT TGCTCAGGGC GGACTGGGTG
     4801 CTCAGGTAGT GGTTGTCGGG CAGCAGCACG GGGCCGTCGC CGATGGGGGT GTTCTGCTGG
     4861 TAGTGGTCGG CGAGCTGCAC GCTGCCGTCC TCGATGTTGT GGCGGATCTT GAAGTTCACC
     4921 TTGATGCCGT TCTTCTGCTT GTCGGCCATG ATATAGACGT TGTGGCTGTT GTAGTTGTAC
     4981 TCCAGCTTGT GCCCCAGGAT GTTGCCGTCC TCCTTGAAGT CGATGCCCTT CAGCTCGATG
     5041 CGGTTCACCA GGGTGTCGCC CTCGAACTTC ACCTCGGCGC GGGTCTTGTA GTTGCCGTCG
     5101 TCCTTGAAGA AGATGGTGCG CTCCTGGACG TAGCCTTCGG GCATGGCGGA CTTGAAGAAG
     5161 TCGTGCTGCT TCATGTGGTC GGGGTAGCGG CTGAAGCACT GCACGCCGTA GGTCAGGGTG
     5221 GTCACGAGGG TGGGCCAGGG CACGGGCAGC TTGCCGGTGG TGCAGATGAA CTTCAGGGTC
     5281 AGCTTGCCGT AGGTGGCATC GCCCTCGCCC TCGCCGGACA CGCTGAACTT GTGGCCGTTT
     5341 ACGTCGCCGT CCAGCTCGAC CAGGATGGGC ACCACCCCGG TGAACAGCTC CTCGCCCTTG
     5401 CTCACCATGG TGGCGACCGG TGGATCCCGG GCCCGCGGTA CCGTCGACTC TAGCGGTACC
     5461 CCGATTGTTT AGCTTGTTCA GCTGCGCTTG TTTATTTGCT TAGCTTTCGC TTAGCGACGT
     5521 GTTCACTTTG CTTGTTTGAA TTGAATTGTC GCTCCGTAGA CGAAGCGCCT CTATTTATAC
     5581 TCCGGCGGTC GAGGGTTCGA AATCGATAAG CTTGGATCCT AATTGAATTA GCTCTAATTG
     5641 AATTAGTCTC TAATTGAATT AGATCCCCGA TGGAAGAGAT GGCGAAGGTT CTCATCCACA
     5701 AGTTGTTGGT GATGCTAACA TTGCTTCCGA GCAATGTATT GCTGCCGAGG GTGTCGATTA
     5761 GTACAGCAAA GCGCGAGAAA CAAGTGGCTG CTGTGGCCGG CGGTACCCAT AAACCAGTTG
     5821 AGATAATTTA GACAATAAAT TGTAATAGGC AATTTGATGA CGTCCCGCAT GTACGGATCG
     5881 TCAACGCTGC ACGCTCGGGA ATAGGGTGAG AAGTGCTGCC CGTTTGTTGG ACAGCGCAGG
     5941 TGGTGATCGT GTTGTTTTGG TATCACGAAA ACAAGATAGA ATCTGAGAAA AAAGAACCTT
     6001 TTTATCACTT TACCCGCACC GATACTGGTC TTTCTTAGAT CATTTCGGGG CGAAGCAAAC
     6061 AAGTCAGCAC GGCATACAAA TTTTGGGATA ATCCCTTAGT CAGTGCCTGT GTCTTTCTAA
     6121 TGAAGATTGA CAATGGCGGC GGCGGCGGCG GTGGTAGCAA CCAACGCACT GGCCTTTGTT
     6181 TTCATACTTT TCAATGAATT CAATTAGCTT AACCAACGAG GTTGTTGTTG ATGTTCTCAC
     6241 GGAAGTCAGG AGAAGCTCCG ACGGGCGTGT AGGGTAAGTC GCAGTTCGGC AGCATTCTGG
     6301 CAGTGTACAC CCAGCACACC GAGCGTTCTA GAACGCTGAA TCGTTGAAAC AGAGCAAATG
     6361 ACTGAAGGGC TGTGTCTATT GCTGGGCTGT ACAGAATGAC GCATTTCGTG TGGCAATGTG
     6421 TTTTCGAAAG ATTGAAGCAT TCTGATAGAG CCGTGTGTCA GGAATGTAAA TATTTAGATA
     6481 CTGCAGCTAA CTACAAACTG AGAGAGCTTT CAGCTGGCAG GTGTCCCTAC TTGGTGGTCT
     6541 TTTCTAGAAA GAATGTAGAA TTGACATGTT GTGATGATTT ACCAAAAACA ATTTGAGAAG
     6601 CTAGAGAGTG AATCGTTCTA GCCACCGGAA CCATCTGGAA TGTTGAAGAT TTTTCTGGAA
     6661 TATTAGTAGA CAAAAAGAAG GAGGTCTCAA CTCCGAGctt GCGTGAAATA TCGTCAGGGA
     6721 CGgcttATCC CCTATAGTGA GTCGTATTAC ATGGTCATAG CTGTTTCCTG GCAGCTCTGG
     6781 CCCGTGTCTC AAAATCTCTG ATGTTACATT GCACAAGATA AAAATATATC ATCATGAACA
     6841 ATAAAACTGT CTGCTTACAT AAACAGTAAT ACAAGGGGTG TTATGAGCCA TATTCAACGG
     6901 GAAACGTCGA GGCCGCGATT AAATTCCAAC ATGGATGCTG ATTTATATGG GTATAAATGG
     6961 GCTCGCGATA ATGTCGGGCA ATCAGGTGCG ACAATCTATC GCTTGTATGG GAAGCCCGAT
     7021 GCGCCAGAGT TGTTTCTGAA ACATGGCAAA GGTAGCGTTG CCAATGATGT TACAGATGAG
     7081 ATGGTCAGAC TAAACTGGCT GACGGAATTT ATGCCTCTTC CGACCATCAA GCATTTTATC
     7141 CGTACTCCTG ATGATGCATG GTTACTCACC ACTGCGATCC CCGGAAAAAC AGCATTCCAG
     7201 GTATTAGAAG AATATCCTGA TTCAGGTGAA AATATTGTTG ATGCGCTGGC AGTGTTCCTG
     7261 CGCCGGTTGC ATTCGATTCC TGTTTGTAAT TGTCCTTTTA ACAGCGATCG CGTATTTCGT
     7321 CTCGCTCAGG CGCAATCACG AATGAATAAC GGTTTGGTTG ATGCGAGTGA TTTTGATGAC
     7381 GAGCGTAATG GCTGGCCTGT TGAACAAGTC TGGAAAGAAA TGCATAAACT TTTGCCATTC
     7441 TCACCGGATT CAGTCGTCAC TCATGGTGAT TTCTCACTTG ATAACCTTAT TTTTGACGAG
     7501 GGGAAATTAA TAGGTTGTAT TGATGTTGGA CGAGTCGGAA TCGCAGACCG ATACCAGGAT
     7561 CTTGCCATCC TATGGAACTG CCTCGGTGAG TTTTCTCCTT CATTACAGAA ACGGCTTTTT
     7621 CAAAAATATG GTATTGATAA TCCTGATATG AATAAATTGC AGTTTCATTT GATGCTCGAT
     7681 GAGTTTTTCT AATCAGAATT GGTTAATTGG TTGTAACACT GGCAGAGCAT TACGCTGACT
     7741 TGACGGGACG GCGCAAGCTC ATGACCAAAA TCCCTTAACG TGAGTTACGC GTCGTTCCAC
     7801 TGAGCGTCAG ACCCCGTAGA AAAGATCAAA GGATCTTCTT GAGATCCTTT TTTTCTGCGC
     7861 GTAATCTGCT GCTTGCAAAC AAAAAAACCA CCGCTACCAG CGGTGGTTTG TTTGCCGGAT
     7921 CAAGAGCTAC CAACTCTTTT TCCGAAGGTA ACTGGCTTCA GCAGAGCGCA GATACCAAAT
     7981 ACTGTTCTTC TAGTGTAGCC GTAGTTAGGC CACCACTTCA AGAACTCTGT AGCACCGCCT
     8041 ACATACCTCG CTCTGCTAAT CCTGTTACCA GTGGCTGCTG CCAGTGGCGA TAAGTCGTGT
     8101 CTTACCGGGT TGGACTCAAG ACGATAGTTA CCGGATAAGG CGCAGCGGTC GGGCTGAACG
     8161 GGGGGTTCGT GCACACAGCC CAGCTTGGAG CGAACGACCT ACACCGAACT GAGATACCTA
     8221 CAGCGTGAGC TATGAGAAAG CGCCACGCTT CCCGAAGGGA GAAAGGCGGA CAGGTATCCG
     8281 GTAAGCGGCA GGGTCGGAAC AGGAGAGCGC ACGAGGGAGC TTCCAGGGGG AAACGCCTGG
     8341 TATCTTTATA GTCCTGTCGG GTTTCGCCAC CTCTGACTTG AGCGTCGATT TTTGTGATGC
     8401 TCGTCAGGGG GGCGGAGCCT ATGGAAAAAC GCCAGCAACG CGGCCTTTTT ACGGTTCCTG
     8461 GCCTTTTGCT GGCCTTTTGC TCACATGTT
//


Supplemental File 1B: Lp::2A10 knockin in genomic context


LOCUS       Lp-2A10                 14919 bp    DNA     linear
FEATURES             Location/Qualifiers
     misc_feature    1..1622
                     /note="Lp promoter"
     exon            5369..9341
                     /note="exon_id=AGAP001826-RA-E2.1"
     misc_feature    7396..7407
                     /note="cleavage site"
     exon            9406..9535
                     /note="exon_id=AGAP001826-RA-E3.1"
     exon            9610..14919
                     /note="exon_id=AGAP001826-RA-E4.1"
     misc_feature    2472..2477
                     /note="perfect intron splice acceptor site"
     misc_feature    2445..2456
                     /note="cleavage site"
     misc_feature    2436..2468
                     /note="encompasses cleavage site"
     misc_feature    complement(2490..3638)
                     /note="3xP3-GFP-pAtub deletes 3 cas9 target sites in
                     intron"
     misc_feature    1623..1642
                     /note="disabled gRNA target site”
     misc_feature    1630..1649
                     /note="disabled gRNA target site"
     misc_feature    1639..1658
                     /note="disabled gRNA target site"
     misc_feature    1650..1668
                     /note="disabled gRNA target site"
     misc_feature    1641..1659
                     /note="disabled gRNA target site"
     misc_feature    1670..1689
                     /note="disabled gRNA target site"
     misc_feature    1629..1703
                     /note="re-coded signal peptide"
     misc_feature    1683..1703
                     /note="disabled gRNA target site"
     misc_feature    2469..2489
                     /note="PRESERVED around splice junction”
     misc_feature    1707..2060
                     /note=“Sc2A10 heavy chain"
     misc_feature    2067..2111
                     /note="(G4S)3-linker"
     misc_feature    2112..2435
                     /note=“Sc2A10 light chain"
     misc_feature    1707..2435
                     /note=“Sc2A10"
     misc_feature    2457..2477
                     /note="peptide added to ApoII"
     misc_feature    3589..3638
                     /note="3xPax6 binding sites"
     misc_feature    3542..3549
                     /note="TATA box"
     misc_feature    2659..3377
                     /note="GFP"
     misc_feature    complement(2490..2658)
                     /note="Dm beta-Tub56D terminator"
     misc_feature    complement(2545..2550)
                     /note="polyA site"
     misc_feature    complement(4639..4667)
                     /note="EM894 GGTCTC-> GGTCTT"
     misc_feature    4658..4658
                     /note="mutated to T"
     source          1..14919
                     /dnas_title=“Sc2A10 inserted in Lp genomic fg"
ORIGIN
        1 TTGCGGGGAA GACACATTCG AGATACGCTA AGTGATTGAG CGATTACGAT CTAGCAAGAC
       61 ATACGTTCAG CTGTGAGAAT AATCATCCAT CTTTCTGCAA TGAACAGTTC ATTCCCGATT
      121 GAGGGATTTT ATTCCCCGGG GGCCTTTTCA AACGGCTTAA TATAAGCAAT TAATAGTATT
      181 TTTTCTTTCA GGTTAGTTTA CTGTAATGGT GTAATTGTCA TCTTACACCT CCGTCTGATA
      241 AGAGATTACG AAGCTCAGTA TGATGAAATA AATAAGATAA ATTTATTTAA AAAAGAACAA
      301 TTGCTATGAG AGTGAAATAC AACAGTGGCG TTCACAATAT TCGAAAAACA ATAAAATTAA
      361 AAAAAAAACA AGAAAACATT CACAAACATA TCAATCTGCT TTCATCGACA CCGAACTGCT
      421 AGCCTCCCCA GTCTAACCGC GGTGGGGACG TTTAATTGCC TTTGTTCTCG CACCCGGTCA
      481 AACATACACT TCGGACCTTG CTCCGAACCC CACTGTAATC CCTAGCTCGT CATCATCATT
      541 GCCGGCATCA TGCTAAGCGT GCATTATTTT CACAACTTAG CGTAATGCTA GCGTGCGCTA
      601 GCAACAAACT CGCCCGCAGA CTCGTCACAG CACCGGTACG ATCGATCGTT TACCGTTCCC
      661 TTTCCCGATC GGGTTGGCTG CGATATCCGT GTCCGGGTAG AAAACTTCCC CTTTTACACA
      721 CACACACTCA CATACACACA GAGCTGAATA GCAACTTACC TTATCTGTTC GTCATCGCTC
      781 GGCCGGATCT GGACGAATCT TCGCACCGAT AACCATGTGG ATCTACGACC TCCGCTTGGC
      841 TGTCTCTCTG CTCATGTGTA TGTCTGTGTG TGTGTGTATG TGAGCTTCTT CCCTCAAATC
      901 CCTCGATCTC GCTGTGCCAA CAATCAAACG TGCAAGTGCA AACATTGCAC CCCATTGATT
      961 ATACACCAAC ACCAACCAAT TCCCCTTGCG GAGGCATCTC TGTGCTCGGC AGCATGTTTA
     1021 CCGCAGATCT ACAGAGAACT TCAATTGAGG TCCTTTCCAC CCCCAGCCCT CAACCGGCAA
     1081 TCCGGCAGCC ACTGGATCAT CAGCGAAAGA GAGAGAGAGC AGAGCAGAAC AGAGGTGACC
     1141 AACTGTGGTA TCGCTTCCCG CGCGCCGGTG TGTTGGTGTC CATTTCGGTG ATCGCGATCC
     1201 CGGCCGCTTC CAGCACCGTC CACCGATCAG TCACAAAAAC GCTCTCCAAA CCCCTTATCA
     1261 GCACCGTTCG CTGATGTGAA CCCCCGTTCA AACCCCAAAT GCAGTGTTTG TATTGCTGTG
     1321 TGTATGTGTA CGTGCGTGTG TGTGGAAATT CTATAAAAGT AGGCACCCGT GGCCGGGATC
     1381 CGTTATTCCC GTCCTGAGGC CCGCCCGGGA TCGCTGGTGA CGACAGACGA GCCGCTGTGT
     1441 GACGTACGTA GTGCCCGATC GGTAAAGAGT GAACCGTCTT CTCTGCAGTG TAGGAGAGAA
     1501 CGGTTTCATC TTTTTCGCCC ACACCCCCCC GGTTACATTC CATGTTGAAC TGTAAGGTCT
     1561 AGTGAACATT TCCTGAGTGT GGAAAGTGTG GTTTAGTGCG TGAGAGTGCA CGGACACGAC
     1621 ACGGAACGAT GTGGGTgCTg GGaGGcAGaA GaCTcCTgTG GtcgTTtCTc GTcagctTgG
     1681 TcCTcATcCA gtccGTcagC gcagccCAGA TCCAGCTGGT GCAGTCGGGC CCCGAGCTGA
     1741 AGAAGCCCGG CGAAACGGTG AAGATCAGCT GTAAGGCGAG CGGTTACACC TTCACCAACT
     1801 ACGGCATCAA CTGGGTGAAG CAGGCGCCGG GCAAGGGTCT GAAGTGGATG GGATGGATCA
     1861 ACACGATCAC CGAGGAGCCC ACCTTCGCCG AAGAGTTCAC GGGACGCTTC GCCTTCAGCC
     1921 TGGAAACCTC GGCGAGCACC GCCTACCTCC AGATCAACAA CCTGAAGAAC GAGGACACGG
     1981 CCACGTACTT CTGTGCGCGC GGCAGCGAGT TCGGACGCCT GGTCTACTGG GGCCAGGGAG
     2041 CGAGCGTGAC CGTCTCGAGC TCGACCGGTG GTGGTGGCAG CGGCGGAGGA GGTAGCGGAG
     2101 GTGGCGGCAG CGACATCCAG ATGACGCAGA CGACGAGCTC GCTGTCGGCC TCGCTGGGCG
     2161 ATCGTGTGAC GATCTCGTGC AGCGCCTCGC AGGGCATCAG CAACTACCTG AACTGGTACC
     2221 AGCAGAAGCC GGACGGTACC GTCAAGCTGC TGATCTTCTA CACCTCGACC CTGTACTCGG
     2281 GTGTGCCGTC GCGTTTCTCG GGTTCGGGCT CGGGCACCGA TTACTCGCTG ACCATCTCGA
     2341 ACCTGGAGCC CGAGGACATC GCGACGTACT ACTGCCAGCA GTACTCGCGT TTCCCGTACG
     2401 TGTTCGGCGG CGGCACGAAG CTGGAGATCA AGCGCGCcAA GGAaCGtTTC CGtCGCGGaA
     2461 TtCGtGAaTC CGCAGGTATG TTCCTATACg aaaccccaac aaaaaccata attgtttAGA
     2521 CTTGTGAACA AAATTGGATC CGACTTTATT GATTACGTTG TTAAGAGAAC AAATCTTTTA
     2581 CAACTGAATT CATTTGTTCT CGTTTCATTT TTTTTCGCAA AACATTGATC GAGAATTCGA
     2641 TTGATTTCCG ATTCGAATTT ACTTGTACAG CTCGTCCATG CCGAGAGTGA TCCCGGCGGC
     2701 GGTCACGAAC TCCAGCAGGA CCATGTGATC GCGCTTCTCG TTGGGGTCTT TGCTCAGGGC
     2761 GGACTGGGTG CTCAGGTAGT GGTTGTCGGG CAGCAGCACG GGGCCGTCGC CGATGGGGGT
     2821 GTTCTGCTGG TAGTGGTCGG CGAGCTGCAC GCTGCCGTCC TCGATGTTGT GGCGGATCTT
     2881 GAAGTTCACC TTGATGCCGT TCTTCTGCTT GTCGGCCATG ATATAGACGT TGTGGCTGTT
     2941 GTAGTTGTAC TCCAGCTTGT GCCCCAGGAT GTTGCCGTCC TCCTTGAAGT CGATGCCCTT
     3001 CAGCTCGATG CGGTTCACCA GGGTGTCGCC CTCGAACTTC ACCTCGGCGC GGGTCTTGTA
     3061 GTTGCCGTCG TCCTTGAAGA AGATGGTGCG CTCCTGGACG TAGCCTTCGG GCATGGCGGA
     3121 CTTGAAGAAG TCGTGCTGCT TCATGTGGTC GGGGTAGCGG CTGAAGCACT GCACGCCGTA
     3181 GGTCAGGGTG GTCACGAGGG TGGGCCAGGG CACGGGCAGC TTGCCGGTGG TGCAGATGAA
     3241 CTTCAGGGTC AGCTTGCCGT AGGTGGCATC GCCCTCGCCC TCGCCGGACA CGCTGAACTT
     3301 GTGGCCGTTT ACGTCGCCGT CCAGCTCGAC CAGGATGGGC ACCACCCCGG TGAACAGCTC
     3361 CTCGCCCTTG CTCACCATGG TGGCGACCGG TGGATCCCGG GCCCGCGGTA CCGTCGACTC
     3421 TAGCGGTACC CCGATTGTTT AGCTTGTTCA GCTGCGCTTG TTTATTTGCT TAGCTTTCGC
     3481 TTAGCGACGT GTTCACTTTG CTTGTTTGAA TTGAATTGTC GCTCCGTAGA CGAAGCGCCT
     3541 CTATTTATAC TCCGGCGGTC GAGGGTTCGA AATCGATAAG CTTGGATCCT AATTGAATTA
     3601 GCTCTAATTG AATTAGTCTC TAATTGAATT AGATCCCCGA TGGAAGAGAT GGCGAAGGTT
     3661 CTCATCCACA AGTTGTTGGT GATGCTAACA TTGCTTCCGA GCAATGTATT GCTGCCGAGG
     3721 GTGTCGATTA GTACAGCAAA GCGCGAGAAA CAAGTGGCTG CTGTGGCCGG CGGTACCCAT
     3781 AAACCAGTTG AGATAATTTA GACAATAAAT TGTAATAGGC AATTTGATGA CGTCCCGCAT
     3841 GTACGGATCG TCAACGCTGC ACGCTCGGGA ATAGGGTGAG AAGTGCTGCC CGTTTGTTGG
     3901 ACAGCGCAGG TGGTGATCGT GTTGTTTTGG TATCACGAAA ACAAGATAGA ATCTGAGAAA
     3961 AAAGAACCTT TTTATCACTT TACCCGCACC GATACTGGTC TTTCTTAGAT CATTTCGGGG
     4021 CGAAGCAAAC AAGTCAGCAC GGCATACAAA TTTTGGGATA ATCCCTTAGT CAGTGCCTGT
     4081 GTCTTTCTAA TGAAGATTGA CAATGGCGGC GGCGGCGGCG GTGGTAGCAA CCAACGCACT
     4141 GGCCTTTGTT TTCATACTTT TCAATGAATT CAATTAGCTT AACCAACGAG GTTGTTGTTG
     4201 ATGTTCTCAC GGAAGTCAGG AGAAGCTCCG ACGGGCGTGT AGGGTAAGTC GCAGTTCGGC
     4261 AGCATTCTGG CAGTGTACAC CCAGCACACC GAGCGTTCTA GAACGCTGAA TCGTTGAAAC
     4321 AGAGCAAATG ACTGAAGGGC TGTGTCTATT GCTGGGCTGT ACAGAATGAC GCATTTCGTG
     4381 TGGCAATGTG TTTTCGAAAG ATTGAAGCAT TCTGATAGAG CCGTGTGTCA GGAATGTAAA
     4441 TATTTAGATA CTGCAGCTAA CTACAAACTG AGAGAGCTTT CAGCTGGCAG GTGTCCCTAC
     4501 TTGGTGGTCT TTTCTAGAAA GAATGTAGAA TTGACATGTT GTGATGATTT ACCAAAAACA
     4561 ATTTGAGAAG CTAGAGAGTG AATCGTTCTA GCCACCGGAA CCATCTGGAA TGTTGAAGAT
     4621 TTTTCTGGAA TATTAGTAGA CAAAAAGAAG GAGGTCTCAA CTCCGAGTAA ATTTGCAAAA
     4681 TAGTTTTGAC TAAGTACAAA AAGGAAAGTT CCAAGATGTC ATACTCTACT AATGCTTCAA
     4741 CTCACACATG ATCGCACACG ATTCTAGAAA CTCGTGGATG CATTAAAGCC AATGCCTTTT
     4801 CACCGGTGAA TGACTGATAA GCAGATGCCT AGTTGGCAGT CTTGGTAGGC CACCATTCTC
     4861 GATCAATGTC AAGTTCAGGA GCAAGGCTTC AAAGCCGACG TACAATGTGT GCAGCGGGCA
     4921 TGGGCGGTTG TAGTGTCCGC GGTGCCGCGT TTAGCATGCG GCTTTGCCAC TTCCCTTATC
     4981 AACAACACTG AGTGTGCACT TTCACGATAA TCGAAGCTGT CGTTTGTAGA TGCCCTCCCG
     5041 ACACCCCGGG GGTCACGTGG GTCAGTACGA AAAATAAAAC TGATAGTAAG CAGCCGCAAG
     5101 TGCTCCGTCA CTCGCTATCA GCCGGAATAG GTCAACAGAC TGCCAGCAGA TTATGCAACA
     5161 TGTGGACGGT TACGCTCGTA CTAATCATTT GCTTCTATGC TTGCTTCTTT TCTACGCCTA
     5221 CCCCCTCGGA CAACAGCATC GTGCAAGTCG GGATGCCCAG CACGTAAGTA TTTAGCACGC
     5281 CGCGGACGAT ACCGATCTAT GGCTGACTGG GTTACGGGCA ATTACTAACG GCAATGTTTC
     5341 TCCTGCTTCC TGCTCCTGCT CACTACAGCT AACAAATCCA GCAAGTACAC GTTCAAGCCC
     5401 GGTTTCGTAT ATCAGTATGA TGTGGACAGC TATGTGCAGC TTCAGCAGTC GGACAAGGAG
     5461 AACAAGCAGA CCACGCTGAA GGTCGACGGC AAGGTGGAGG TGTACGCCGG CGACAACTGC
     5521 CAGTACACGC TGAAGGTGGT CTCGCTGACC TCGTACGCAC CGGACGGCAA GAAGACGGCG
     5581 TTCGGGGCCG ACATCAGCAA GCCGGTCCAG TTCACGCTGT CGAACGATGA GCTGCTGCCC
     5641 GAGATCTGTA CCGAAGCGGA CGATACGGAC TTTTCGCTGA ACGTGAAGCG TGGTTTGATT
     5701 TCGCTGTTCC AGGTCGCGCA GGAGAAGAGC ACCGAGACGG ACGTGTTCGG CGTGTGCCAG
     5761 ACGTCCTTCT CGAGCTACCC GTCCGGCGAT GCGACCGTCG TGGAGAAGGT GCGCGACCTG
     5821 GGCAACTGTG CCTACCGCGA ATCGCTCTCG AACAGCTTCG TGACCCGCAT CGTCAACAGC
     5881 AAGGCCGGCA TCAAGTCGAC GCCGCTGCTG CAAAGCTCGT ACAATGCCCA GCAAACGATC
     5941 AAGGGTGGTC TGCTGTCGGC GGTCAAGCTG TCGGAGGAGT ACCAGTATCT GCCCTACCTG
     6001 AAGGACAAGG TTGGCGTGAC CGCGAAGGTA ACGACGAAGC TGACGCTCAC CGGCAACAAG
     6061 GCGGGTGCTG CGCCGGCCCT CGGCGCTGCG AGTGAGCCGC GCACGATCAT CTTCGAAAAT
     6121 CCCGACCAGC AGCCGGCCGG CAATCTGGCC GTCATCAAGC AGGAGCTGAA ATCGACCGTC
     6181 GAATCGTACA CGAACGGCAA CGTGGGCAAG AAGACGGCCA ACCTGTTCGT GGAGCTGATC
     6241 CAGCTGATGC GCTACTCGAA GAAGGAGGAC CTGCTGACGC TGTACAACCA GGTGAAGGCG
     6301 GGCAGCGTAC ACTCGAACAA GTCGCTCGCG CGCAAGGTGT ACCTGGACGC GCTGTTCCGC
     6361 GTGGGCACGG GCGATGCGGT CGAAGCGATC ACGCAGCTGT ACAAGAACAA GGAGCTGACC
     6421 GGTGCGCAGG AGCAGAAGCT CGCCTTCGTT TCGCTCACGC TGGTACAGTC GATGACGCAG
     6481 GATGCGCTGA AGGCGGTCAA CAAGCTGCTG GACGGCAATC CGCCCCGCGA GGCGTACCTG
     6541 AGCGTGGGTT CGCTGGTGAG CAAGTACTGC CAGAAGCACG GCTGCCAGTC GTCGGATGTG
     6601 AAGGAAATCT CGAACAAGTT CGGCGCCAAG CTGGGCAAAT GCCAGTCGAC GTCGCGCGCC
     6661 CAGGAGGACG TGATCGTGGC GGTGCTGAAG GGTGTCCGCA ACTCGGACAA TCTGGTCGCC
     6721 CCGCTGCTCG ATAAGGTGAT CCAGTGCGCC GGTCCCGATG CTTCGTCGCG TGTGCGCGTG
     6781 GCTGCCCTGC AAGCCTACCC GGCCGCTTCC TGCAACAAGA AGATCGTCAA CGCGGCGCTC
     6841 AGCACGCTGA AGGACACGAA CGAAGACTCG GAAATTCGCA TCCACGCGTA CCTGTCGCTG
     6901 GTCGAGTGCC CGTCGGCGAA CGTTGCCAAC GAGCTGAAGG CACTGCTGGA CGCGGAGAAG
     6961 GTTTACCAGG TCGGTTCGTT CATCACGTCC CATCTGGCCA GCCTGCGGGC GTCGGTCGAT
     7021 CCGACGCGGG ACGCTGCCCG GCAGCACTTT GGCAAGATCC GCACCTCCAA CAAGTTCCCG
     7081 TTCGATGTGC GCCGCTACTC GTTCAACCGG GAGTTTTCGT ACGCGGTCGA GTCGCTCGGT
     7141 GTCGGCGCCA GTGCCGAGAC GAGCGTGATC TACTCGCAGA AGAGCTTCCT GCCGAAGTCG
     7201 GTTGGACTGA ACTTTACGGC CGAGCTGTTT GGCAATGGGC TGAACGTGTT CGAGCTCGAG
     7261 GGACGCCAGG ACAATCTGGA ACGGCTGGTG GAGCATTACT TCGGACCGAA GGGCTTCTTC
     7321 TCCGGCATGG ACATGCAGGC GGCGTACGAT CTCCTGGCCG AACAGTACCA GAAGCTGTCC
     7381 GGCAAGGCGA AGGAGCGCTT CCGCCGCGGC ATCCGCGAGG ACATCCGTGC GCTGGCCCGC
     7441 ACCGTCGATC TGCACAATGA CGCGCTGAAG GACTTCAACC TGGACGTTAC GATGAAGGTG
     7501 TTCGGCTCGG AGCTGTTCTT CCTGAGCACG GGCGAGAACG TGCCGACCGA TCCGGAGCAG
     7561 TTCCTGGACA AAGCGCTTGA ATGCTTCGAC AAGATGATCG AAGGGGCCAA GAAGTTCGAG
     7621 CACACGTTCG AGCACCATGC CCTGTTCCTG GACACGGATC TGGTGTACAC TACCGCTCTC
     7681 GGTCTGCCAC TGAAGCTTTC GGCGCAGGGT GCCGGTGTGG CGCGTGTCGA TGCCGCTCTC
     7741 GAGCTGGACG TCAAGAGCCT GGTGAAGGAC TACAGCAACG CCAAGTTCAA CACGAAGTTC
     7801 CAGCCAAGCG GCAGCTTCGA GGTGACCGGA ACGATGAGCG TGGACGCGTT CAACGTGATG
     7861 ACGGGCATGC AGGTGGCCGT GTCGGGCCAC TCTTCGGGCG GTGCGGCCGT CAAGTTTGCG
     7921 CTGCACGATG CCACCGCCTA CGATCTGACG GTCGACGCGC TCCAGGGCAA GCAGGAGCTC
     7981 GTGTCGGTCA ACTTCCGCGA GGTGATCATT ACGCGCGAAC GGGGCAACCA GCTGATTACG
     8041 CTGCCCGCCA AACGCCAGGA CTTTGGCTCG AAGTTCAGCG AGTGCTTCGA CAATCTGTAC
     8101 AGCGTGATCG GTGTGACGGT GTGCGCCGCA CACAAGGTCG ACCAGGAGGA GCTGTTCGAG
     8161 GTGTACCTGG AGGTGGAGCC GAAGTTCCAC TTCTCGGGCA AGTTCGACAA CTCGAACCCG
     8221 CAGCACCTGC AGCTGTCCCT CAGCTTCGAT ACCCCCGGGT CGCAGACGAA GCGTCTGACG
     8281 AACCTGCGAC TGGAGGGTGT GACCGCCGGC GAGCTGTACG TGAAGGCGAC GCTCGAGTCA
     8341 CCGATCCGCA ACGTCGACTT CAAGCTGGGA GTAAACAACA ACGACAAGGA GGTGGCCCTG
     8401 TACGCCGTTG CGCACAATGG CGTGGAGGAG TACCTGGCCA AGATTGGCTT CCAGAAGGGT
     8461 GCGTCCGGTG GGCGCGATGA GTACGTGCCG ATCTTCACCA TCCGCTCGCC GAACGGCGAC
     8521 GCCCAGGTGT CCAAGTTTGT GCAGACGACC GGCAAGATCG TGGTGGAGAG CCTGGATGGC
     8581 GGCAAGCGCA AGTACAACCT GGAGAACATC GAGTGGACCA GCCCGTACGC CCCGAAGACG
     8641 ACCATCAACG GGCACGTCGT GTCGAACGGC GAGCGTAGCT TCGATGCCAA CGTGGACGTG
     8701 AACGTCGGTG ACGTGAAGAA CAACGTGGTG GGCCATCTGG ACTTTGATCT GAAGCACGTC
     8761 AAGCTCGACC TGGAGAAGAA GACGCCGAGC GATGCGAACA AAAACTTCAA GGTCAACCTG
     8821 GAGGTGGCGT ACACGGACAA CTCGTTCAAG AACCTGTTCT CGTTCGCCAG CGGCAAGGAC
     8881 TTCAACAATC CCTCGAACAA GTACGAGCTG AGCCAGTACG CTGAGTTTGA GCTCAAGCCC
     8941 GAGGCGCAGG GACTGGAGTC GCTGAAGCTG GAGAACAAGC TGCAGCTGCC GAAGCAGCTG
     9001 ATCCGACTGG ACTTTGCCAC GAACAAGAAC AAGTTCTACC TGGACGGTGA GTACGGGTAC
     9061 GACAAGTACA AGATTGCGGC GAACGTGGAT GCGAAGTACA ACGAGAAGAC GCCCGGGGAT
     9121 TACGATGTGC AGCTGGGCGG TTCGCTCAAC AAGCACTTCT TCAAGTTCTT CTCCAAGCGC
     9181 ATCGTGGAGG CCAACAAGAG CCGCTTCAGC AACAAGCTGA CGGCCAGCAC CGGTACCAAG
     9241 TTCGAGCTGA ACGGTGCCGT CACGAACCGC TTCACCAGCC AGGATGGTGA GCTTAACCTC
     9301 GAGGGATCTC TCATCGCTGT CGAGAAGGCT TCTCCTTACA AGTGAGTAGG AAACGCAATG
     9361 ATTATTGGCA GGGCAATGAT ATCACTAATT GGTTGTGCTG TTCAGGCTTT CTTTGACAGT
     9421 TCAGTTCTCG GCCGCGAACG TCCTCTCCAA CGCCAAGGTG CTGGTGGATA AGGAAGAGTT
     9481 CGCTACGTAC GACTTCAAGC TAGAGCGGGG ACAGGATCCG AACGGCAAAT TCACGGTAAG
     9541 TCTCCGCAAA CAACTCTCAA CTCGTTTCAG TTTTCCTTAC GCCACATGTT TCATTGACTC
     9601 TCCCAACAGT TTGTGGTGAA GGACTTCTTC AACGGAAATG GTGAACTAAA GTCGGCCAAG
     9661 GGTCAGGGTG AGCTGTTCGC CCTGGTCACG TTCGTCAAGC AGGACCGCAA GGTGAAGCTG
     9721 GACAGCAAGT TCAAGGTGAA CGCACCGGTG TACGATGTGG CTGCCGATTT CTACTACGAC
     9781 TTTGAGAAGG ACAACAGCAA GAAGGTACGC TTCGAAACCA AGAACAAGGT CACCCAGACC
     9841 AGCTTCGACA GCAAGAACAA GGTGGAGGTG TTCTCGGAGA AGTACGAGCT GAACGTACAG
     9901 AGCCAGGGCA ACCCGAAACC GGTCGATGGC AAGTTCAACG TCAAGGTCAG CCTGCTGCTC
     9961 CCGACCGGAC GCCAGTTCGG CGGCGAGTTC CAGCGCGATG CCTCGACCAA GGACGAGAAG
    10021 CGCTCCGGCA AGATGGCGGC CAGCGTGTAC GACAAGCAGC CCGGTGGCAA GAAGCGCTCG
    10081 GTCGAGTGGG CCGGCGAGCT GAAGGATATG GACGTGAAGA CGAAGTTCTT CGACGCGGTG
    10141 CACAATGTCA AGTACTCCGA TCTCGAAGGC AAGGATGTGG TGCTGGACGT GACGCTAAAG
    10201 CATGCGCCGG CTGGTAGCTA CAAGTCGGCC GCCGGTAGCT TGAAGGTTTC GGGCAGCTTG
    10261 CTTCCCCAGG TGACGGAACT GAGCGTGGTC GTCGATGAGT ACTGTGAGCA TCACGCCAAG
    10321 TACCACGTCA ACGGCAAGTA CGGGGCGGAC TTTACGGCTG CACTGGTCGG TGGGTACCAC
    10381 ACCGGAGGGC ACGGCAAGCC GGCCACCCAC GACCTGAAGG TTGACGTGGG TGCTCCATCG
    10441 TACAAGGTGG GCGTATCGTC CAGCGGCAAG TACCTGCAGC CAGAGTCGGA CGACGGTGTG
    10501 TACGAGCTCG ACTACAGCGG ATCGGTCGAC TTCAACGGCA AGAGTGCGTC GGTGTCGACG
    10561 CAGGCCAAGG GCAACTACAA CCGTGGCAAT GGAAAGCTGA ACCTGAACCT CCCGAACGTG
    10621 GACCCGATCG CGGCCGAAGG TTCGTACACG TACGACGCGA AGGAGGAGGG ACCGTTCCAG
    10681 ACGAACGGCG CCCTGAAGGT GTCGTACGGT GCGGGCAAGA ACTTTGAGTT CACCGGCACC
    10741 GCCAAGGCCC CGTCGATGGA CGATATCCAG GTGCACGCGA CGCTCAAGTC GGAGTTCGAG
    10801 AACGTGCGCA GCGTCGACCT GACCTTCAAG CACGCCAAGT CGAGCGACAG TGCGTACAAC
    10861 ACGAAGCTGC AGCTGACCGC CGACGACAAG AAGTTCAGCG TGGAGAACGC GGTGGTCGTG
    10921 TCGGAGACGA ACCCGTCCGT TGACTTTACG CTCGGCTATC CGGGCAAGAC GGTGAAGGTG
    10981 TCGGGCAGCT ACAAGGCGCT CGGCCACAGC GCATTCAAGG CGGACGCCAA GGTGCAGAAC
    11041 CTGGCCAACT TCGACATGGA AGCGAACGTG GAGGCCAACT TTGACAGCTA CCAGACGTTC
    11101 TACGTGAAGC TGTACGGCGA TGCGCCGATG CTGAACACGA ACAAGTTCTC GGTGGAGGTG
    11161 AACGCGAAGC CGGGCTCGAA CGGCAAGGGC GTTAACTTCC GCGCGTCCGA GGGCGGTAAG
    11221 GACATACTGA GCGGCTTTGC CGACTACTCG GTGAAGGAGC AGGGCAAGGC GATGGTCATC
    11281 GAGGGCCAGG GCAACGTGAA GCTGTACGAC AAGCAGCAGA CCGCCACCTT CAAGCTGATC
    11341 CGCGAGAAGC TGTCCGAGTC GGGCATGTCG GCCACGCTGA CGGCCTCGGT GGGCAAGTTC
    11401 ACCGTGCTGC ACGAGTCCCG CGTCCAGCCG AACGATTTCC GCGTCAAGAC GAGCGTGTGC
    11461 GACGAGAAGA AGAAGTGCAC CAAGCTGGAG CTGCTCTCGA AGCTGGAGCG CGCGGCCGGT
    11521 GCGTTCAAGC ACGAGGCGCT GGTGTCGGTG GAGGTGCAGC AGATGGGCTA CGAGCACGAG
    11581 TTCGGCCTGT CGGCCAAGAC GAGCGCCAAC GGGCTGAAGT TCGACCACAC CACGGACGTG
    11641 CAGCTGAAGG AGAAGAACCA GCCGAAGTAC CAGTACCTGT TCTACGTGCA TCCGACCTCG
    11701 GCCGGAGCGA GCCTAATACT GCCGACGCGC ACGGTCGCCG TGGAGGGTGT GCTGAACCTG
    11761 CCCAAGGACA AGTTCGGACC GTTCGACGGC AGCGTGTCGT TCTATCTGGA CAAGAAGAAC
    11821 GAGCCGGACA ACAAGGCCAC GTTTGCGGTG CGCGGCGAAA CGAAGATGCT CGGCTCGACC
    11881 GGCGTCAGTG CGAACGGTGC GCTGACGTTC TCGCATCCGA CGATCAAGGC GCTGGCCATC
    11941 CGGGCCAAGG GTACGCTCGA TGGGGACAAG CAGACGGCGG ACGGATCGGT TGAGTTTGAC
    12001 GTGTTCAAGA AGGCCGGCGA CAAGATCGTT GCCAAGGTGC GCTACGCCAA CAGCGACCAG
    12061 TCGTTCAAGG GCTTCAACAT CACGACCGAG GCAAGCGTGT CAAGCAAGGG CCTGGGGCTG
    12121 AAGTGTGGCT TCAGCGGCCA CTCGGCCATG TCGCTCGCAA CCCGCCAGGC CAGTGCGGCC
    12181 GGTTCGCTCA CGCTGCCGTT CGAGGGCTAC ACGTTTGGCT CGTACTTCTT CGGCAGTCCG
    12241 GAATCGTTCG ACTTCCTGCT GACCCGCTTC GGGGACGACT TTGTGCGCGC GCACGGCACG
    12301 TACGATGCGA AGAAGTACCG CGGCGACCTC ACCTCCACCT TCAAGTTCCT GCCCAACCGG
    12361 CCGGTGGTGT TCGAGTCGCA GCTGAACGGT CTGTCGTCGG CCAAGTTTGC CTTCAAGCAG
    12421 GACCAGTTCT TCAGCGCGGA CGGTACGTTC GGCGTGGACA AGGCGATGGT GCTGAAGGTG
    12481 ATGGGCGAGG GTAAGCCACT GCTGAACGCG AAGGTAACGC TGGACGCCAG CCACTTCCTA
    12541 TCGACCGAGT ACAATGTGGA CGAGGCGAAC GCGAAGGCGT TCCTGGTGTC GCTCAAGAAC
    12601 CAGCTGAACG CCGACTTCGA GGTGACGCGG GCGGACGTCA GCCAACGCTA CGCCAAGCTG
    12661 GCCGAGGAGC TGAACAAGCT GTCGACGAAC CTGGTGAGCG CGCTGCCCGA GTTCGGCAAG
    12721 TTCCAGGAAA GCTACGCCAA GCAGCTGCAG AAGCTGCAGG AGGACATCAT GAGCGATCCG
    12781 GCCCTGGCTG AGTTCGTGAA GGCTGCCACC AAGATCTTCC AGCAGGTGAG CGAAGTGTTC
    12841 GGCCAGCTGT CGCAGGTGTA CGTGGAAAGC TTCCGCAAGA TGTCCGCCCT GGTGAACGAC
    12901 GTCGTCGCCC AGCTGATGGA AACGTTCAAC ACGAAGGTAC TGCCGGCGCT GAAGGAGCTG
    12961 TCCACCAAGG TGGAGGCGAT CTTCTTCAAC GTGTACGAGG AGACGGTCAA GCTGGTGGTG
    13021 GCCGTGTTCG AGCGCACCGT CAAGGCGCTG AAGGTGTTCG AGGAGGACTT CAACAAGATC
    13081 GCCACGAGCG TGTCGGAGCT GTTCCGCACG TTCGCGCAAA CGTTCAGCAA GGCCGTCCAG
    13141 GTGCTGGAGA AGGAGCTGAA GGAGCTGTAC AAGCTGGTGC AGGAGTACTT CGACACGTTC
    13201 GACGAGTTCA AGGCGGTGAA GGAAACGTTC AAGGAGTACT TCGACGGGTT CGACCGGTAC
    13261 GCGTACCAGC TGCTGAAGGA GCTGCTGTCG CTGGTCGAGG CGGTCTACCC GATGCCGGAG
    13321 GTGACCGAGC TGACCACCGC GATCAACAAG TACATCACCA GCAAGCTGGA CAACAAGCCG
    13381 GTCAACGATG TGGAGGAGCT GAAGACGCTG TTCGTGAGCC TCGTCAAGGT GCTGAACCAG
    13441 GCGGTGGAGC GCCTCATCGC GGGAGTGAAC GTGCAGCTGT CCGAGCCGAC GTTCGGCAGC
    13501 GACAGCTTCA CCTCGTTCGT CACCTTCAAG TTCCTGCCGT ACGTGTCGAG CATCCAGTTC
    13561 AGCCCGTGGA ACTTTGTGCG CAACGAGAAG TTCTACTCGG TGCGCGACCT GATCCACCAG
    13621 CTGCGCCTGT ACGCGTTCAA TCCGTTTGCG CGCGTGCCGA TGTTCCACAT GCACGCCCAG
    13681 CTCGGCGACG GTGGCCACTT CTTCACGTTC GACGACAAGC ACTTTACGTT CGCGGGCAGC
    13741 TGCTCGTACC TGCTCGCGTC CGACCTGGTC GACGGTAACT TCAGCATCGT GGCGGACATG
    13801 GACGGCGGAC GCCTGAAGTC GGTGACGCTC GTGGACAAGG ACTCGACGGT CGAGCTGACC
    13861 GCCAAGGCGG TGGTGAAGTA CAACGGCAAG GAGACGGATC TGCCCATCCA CCAGAAGGAT
    13921 GTGTACGTGT TCCGCAAGTA CTACACGGTC ACGGTCGGCA CCAAGTACGG TGCGCAGGTG
    13981 ATGTGTACGA CCGACCTGAA GATCTGTCAC TTCTTCGTGT CCGGGTTCTA CTTCGGGCGG
    14041 CTGCGCGGTC TGCTCGGCAA CGGCAATTAC GAGCCGTACG ACGATCTGGC GGTGCCGAAC
    14101 GGCAAGATCA CCGAGGTGTC GACGGACTTT GCGAACAGCT ACAAGACGAG CCAGGCGTGT
    14161 GCGGCAGTCG CCGACCACGG TCACGATTCG CACGACCACT CCAGCCCGAC CTGTGCCAAG
    14221 TTCTTCGGCT CGGAGTCGTC CCTGAAGCTG TGCTCGTACC TGCGCGACCA GACCGGCTAC
    14281 AAGGAGGCGT GCAACCATGC GGCGCACGAT GCGGGCGAGA AGGCGGACGA GGCCGCTTGC
    14341 GGCATTGCCC GGCTGTACGT GTCCGCCTGC ACGCTGTACG GCATCCCGAC CGTGCTGCCC
    14401 AGCCAGTGCG AGAAGTGCTC GACCGACGAG GGCCGCTCGG TCGATCTGGG CGACTGGTAC
    14461 TCGGTGAAGG CGCCGCAGAA GAAGGCCGAC GTGGTGGTGG TGGTCGACAC GTCGCTCGGC
    14521 ACGCTGCTCG GCGAGCTGGT GCAGTCCACG ATCAACGATC TGCGCAAGGA GCTGAAGGCG
    14581 ACCGGCATCA GCGACGTGAA CGTTGCGGTG ATCGGCTACA GCAAGACGGA CAAGTACACG
    14641 AGCCTGTTCT CGAACGGCGG CAAACTGGAC TACACGGGCA AGCTCGGCCA GGCCGACGTC
    14701 AGCAGCGGAC CGAAGCAGTG CCGCGGGCTG GTGACGGGCA TCGAGTCGGT GGACGCGTTC
    14761 CTGCAGCTGC TGCAGAAGAT GGGCGAACAG TCGCGCGAGA ACTTGGGCGC TACGACCGAG
    14821 GTGTACGCGC TGCGCCGTGC CTTCAGCTAT CCGTTCCGTG CCTCCGCCAG CAAGTCGGTG
    14881 CTGGTCTTCC GTTCCGATTC GTTCGAAATC GCTCAACCG
//


Supplemental File 1C: Lp gene drive plasmid


LOCUS       Lp-GD               13956 bp ds-DNA     circular
DEFINITION  synthetic circular DNA
ACCESSION   .
VERSION     .
KEYWORDS    .
SOURCE      synthetic DNA construct
  ORGANISM  synthetic DNA construct
REFERENCE   1  (bases 1 to 13956)
  AUTHORS   Emily GREEN
  TITLE     Direct Submission
  JOURNAL   Exported Wednesday, Sep 18, 2019 from SnapGene 4.3.11
            https://www.snapgene.com
FEATURES             Location/Qualifiers
     terminator      268..295
                     /label=rrnB T2 terminator
                     /note="transcription terminator T2 from the E. coli rrnB
                     gene"
     terminator      387..473
                     /dnas_title="Escherichia coli rrnB"
                     /gene="Escherichia coli rrnB"
                     /label=rrnB T1 terminator
                     /note="transcription terminator T1 from the E. coli rrnB
                     gene"
     primer_bind     537..553
                     /label=M13 fwd
                     /note="common sequencing primer, one of multiple similar
                     variants"
     misc_feature    771..773
                     /label=lac promoter
                     /note="lac promoter"
     misc_feature    774..2326
                     /label=Lp promoter
                     /note="Lp promoter"
     misc_feature    2341..2665
                     /label=U6 promoter (ag)
                     /note="U6 promoter (ag)"
     misc_feature    2671..2740
                     /label=tRNA-gly
                     /note="tRNA-gly (missing terminal A, which is added when
                     cloning guide linker)"
     misc_feature    2741..2848
                     /label=gRNA template
                     /note="gRNA template"
     misc_feature    2854..2923
                     /label=tRNA-gly
                     /note="tRNA-gly (missing terminal A, which is added when
                     cloning guide linker)"
     misc_feature    2924..3031
                     /label=gRNA template
                     /note="gRNA template"
     misc_feature    3037..3106
                     /label=tRNA-gly
                     /note="tRNA-gly (missing terminal A, which is added when
                     cloning guide linker)"
     misc_feature    3107..3214
                     /label=gRNA template
                     /note="gRNA template"
     misc_feature    3220..3289
                     /label=tRNA-gly
                     /note="tRNA-gly (missing terminal A, which is added when
                     cloning guide linker)"
     misc_feature    3290..3397
                     /label=gRNA template
                     /note="gRNA template %22misc_feature    3403..3473
                     /label=tRNA-gly /note=%22tRNA-gly"
     misc_feature    4556..4624
                     /label=3xFLAG
                     /note="3xFLAG"
     misc_feature    4625..4675
                     /label=NLS
                     /note="NLS"
     misc_feature    4676..8776
                     /label=Cas9
                     /note="Cas9"
     misc_feature    9870..9916
                     /label=3x Pax6 binding sites
                     /note="3x Pax6 binding sites"
                     /label=TATA
                     /note="TATA"
     CDS             10115..10848
                     /label=DsRed2nls
     polyA_signal    10856..11093
                     /label=SV40
     promoter        complement(12198..12216)
                     /label=T7 promoter
                     /note="promoter for bacteriophage T7 RNA polymerase"
     primer_bind     complement(12221..12237)
                     /label=M13 rev
                     /note="common sequencing primer, one of multiple similar
                     variants"
     CDS             12350..13159
                     /dnas_title="aph(3')-Ia"
                     /codon_start=1
                     /gene="aph(3')-Ia"
                     /product="aminoglycoside phosphotransferase"
                     /label=KanR
                     /note="confers resistance to kanamycin in bacteria or
                     G418 (Geneticin(R)) in eukaryotes"
     rep_origin      13306..13894
                     /direction=RIGHT
                     /label=ori
                     /note="high-copy-number ColE1/pMB1/pBR322/pUC origin of
                     replication"
     misc_feature    complement(11094..12189)
                     /note="3%82%C4%F4flk homology arm"
     misc_feature    complement(8828..9869)
                     /note="zpg terminator"
     misc_feature    complement(3481..4554)
                     /note="zpg promoter"
     misc_feature    11145..11164
                     /note="gRNA4target site present by mistake%E2%80%9C
                     misc_feature    2742..2761 /note=%22gRNA1"
     misc_feature    2925..2944
                     /note="gRNA2"
     misc_feature    3108..3127
                     /note="gRNA3"
     misc_feature    3291..3310
                     /note="gRNA4"
     misc_feature    2742..2761
                     /note="gRNA1"
     source          1..13956
                     /dnas_title="pENTR-Lp5'-sgRNAx4(lp)
                     -ZpgCas9Zpgt-DsRed-Lp5'"
ORIGIN
        1 ctttcctgcg ttatcccctg attctgtgga taaccgtatt accgcctttg agtgagctga
       61 taccgctcgc cgcagccgaa cgaccgagcg cagcgagtca gtgagcgagg aagcggaaga
      121 gcgcccaata cgcaaaccgc ctctccccgc gcgttggccg attcattaat gcagctggca
      181 cgacaggttt cccgactgga aagcgggcag tgagcgcaac gcaattaata cgcgtaccgc
      241 tagccaggaa gagtttgtag aaacgcaaaa aggccatccg tcaggatggc cttctgctta
      301 gtttgatgcc tggcagttta tggcgggcgt cctgcccgcc accctccggg ccgttgcttc
      361 acaacgttca aatccgctcc cggcggattt gtcctactca ggagagcgtt caccgacaaa
      421 caacagataa aacgaaaggc ccagtcttcc gactgagcct ttcgttttat ttgatgcctg
      481 gcagttccct actctcgcgt taacgctagc atggatgttt tcccagtcac gacgttgtaa
      541 aacgacggcc agtcttaagc tcgggcccct acaggtcact aataccatct aagtagttga
      601 ttcatagtga ctggatatgt tgtgttttac agtattatgt agtctgtttt ttatgcaaaa
      661 tctaatttaa tatattgata tttatatcat tttacgtttc tcgttcaact tttctataca
      721 aagttggtac cgggcccccc gctagcgtcg acggtatcga taagcttgat cggatccaac
      781 agttcattcc cgattgaggg attttattcc ccgggggcct tttcaaacgg cttaatataa
      841 gcaattaata gtattttttc tttcaggtta gtttactgta atggtgtaat tgtcatctta
      901 cacctccgtc tgataagaga ttacgaagct cagtatgatg aaataaataa gataaattta
      961 tttaaaaaag aacaattgct atgagagtga aatacaacag tggcgttcac aatattcgaa
     1021 aaacaataaa attaaaaaaa aaaaacaaga aaacattcac aaacatatca atctgctttc
     1081 atcgacacca aactgctagc ctccccagtc taaccgcggt ggggacgttt aattgccttt
     1141 gttctcgcac ccggtcaaac atacacttcg gaccttgctc cgaaccccac tgtaatccct
     1201 agctcgtcat catcattgcc ggcatcatgc taagcgtgca ttattttcac aacttagcgt
     1261 aatgctagcg tgcgctagca acaaactcgc ccgcagactc gtcacagcac cggtacgatc
     1321 gatcgtttac cgttcccttt cccgatcggg ttggctgcga tatccgtgtc cgggtagaaa
     1381 acttcccctt ttacacatac acatacactc acatacacac agagctgaat agcaacttac
     1441 cttatctgtt cgtcatcgct cggccggatc tggatgaatc ttcgcaccga taaccatgtg
     1501 gatctacgac ctccgcttgg ctgtctctct gctcatgtgt atgtctgtgt gtgtgtgtgt
     1561 gtgtgtgtgt gtgtgtgtgt atgtgagctt cttccctcaa atccctcgat ctcgctgtgc
     1621 caacaatcaa acgtgcaagt gcaaacattg caccccattg attatacacc aacaccaacc
     1681 aattcccctt gcggaggcat cactgtgctc ggcagcatgt ttaccgcaga tctacagaga
     1741 acttcaattg aggtcctttc cacccccagc cctcaaccgg caatccggca gccactggat
     1801 catcagcgaa agagagagag agcagagcag aacagaggtg accaactgtg gtatcgcttc
     1861 ccgcgcgccg gtgtgttggt gtccatttcg gtgatcgcga tcccggccgc ttccagcacc
     1921 gtccaccgat cagtcaccaa aacgctctcc aaacccctta tcagcaccgt tcgctgatgt
     1981 gaacccccgt tcaaacccca aatgcagtgt ttgtattgct gtgtgtatgt gtgcgtgcgt
     2041 gtgtgtggaa attctataaa agtaggcaac cgtggccggg atccgttatt cccgtcctga
     2101 ggcccgcccg ggatcgctgg tgacgacaga cgagccgctg tgtgacgtac gtagtgcccg
     2161 atcggtaaag agtgaaccgt cttctctgca gtgttggaga gaacggtttc atctttttcg
     2221 cccacacccc ccccggttac attccatgtt gagctgtaag gtctagtgaa catttcctga
     2281 gtgtggaaag tgtggtttag tgcgtgagag tgcacggaca cgacacggaa cgatgtgggt
     2341 ttatccacat atacactgaa gcggtttggc tcttggttat tctagaaaac tctgtttttg
     2401 aattaatcat aggaaaactg tttattcttt tttgatattt caacgtatta caatgaatag
     2461 taacttttga ttaaaaaaga gatgaagcaa tacaagaaat gttgaaaagt ttatgaaaca
     2521 tcacattagc gtgagttacg gcaggatcaa aaccttataa cagtcacact caggcaaaaa
     2581 atccttcttg aatatccttt atgcatcgct agagcaagga ttgaaagcgc aaagtatata
     2641 tacaaccttt tttcccctcg tccttgtaaa gcatcggtgg ttcagtggta gaatgctcgc
     2701 ctgccacgcg ggcggcccgg gttcgattcc cggccgatgc acacggaacg atgtgggtcc
     2761 tgtttcagag ctatgctgga aacagcatag caagttgaaa taaggctagt ccgttatcaa
     2821 cttgaaaaag tggcaccgag tcggtgctag agagcatcgg tggttcagtg gtagaatgct
     2881 cgcctgccac gcgggcggcc cgggttcgat tcccggccga tgcagaggct gctctggagc
     2941 ttccgtttca gagctatgct ggaaacagca tagcaagttg aaataaggct agtccgttat
     3001 caacttgaaa aagtggcacc gagtcggtgc tccatagcat cggtggttca gtggtagaat
     3061 gctcgcctgc cacgcgggcg gcccgggttc gattcccggc cgatgcactg attcaaagtg
     3121 tgtccgcgtt tcagagctat gctggaaaca gcatagcaag ttgaaataag gctagtccgt
     3181 tatcaacttg aaaaagtggc accgagtcgg tgctggaaag catcggtggt tcagtggtag
     3241 aatgctcgcc tgccacgcgg gcggcccggg ttcgattccc ggccgatgca gcgtgaaata
     3301 tcgtcaggga gtttcagagc tatgctggaa acagcatagc aagttgaaat aaggctagtc
     3361 cgttatcaac ttgaaaaagt ggcaccgagt cggtgctacg aagcatcggt ggttcagtgg
     3421 tagaatgctc gcctgccacg cgggcggccc gggttcgatt cccggccgat gcattttttt
     3481 attcgctggc ggtggggaca gctccggctg tggctgttct tgcgagtcct cttcctgcgg
     3541 cacatccctc tcgtcgacca gttcagtttg ctgagcgtaa gcctgctgct gttcgtcctg
     3601 catcatcggg accatttgta tgggccatcc gccaccacca ccatcaccac cgccgtccat
     3661 ttctaggggc atacccatca gcatctccgc gggcgccatt ggcggtggtg ccaaggtgcc
     3721 attcgtttgt tgctgaaagc aaaagaaagc aaattagtgt tgtttctgct gcacacgata
     3781 gttttcgttt cttgccgcta gacacaaaca acactgcatc tggagggaga aatttgacgc
     3841 ctagctgtat aacttacctc aaagttattg tccatcgtgg tataatggat ctatcgagcc
     3901 cggttacact acacaaagca agattatgcg acaaaatcac agcgaaaact agtaattttc
     3961 atctatcgaa agcggccgag cagagagttg tttggtattg caacttgaca ttctgctgcg
     4021 ggataaaccg cgacgggcta ccatggcgca cctgtcagat ggctgtcaaa tttggcccgg
     4081 tttgcgatat ggagtgggtg aaattatatc ccactcgctg atcgtgaaaa tagacacctg
     4141 aaaacaataa ttgttgtgtt aattttacat tttgaagaac agcacaagtt ttgctgacaa
     4201 tatttaatta cgtttcgtta tcaacggcac ggaaagatta tctcgctgat tatccctctc
     4261 gctctctctg tctatcatgt cctggtcgtt ctcgcgtcac cccggataat cgagagacgc
     4321 catttttaat ttgaactact acaccgacaa gcatgccgtg agctctttca agttcttctg
     4381 tccgaccaaa gaaacagaga ataccacccg gacagtgccc ggagtgatcg atccatagaa
     4441 aatcgcccat catgtgccac tgaggcgaac cggcgtagct tgttccgaat ttccaagtgc
     4501 ttccccgtaa catccgcata taacaaacag cccaacaaca aatacagcat cgagaatgga
     4561 ctataaggac cacgacggag actacaagga tcatgatatt gattacaaag acgatgacga
     4621 taagatggcc ccaaagaaga agcggaaggt cggtatccac ggagtcccag cagccgacaa
     4681 gaagtacagc atcggcctgg acatcggcac caactctgtg ggctgggccg tgatcaccga
     4741 cgagtacaag gtgcccagca agaaattcaa ggtgctgggc aacaccgacc ggcacagcat
     4801 caagaagaac ctgatcggag ccctgctgtt cgacagcggc gaaacagccg aggccacccg
     4861 gctgaagaga accgccagaa gaagatacac cagacggaag aaccggatct gctatctgca
     4921 agagatcttc agcaacgaga tggccaaggt ggacgacagc ttcttccaca gactggaaga
     4981 gtccttcctg gtggaagagg ataagaagca cgagcggcac cccatcttcg gcaacatcgt
     5041 ggacgaggtg gcctaccacg agaagtaccc caccatctac cacctgagaa agaaactggt
     5101 ggacagcacc gacaaggccg acctgcggct gatctatctg gccctggccc acatgatcaa
     5161 gttccggggc cacttcctga tcgagggcga cctgaacccc gacaacagcg acgtggacaa
     5221 gctgttcatc cagctggtgc agacctacaa ccagctgttc gaggaaaacc ccatcaacgc
     5281 cagcggcgtg gacgccaagg ccatcctgtc tgccagactg agcaagagca gacggctgga
     5341 aaatctgatc gcccagctgc ccggcgagaa gaagaatggc ctgttcggaa acctgattgc
     5401 cctgagcctg ggcctgaccc ccaacttcaa gagcaacttc gacctggccg aggatgccaa
     5461 actgcagctg agcaaggaca cctacgacga cgacctggac aacctgctgg cccagatcgg
     5521 cgaccagtac gccgacctgt ttctggccgc caagaacctg tccgacgcca tcctgctgag
     5581 cgacatcctg agagtgaaca ccgagatcac caaggccccc ctgagcgcct ctatgatcaa
     5641 gagatacgac gagcaccacc aggacctgac cctgctgaaa gctctcgtgc ggcagcagct
     5701 gcctgagaag tacaaagaga ttttcttcga ccagagcaag aacggctacg ccggctacat
     5761 tgacggcgga gccagccagg aagagttcta caagttcatc aagcccatcc tggaaaagat
     5821 ggacggcacc gaggaactgc tcgtgaagct gaacagagag gacctgctgc ggaagcagcg
     5881 gaccttcgac aacggcagca tcccccacca gatccacctg ggagagctgc acgccattct
     5941 gcggcggcag gaagattttt acccattcct gaaggacaac cgggaaaaga tcgagaagat
     6001 cctgaccttc cgcatcccct actacgtggg ccctctggcc aggggaaaca gcagattcgc
     6061 ctggatgacc agaaagagcg aggaaaccat caccccctgg aacttcgagg aagtggtgga
     6121 caagggcgct tccgcccaga gcttcatcga gcggatgacc aacttcgata agaacctgcc
     6181 caacgagaag gtgctgccca agcacagcct gctgtacgag tacttcaccg tgtataacga
     6241 gctgaccaaa gtgaaatacg tgaccgaggg aatgagaaag cccgccttcc tgagcggcga
     6301 gcagaaaaag gccatcgtgg acctgctgtt caagaccaac cggaaagtga ccgtgaagca
     6361 gctgaaagag gactacttca agaaaatcga gtgcttcgac tccgtggaaa tctccggcgt
     6421 ggaagatcgg ttcaacgcct ccctgggcac ataccacgat ctgctgaaaa ttatcaagga
     6481 caaggacttc ctggacaatg aggaaaacga ggacattctg gaagatatcg tgctgaccct
     6541 gacactgttt gaggacagag agatgatcga ggaacggctg aaaacctatg cccacctgtt
     6601 cgacgacaaa gtgatgaagc agctgaagcg gcggagatac accggctggg gcaggctgag
     6661 ccggaagctg atcaacggca tccgggacaa gcagtccggc aagacaatcc tggatttcct
     6721 gaagtccgac ggcttcgcca acagaaactt catgcagctg atccacgacg acagcctgac
     6781 ctttaaagag gacatccaga aagcccaggt gtccggccag ggcgatagcc tgcacgagca
     6841 cattgccaat ctggccggca gccccgccat taagaagggc atcctgcaga cagtgaaggt
     6901 ggtggacgag ctcgtgaaag tgatgggccg gcacaagccc gagaacatcg tgatcgaaat
     6961 ggccagagag aaccagacca cccagaaggg acagaagaac agccgcgaga gaatgaagcg
     7021 gatcgaagag ggcatcaaag agctgggcag ccagatcctg aaagaacacc ccgtggaaaa
     7081 cacccagctg cagaacgaga agctgtacct gtactacctg cagaatgggc gggatatgta
     7141 cgtggaccag gaactggaca tcaaccggct gtccgactac gatgtggacc atatcgtgcc
     7201 tcagagcttt ctggccgacg actccatcga caacaaggtg ctgaccagaa gcgacaagaa
     7261 ccggggcaag agcgacaacg tgccctccga agaggtcgtg aagaagatga agaactactg
     7321 gcggcagctg ctgaacgcca agctgattac ccagagaaag ttcgacaatc tgaccaaggc
     7381 cgagagaggc ggcctgagcg aactggataa ggccggcttc atcaagagac agctggtgga
     7441 aacccggcag atcacaaagc acgtggcaca gatcctggac tcccggatga acactaagta
     7501 cgacgagaat gacaagctga tccgggaagt gaaagtgatc accctgaagt ccaagctggt
     7561 gtccgatttc cggaaggatt tccagtttta caaagtgcgc gagatcaaca actaccacca
     7621 cgcccacgac gcctacctga acgccgtcgt gggaaccgcc ctgatcaaaa agtaccctgc
     7681 gctggaaagc gagttcgtgt acggcgacta caaggtgtac gacgtgcgga agatgatcgc
     7741 caagagcgag caggaaatcg gcaaggctac cgccaagtac ttcttctaca gcaacatcat
     7801 gaactttttc aagaccgaga ttaccctggc caacggcgag atccggaagg cgcctctgat
     7861 cgagacaaac ggcgaaaccg gggagatcgt gtgggataag ggccgggatt ttgccaccgt
     7921 gcggaaagtg ctgagcatgc cccaagtgaa tatcgtgaaa aagaccgagg tgcagacagg
     7981 cggcttcagc aaagagtcta tcctgcccaa gaggaacagc gataagctga tcgccagaaa
     8041 gaaggactgg gaccctaaga agtacggcgg cttcgacagc cccaccgtgg cctattctgt
     8101 gctggtggtg gccaaagtgg aaaagggcaa gtccaagaaa ctgaagagtg tgaaagagct
     8161 gctggggatc accatcatgg aaagaagcag cttcgagaag aatcccatcg actttctgga
     8221 agccaagggc tacaaagaag tgaaaaagga cctgatcatc aagctgccta agtactccct
     8281 gttcgagctg gaaaacggcc ggaagagaat gctggcctct gccggcgaac tgcagaaggg
     8341 aaacgaactg gccctgccct ccaaatatgt gaacttcctg tacctggcca gccactatga
     8401 gaagctgaag ggctcccccg aggataatga gcagaaacag ctgtttgtgg aacagcacaa
     8461 gcactacctg gacgagatca tcgagcagat cagcgagttc tccaagagag tgatcctggc
     8521 cgacgctaat ctggacaaag tgctgtccgc ctacaacaag caccgggata agcccatcag
     8581 agagcaggcc gagaatatca tccacctgtt taccctgacc aatctgggag cccctgccgc
     8641 cttcaagtac tttgacacca ccatcgaccg gaagaggtac accagcacca aagaggtgct
     8701 ggacgccacc ctgatccacc agagcatcac cggcctgtac gagacacgga tcgacctgtc
     8761 tcagctggga ggcgacaaaa ggccggcggc cacgaaaaag gccggccagg caaaaagaaa
     8821 aagtaagaat tgaggacggc gagaagtaat catatgtccg cattttgcgc aaaccaggcg
     8881 cttagacaat ttgcgcgtaa gcacattcga aatgtgaaaa gctgaaagca gtggtttcgc
     8941 cagcccgagt tcagcgaaac ggattccttc caagtgtttg cattcctggc ggagtgttcc
     9001 tcccaaaatg cactcaccct gcgtgcagtg ccaaatcgtg agtttcttaa ttttttcata
     9061 ttgtttatta cctaccaact aaagttgttg ttatatattg cgttttacgt acgacaaata
     9121 agttcgtatt cagaaatatt tgcgataaga gagaactcat ttgcgatgaa tctcattgta
     9181 tttagctaag tgccttgata agtaagcgga acagcaggaa tatgacactc cttgggaaat
     9241 acatgtaagc gtctgtgaat tagatatata tacacgcaac caaacggtcc atggttgatt
     9301 taagcactgc ctgttgtcga acattgctat aagcaaaata aagaagcatt cattaatcta
     9361 aaatttcttc aaagtgactt caatgatgat ctctaggcta tagtgaaagc tgaaagctta
     9421 ttcgacaatg caagggaaag tgacgcacgt gtgtcgtatg ggaccgcgcg catttattct
     9481 ctcagctaat tcccctaatc attactaatt gacggcacga tttctgcttc ttacttcctt
     9541 ttactttgga gcttttcatc aataaaacca gtaccatggc cgtacgctca acggaaaagc
     9601 attcaaaaaa acccgcgttc ctcgtgtgat ttgtgggtga gtggcgccat ctattagaga
     9661 atagctgtac tacatctcgt ggacgaaggg gtcagagaag ttgaaagaga gcttgatcga
     9721 ctgctatcca agctaggcga ggaagggaga tcgctagagc aaaagaaaaa aaataagcaa
     9781 atatcttttt ttataacaaa tcgacgttag cgaaatatgt ttgaatcgat ttaacggtta
     9841 gaattccctt tggttcgttc attatgcgat atctaattca attagagact aattcaatta
     9901 gagctaattc aattaggatc caagcttatc gatttcgaac cctcgaccgc cggagtataa
     9961 atagaggcgc ttcgtctacg gagcgacaat tcaattcaaa caagcaaagt gaacacgtcg
    10021 ctaagcgaaa gctaagcaaa taaacaagcg cagctgaaca agctaaacaa tcggggtacc
    10081 gctagagtcg acggtaccgc gggcccggga tccaccggtc gccaccatgg cctcctccga
    10141 gaacgtcatc accgagttca tgcgcttcaa ggtgcgcatg gagggcaccg tgaacggcca
    10201 cgagttcgag atcgagggcg agggcgaggg ccgcccctac gagggccaca acaccgtgaa
    10261 gctgaaggtg accaagggcg gccccctgcc cttcgcctgg gacatcctgt ccccccagtt
    10321 ccagtacggc tccaaggtgt acgtgaagca ccccgccgac atccccgact acaagaagct
    10381 gtccttcccc gagggcttca agtgggagcg cgtgatgaac ttcgaggacg gcggcgtggc
    10441 gaccgtgacc caggactcct ccctgcagga cggctgcttc atctacaagg tgaagttcat
    10501 cggcgtgaac ttcccctccg acggccccgt gatgcagaag aagaccatgg gctgggaggc
    10561 ctccaccgag cgcctgtacc cccgcgacgg cgtgctgaag ggcgagacgc acaaggccct
    10621 gaagctgaag gacggcggcc actacctggt ggagttcaag tccatctaca tggccaagaa
    10681 gcccgtgcag ctgcccggct actactacgt ggacgccaag ctggacatca cctcccacaa
    10741 cgaggactac accatcgtgg agcagtacga gcgcaccgag ggccgccacc acctgttcct
    10801 gagatctcga cccaagaaaa agcggaaggt ggaggacccg taagatccac cggatctaga
    10861 taactgatca taatcagcca taccacattt gtagaggttt tacttgcttt aaaaaacctc
    10921 ccacacctcc ccctgaacct gaaacataaa atgaatgcaa ttgttgttgt taacttgttt
    10981 attgcagctt ataatggtta caaataaagc aatagcatca caaatttcac aaataaagca
    11041 tttttttcac tgcattctag ttgtggtttg tccaaactca tcaatgtatc ttaacgccta
    11101 gtgctgattc aaagtgtgtc tgcaggtatg ttcctataca ttacgcgtga aatatcgtca
    11161 gggatggaag agatggcgaa ggttctcagc attggagcat ccacaagttg ttggtgatgc
    11221 taacattgct tccgagcaat gtattgttgc cgagggtgtc gattagtaca gcaaagcgcg
    11281 agaaacaagt ggctgctgtg gccggcggta cccataaacc agttgagata atttagacaa
    11341 taaattgtaa taggcaattt gatgacgtcc cgcatgtacg gatcgtcaac gctgcacgct
    11401 cgggaatagg gtgagaagtg ctgcacggtt actgaacagc gcaggtggtg attgtgttgt
    11461 tttggtatca cgaaaacaag atagaaactg agaaaaaaga acctttttat cactttaccc
    11521 gcaccgatac tggtctttcc tggatctttt cggggcgaag caaacaactc agcacgacat
    11581 acaaattttg ggataatccc ttagtcagtg cctgtgtctt tctaatgaag attgacaatg
    11641 gcggcggcgg cggcggtggt agcaaccaac gcactggcct ttgttttcat acttttcaat
    11701 gaattcaatt agcttaacca acgaggttgt tgttgatgtt ctcacggaag tcaggagaag
    11761 ctccggcggg cgtgtagggt aagtcgcagt tcggcagcat tctggcagtg tacatccagc
    11821 acaccgagcg ttctagaacg ctgaatcgtt gaaacagagc aaatgactga agggctgtgt
    11881 ctattgctgg gctgtacaga atgacgcatt tcgtgtggca atgtgttttc gaaagattga
    11941 agcattctga tagcgccgtg tgtcaggaat gtaaatattt agatactgca gctaactaca
    12001 aactgagaga gcttccagca agcaggtgta cctacgtggt gatcttttct agaaagaatg
    12061 tagaattgac atgttgtgat gatttaccaa aaacaatttg agaagctaga gagtggattg
    12121 ttctagccac cggaaccatc tggaatgttg aagttttctc gggaagaata gtagacaaaa
    12181 agaaggaggg cttatcccct atagtgagtc gtattacatg gtcatagctg tttcctggca
    12241 gctctggccc gtgtctcaaa atctctgatg ttacattgca caagataaaa atatatcatc
    12301 atgaacaata aaactgtctg cttacataaa cagtaataca aggggtgtta tgagccatat
    12361 tcaacgggaa acgtcgaggc cgcgattaaa ttccaacatg gatgctgatt tatatgggta
    12421 taaatgggct cgcgataatg tcgggcaatc aggtgcgaca atctatcgct tgtatgggaa
    12481 gcccgatgcg ccagagttgt ttctgaaaca tggcaaaggt agcgttgcca atgatgttac
    12541 agatgagatg gtcagactaa actggctgac ggaatttatg cctcttccga ccatcaagca
    12601 ttttatccgt actcctgatg atgcatggtt actcaccact gcgatccccg gaaaaacagc
    12661 attccaggta ttagaagaat atcctgattc aggtgaaaat attgttgatg cgctggcagt
    12721 gttcctgcgc cggttgcatt cgattcctgt ttgtaattgt ccttttaaca gcgatcgcgt
    12781 atttcgtctc gctcaggcgc aatcacgaat gaataacggt ttggttgatg cgagtgattt
    12841 tgatgacgag cgtaatggct ggcctgttga acaagtctgg aaagaaatgc ataaactttt
    12901 gccattctca ccggattcag tcgtcactca tggtgatttc tcacttgata accttatttt
    12961 tgacgagggg aaattaatag gttgtattga tgttggacga gtcggaatcg cagaccgata
    13021 ccaggatctt gccatcctat ggaactgcct cggtgagttt tctccttcat tacagaaacg
    13081 gctttttcaa aaatatggta ttgataatcc tgatatgaat aaattgcagt ttcatttgat
    13141 gctcgatgag tttttctaat cagaattggt taattggttg taacactggc agagcattac
    13201 gctgacttga cgggacggcg caagctcatg accaaaatcc cttaacgtga gttacgcgtc
    13261 gttccactga gcgtcagacc ccgtagaaaa gatcaaagga tcttcttgag atcctttttt
    13321 tctgcgcgta atctgctgct tgcaaacaaa aaaaccaccg ctaccagcgg tggtttgttt
    13381 gccggatcaa gagctaccaa ctctttttcc gaaggtaact ggcttcagca gagcgcagat
    13441 accaaatact gttcttctag tgtagccgta gttaggccac cacttcaaga actctgtagc
    13501 accgcctaca tacctcgctc tgctaatcct gttaccagtg gctgctgcca gtggcgataa
    13561 gtcgtgtctt accgggttgg actcaagacg atagttaccg gataaggcgc agcggtcggg
    13621 ctgaacgggg ggttcgtgca cacagcccag cttggagcga acgacctaca ccgaactgag
    13681 atacctacag cgtgagctat gagaaagcgc cacgcttccc gaagggagaa aggcggacag
    13741 gtatccggta agcggcaggg tcggaacagg agagcgcacg agggagcttc cagggggaaa
    13801 cgcctggtat ctttatagtc ctgtcgggtt tcgccacctc tgacttgagc gtcgattttt
    13861 gtgatgctcg tcaggggggc ggagcctatg gaaaaacgcc agcaacgcgg cctttttacg
    13921 gttcctggcc ttttgctggc cttttgctca catgtt
//
```

**Supplemental File 1D: *SagGD^zpg^***

```
LOCUS       SagGDzpg                16588 bp    DNA     circular       14-DEC-2021
FEATURES             Location/Qualifiers
     misc_feature    complement(603..727)
                     /note="attR4"
     misc_feature    537..552
                     /note="M13F"
     misc_feature    complement(14849..14867)
                     /note="M13R"
     misc_feature    843..2587
                     /note="5' homology arm saglin"
                     /note="saglin ORF"
     misc_feature    complement(2568..2587)
                     /note="5' flanking reverse primer"
     misc_feature    complement(2753..3073)
                     /note="U6 promoter"
     misc_feature    complement(2621..2732)
                     /note="gRNA template"
     misc_feature    2621..3073
                     /note="p609"
     misc_feature    2733..2752
                     /note="linker EM661/662"
     misc_feature    2621..2649
                     /note="EM556"
     misc_feature    complement(3206..3526)
                     /note="U6 promoter"
     misc_feature    complement(3074..3185)
                     /note="gRNA template"
     misc_feature    3074..3526
                     /note="p610"
     misc_feature    3186..3205
                     /note="linker EM663/664"
     misc_feature    complement(3527..3634)
                     /note="gRNA template"
     misc_feature    3527..3979
                     /note="p611"
     misc_feature    complement(3658..3979)
                     /note="U6 promoter"
     misc_feature    3635..3658
                     /note="Linker EM665/666"
     misc_feature    3980..3995
                     /note="EM723"
     regulatory      6418..7491
                     /regulatory_class="promoter"
                     /note="Anopheles gambiae zero population growth"
     gene            7498..11769
                     /gene="hCas9"
     CDS             7498..11769
                     /gene="hCas9"
                     /codon_start=1
                     /transl_table=11
                     /product="hCas9"
                     /protein_id="AXV43861.1"
     misc_feature    complement(7458..7491)
                     /note="EM1384"
     regulatory      11778..12815
                     /regulatory_class="terminator"
                     /note="Anopheles gambiae zero population growth"
     misc_feature    12822..14762
                     /note="Saglin 3' homology arm"
     misc_feature    complement(4467..4574)
                     /label=gRNA template
                     /note="gRNA template"
     misc_feature    4467..4483
                     /label=EM701
                     /note="EM701"
     misc_feature    complement(4441..4466)
                     /label=EM702
                     /note="EM702"
     misc_feature    complement(4392..4461)
                     /label=tRNA-gly
                     /note="tRNA-gly (missing terminal A, which is added when
                     cloning guide linker)"
     misc_feature    complement(4371..4395)
                     /label=EM1566
                     /note="EM1566"
     misc_feature    complement(4284..4391)
                     /label=gRNA template
                     /note="gRNA template"
     misc_feature    4367..4391
                     /label=EM1567
                     /note="EM1567"
     misc_feature    4284..4300
                     /label=EM703
                     /note="EM703"
     misc_feature    complement(4258..4283)
                     /label=EM704
                     /note="EM704"
     misc_feature    complement(4209..4278)
                     /label=tRNA-gly
                     /note="tRNA-gly (missing terminal A, which is added when
                     cloning guide linker)"
     misc_feature    complement(4188..4212)
                     /label=EM1568
                     /note="EM1568"
     misc_feature    complement(4101..4208)
                     /label=gRNA template
                     /note="gRNA template"
     misc_feature    4184..4208
                     /label=EM1569
                     /note="EM1569"
     misc_feature    complement(4086..4110)
                     /label=EM850
                     /note="EM850"
     misc_feature    complement(4025..4095)
                     /label=tRNA-gly
                     /note="tRNA-gly"
     misc_feature    4018..4041
                     /label=EM717
                     /note="EM717"
     misc_feature    4018..4019
                     /note="EM1760"
     misc_feature    complement(4833..5157)
                     /label=U6 promoter (ag)
                     /note="U6 promoter (ag)"
     primer_bind     complement(4876..4894)
                     /label=EM1084
     misc_feature    complement(4807..4832)
                     /label=EM700
                     /note="EM700"
     misc_feature    complement(4758..4827)
                     /label=tRNA-gly
                     /note="tRNA-gly (missing terminal A, which is added when
                     cloning guide linker)"
     misc_feature    complement(4737..4761)
                     /label=EM1562
                     /note="EM1562"
     misc_feature    complement(4650..4757)
                     /label=gRNA template
                     /note="gRNA template"
     misc_feature    4733..4757
                     /label=EM1563
                     /note="EM1563"
     primer_bind     4740..4757
                     /label=EM1392
     misc_feature    4650..4666
                     /label=EM705
                     /note="EM705"
     misc_feature    complement(4624..4649)
                     /label=EM924
                     /note="EM924"
     misc_feature    complement(4575..4644)
                     /label=tRNA-gly
                     /note="tRNA-gly (missing terminal A, which is added when
                     cloning guide linker)"
     misc_feature    complement(4554..4578)
                     /label=EM1564
                     /note="EM1564"
     misc_feature    4550..4574
                     /label=EM1565
                     /note="EM1565"
     misc_feature    complement(5135..5157)
                     /label=EM895
                     /note="EM895"
     primer_bind     complement(5135..5157)
                     /label=EM895
     misc_feature    5190..6413
                     /note="3xP3 DsRedNLS sv40 from 1154"
ORIGIN
        1 CTTTCCTGCG TTATCCCCTG ATTCTGTGGA TAACCGTATT ACCGCCTTTG AGTGAGCTGA
       61 TACCGCTCGC CGCAGCCGAA CGACCGAGCG CAGCGAGTCA GTGAGCGAGG AAGCGGAAGA
      121 GCGCCCAATA CGCAAACCGC CTCTCCCCGC GCGTTGGCCG ATTCATTAAT GCAGCTGGCA
      181 CGACAGGTTT CCCGACTGGA AAGCGGGCAG TGAGCGCAAC GCAATTAATA CGCGTACCGC
      241 TAGCCAGGAA GAGTTTGTAG AAACGCAAAA AGGCCATCCG TCAGGATGGC CTTCTGCTTA
      301 GTTTGATGCC TGGCAGTTTA TGGCGGGCGT CCTGCCCGCC ACCCTCCGGG CCGTTGCTTC
      361 ACAACGTTCA AATCCGCTCC CGGCGGATTT GTCCTACTCA GGAGAGCGTT CACCGACAAA
      421 CAACAGATAA AACGAAAGGC CCAGTCTTCC GACTGAGCCT TTCGTTTTAT TTGATGCCTG
      481 GCAGTTCCCT ACTCTCGCGT TAACGCTAGC ATGGATGTTT TCCCAGTCAC GACGTTGTAA
      541 AACGACGGCC AGTCTTAAGC TCGGGCCCCT ACAGGTCACT AATACCATCT AAGTAGTTGA
      601 TTCATAGTGA CTGGATATGT TGTGTTTTAC AGTATTATGT AGTCTGTTTT TTATGCAAAA
      661 TCTAATTTAA TATATTGATA TTTATATCAT TTTACGTTTC TCGTTCAACT TTTCTATACA
      721 AAGTTGGTAC CGGGCCCCCC GCTAGCGTCG ACGGTATCGA TAAGCTTGAT cggatccTGC
      781 GGGTGCCAGG GCGTGCCCTT GGGCTCCCCG GGCGCGTACT CCACCTCACC CATaggcctg
      841 ataacaCACT GGCACGACGA TCATCCCTTT CTGCTCGAAA CGGTCGCGAA GGCGTACGAG
      901 CCGCTCAAGG CCGCGCTGGA GTCGGAGCAG GATGCGCATC GAAGCCTTCG GCTGGTGCCC
      961 GAGCTGCAGC GCAAGCAGCT CATCTACAGC ATCGAGGCGG GCCGGCTCGA GGACGCCCAC
     1021 ATCCTGCACA TGACGCTCAA GGGCCAGTGG AAGCCGGACC AAATCGTGTC GGCCATCCAG
     1081 GACGGCTACC ACGTCAACCC GACGATCATG GAGCATCTGC TCGAGTTCGT CCGCGCCATC
     1141 CCGGTGCGGA AGGAGCGGGC CGCCTACTAC AAAGCGATCG GGCCGGTCAT TCGCAACTTC
     1201 AAGCTGACCC TGACGTACGT GACGCTCCTG TTCGCGGGCG ATGCGACCGG CGTGTTTGAC
     1261 ACGCGAAAGG ATCGCGACGA CTACATCTCG CAACCGCTCA CCGTGTTCGT GCACATGCTG
     1321 CGCTGGCAGC TGGCGAACCT CGAGTTCGAG TTCCACCTGT CGCTGGCGGA GCGCTTCCCC
     1381 CGCTACTACT CGCTGCACAT CGAGCAAATC TTCTTCTTTG CGCCCCCGCA CTGGCAGAAG
     1441 GCGGAGAAGC GCCGGCTGTT CGAGATCGCG AGCCTGTTCA AGGCGAAGGG GCACCGGTTT
     1501 GCGGCGATCG AGCAGCTGCT GAAGTGGGCG CACCAGTACG CACCGATGCG CGGTGGCAAA
     1561 GCCCACCTGG AGGAAATACT GCCCACGCTG GCGCTCGAAA CGGAGAAGCT GCGGCGCACG
     1621 GTCGAGCAGA CGGGCAAGAG CTCAGCGGAG CTGGTGCGGC TCAAGAAGCT GGAGGAAAAG
     1681 TTCGTAAGCA AAAAGGATCG CTGGAACACG AACACCTACC GCACGTACCT GCACATGATC
     1741 AAGACGAAGG GTAAGTTCGG CCAGCATAGG ATGGAAATGG AGCGCAAGGG TTAAATAGCG
     1801 GGAATGGCAC ATGCGAGGCG ATGGATGAAA TACAAGTCGT TTGGTGAGCG GAGTCACCAA
     1861 GCAAGCATAA CTTTCATGCG TACAATGTGT ATGATCACTG ATGACAAAGC GCTTGGGTGG
     1921 GGCGCTTGCT GCTGAAGCGT GGCTGACAAA TGGGGGGGGG GCTTGAACAA ATATTTGCAC
     1981 ACACGATAAT GCTCTTGTTT GCCATGCGCT GACGGAAGAC AAATCGCTGC AAGCGACGGC
     2041 TCAAACCTGC TCAAACCAGG GAAGGTTTGC ATAAGGTTCA CTTATAAATG AGCGTTCGTG
     2101 GCTACTCGGG CGTTCAAGTC ATCTCGAGCA GGAAGCACCG CAGCATGTCA CGTTTGCCCA
     2161 CTGTACTGTT GCTCTTAGCG AGCGCCGCAG TCCTCGCGGC TGGCGGCCAG GAAGCAACCG
     2221 AAGACCCGTT CGCGGACGAA ACCGACCAGT GCCAGATCAG CGTTAGTGCC GAGACGATGA
     2281 AATCCCTCCA CGGCGGGTCG ATGCAGCCCG ACGGCACGTG CGACAACCTG TGGGAAAGCT
     2341 TTCTATCTCA GTTTCACCAA GTCAGGGAAA ACCTGACCGC GTGCCAGGAG CGGGCAGCCG
     2401 CCGGACCAGC GCCCGATCCT TCCAGCCAGT TCTGTCAGCA GCTGCTGGAC GATGCGCAGC
     2461 GGCAGATGGA GCAGGAGCAT CGCCAGTACG CCGCCACCCT GGAGGAGCAG CTGCACGCAG
     2521 CGCAGCAGGA AACCCAGCAG GAGCAGGAGA TGAAGAAGGC GCTGCAGAAG CAGCTCGACG
     2581 CGCTCACGTA AGCGTCGACG GTATCGATAA GCTTGATCgg atccGATGAA AATAAGAAAA
     2641 ACATTTGACA AAAAAAGCAC CGACTCGGTG CCACTTTTTC AAGTTGATAA CGGACTAGCC
     2701 TTATTTTAAC TTGCTATTTC TAGCTCTAAA ACCCTGCTTG AAGTTACCGT TCAAGGACGA
     2761 GGGGAAAAAA GGTTGTATAT ATACTTTGCG CTTTCAATCC TTGCTCTAGC GATGCATAAA
     2821 GGATATTCAA GAAGGATTTT TTGCCTGAGT GTGACTGTTA TAAGGTTTTG ATCCTGCCGT
     2881 AACTCACGCT AATGTGATGT TTCATAAACT TTTCAACATT TCTTGTATTG CTTCATCTCT
     2941 TTTTTAATCA AAAGTTACTA TTCATTGTAA TACGTTGAAA TATCAAAAAA GAATAAACAG
     3001 TTTTCCTATG ATTAATTCAA AAACAGAGTT TTCTAGAATA ACCAAGAGCC AAACCGCTTC
     3061 AGTGTATATG TGGggaaGAT GAAAATAAGA AAAACATTTG ACAAAAAAAG CACCGACTCG
     3121 GTGCCACTTT TTCAAGTTGA TAACGGACTA GCCTTATTTT AACTTGCTAT TTCTAGCTCT
     3181 AAAACCAACG CGCTGTACAT CGATCAAGGA CGAGGGGAAA AAAGGTTGTA TATATACTTT
     3241 GCGCTTTCAA TCCTTGCTCT AGCGATGCAT AAAGGATATT CAAGAAGGAT TTTTTGCCTG
     3301 AGTGTGACTG TTATAAGGTT TTGATCCTGC CGTAACTCAC GCTAATGTGA TGTTTCATAA
     3361 ACTTTTCAAC ATTTCTTGTA TTGCTTCATC TCTTTTTTAA TCAAAAGTTA CTATTCATTG
     3421 TAATACGTTG AAATATCAAA AAAGAATAAA CAGTTTTCCT ATGATTAATT CAAAAACAGA
     3481 GTTTTCTAGA ATAACCAAGA GCCAAACCGC TTCAGTGTAT ATGTGGagag GATGAAAATA
     3541 AGAAAAACAT TTGACAAAAA AAGCACCGAC TCGGTGCCAC TTTTTCAAGT TGATAACGGA
     3601 CTAGCCTTAT TTTAACTTGC TATTTCTAGC TCTAAAACCT CGTACTACAA CCTGATGCAA
     3661 GGACGAGGGG AAAAAAGGTT GTATATATAC TTTGCGCTTT CAATCCTTGC TCTAGCGATG
     3721 CATAAAGGAT ATTCAAGAAG GATTTTTTGC CTGAGTGTGA CTGTTATAAG GTTTTGATCC
     3781 TGCCGTAACT CACGCTAATG TGATGTTTCA TAAACTTTTC AACATTTCTT GTATTGCTTC
     3841 ATCTCTTTTT TAATCAAAAG TTACTATTCA TTGTAATACG TTGAAATATC AAAAAAGAAT
     3901 AAACAGTTTT CCTATGATTA ATTCAAAAAC AGAGTTTTCT AGAATAACCA AGAGCCAAAC
     3961 CGCTTCAGTG TATATGTGGa acaCACTGGC ACGACATCGT CCCCACCGCC AGCGAATaaa
     4021 aaaatgcatc ggccgggaat cgaacccggg ccgcccgcgt ggcaggcgag cattctacca
     4081 ctgaaccacc gatgcttcgt agcaccgact cggtgccact ttttcaagtt gataacggac
     4141 tagccttatt tcaacttgct atgctgtttc cagcatagct ctgaaactcc ctgacgatat
     4201 ttcacgctgc atcggccggg aatcgaaccc gggccgcccg cgtggcaggc gagcattcta
     4261 ccactgaacc accgatgctt tccagcaccg actcggtgcc actttttcaa gttgataacg
     4321 gactagcctt atttcaactt gctatgctgt ttccagcata gctctgaaac gcggacacac
     4381 tttgaatcag tgcatcggcc gggaatcgaa cccgggccgc ccgcgtggca ggcgagcatt
     4441 ctaccactga accaccgatg ctatggagca ccgactcggt gccacttttt caagttgata
     4501 acggactagc cttatttcaa cttgctatgc tgtttccagc atagctctga aacggaagct
     4561 ccagagcagc ctctgcatcg gccgggaatc gaacccgggc cgcccgcgtg gcaggcgagc
     4621 attctaccac tgaaccaccg atgctctcta gcaccgactc ggtgccactt tttcaagttg
     4681 ataacggact agccttattt caacttgcta tgctgtttcc agcatagctc tgaaacagga
     4741 cccacatcgt tccgtgtgca tcggccggga atcgaacccg ggccgcccgc gtggcaggcg
     4801 agcattctac cactgaacca ccgatgcttt acaaggacga ggggaaaaaa ggttgtatat
     4861 atactttgcg ctttcaatcc ttgctctagc gatgcataaa ggatattcaa gaaggatttt
     4921 ttgcctgagt gtgactgtta taaggttttg atcctgccgt aactcacgct aatgtgatgt
     4981 ttcataaact tttcaacatt tcttgtattg cttcatctct tttttaatca aaagttacta
     5041 ttcattgtaa tacgttgaaa tatcaaaaaa gaataaacag ttttcctatg attaattcaa
     5101 aaacagagtt ttctagaata accaagagcc aaaccgcttc agtgtatatg tggataaACC
     5161 CACATCGTTC CGTATCTGAG ACCGATATCT AATTCAATTA GAGACTAATT CAATTAGAGC
     5221 TAATTCAATT AGGATCCAAG CTTATCGATT TCGAACCCTC GACCGCCGGA GTATAAATAG
     5281 AGGCGCTTCG TCTACGGAGC GACAATTCAA TTCAAACAAG CAAAGTGAAC ACGTCGCTAA
     5341 GCGAAAGCTA AGCAAATAAA CAAGCGCAGC TGAACAAGCT AAACAATCGG GGTACCGCTA
     5401 GAGTCGACGG TACCGCGGGC CCGGGATCCA ccggtcgcca ccatggcctc ctccgagaac
     5461 gtcatcaccg agttcatgcg cttcaaggtg cgcatggagg gcaccgtgaa cggccacgag
     5521 ttcgagatcg agggcgaggg cgagggccgc ccctacgagg gccacaacac cgtgaagctg
     5581 aaggtgacca agggcggccc cctgcccttc gcctgggaca tcctgtcccc ccagttccag
     5641 tacggctcca aggtgtacgt gaagcacccc gccgacatcc ccgactacaa gaagctgtcc
     5701 ttccccgagg gcttcaagtg ggagcgcgtg atgaacttcg aggacggcgg cgtggcgacc
     5761 gtgacccagg actcctccct gcaggacggc tgcttcatct acaaggtgaa gttcatcggc
     5821 gtgaacttcc cctccgacgg ccccgtgatg cagaagaaga ccatgggctg ggaggcctcc
     5881 accgagcgcc tgtacccccg cgacggcgtg ctgaagggcg agacgcacaa ggccctgaag
     5941 ctgaaggacg gcggccacta cctggtggag ttcaagtcca tctacatggc caagaagccc
     6001 gtgcagctgc ccggctacta ctacgtggac gccaagctgg acatcacctc ccacaacgag
     6061 gactacacca tcgtggagca gtacgagcgc accgagggcc gccaccacct gttcctgaga
     6121 tctcgaccca agaaaaagcg gaaggtggag gacccgtaag atccaccgga tctagataac
     6181 tgatcataat cagccatacc acatttgtag aggttttact tgctttaaaa aacctcccac
     6241 acctccccct gaacctgaaa cataaaatga atgcaattgt tgttgttaac ttgtttattg
     6301 cagcttataa tggttacaaa taaagcaata gcatcacaaa tttcacaaat aaagcatttt
     6361 tttcactgca ttctagttgt ggtttgtcca aactcatcaa tgtatcttaT TATcggtcag
     6421 cgctggcggt ggggacagct ccggctgtgg ctgttcttgc gagtcctctt cctgcggcac
     6481 atccctctcg tcgaccagtt cagtttgctg agcgtaagcc tgctgctgtt cgtcctgcat
     6541 catcgggacc atttgtatgg gccatccgcc accaccacca tcaccaccgc cgtccatttc
     6601 taggggcata cccatcagca tctccgcggg cgccattggc ggtggtgcca aggtgccatt
     6661 cgtttgttgc tgaaagcaaa agaaagcaaa ttagtgttgt ttctgctgca cacgataatt
     6721 ttcgtttctt gccgctagac acaaacaaca ctgcatctgg agggagaaat ttgacgccta
     6781 gctgtataac ttacctcaaa gttattgtcc atcgtggtat aatggaccta ccgagcccgg
     6841 ttacactaca caaagcaaga ttatgcgaca aaatcacagc gaaaactagt aattttcatc
     6901 tatcgaaagc ggccgagcag agagttgttt ggtattgcaa cttgacattc tgctgcggga
     6961 taaaccgcga cgggctacca tggcgcacct gtcagatggc tgtcaaattt ggcccggttt
     7021 gcgatatgga gtgggtgaaa ttatatccca ctcgctgatc gtgaaaatag acacctgaaa
     7081 acaataattg ttgtgttaat tttacatttt gaagaacagc acaagttttg ctgacaatat
     7141 ttaattacgt ttcgttatca acggcacgga aagattatct cgctgattat ccctctcgct
     7201 ctctctgtct atcatgtcct ggtcgttctc gcgtcacccc ggataatcga gagacgccat
     7261 ttttaatttg aactactaca ccgacaagca tgccgtgagc tctttcaagt tcttctgtcc
     7321 gaccaaagaa acagagaata ccgcccggac agtgcccgga gtgatcgatc catagaaaat
     7381 cgcccatcat gtgccactga ggcgaaccgg cgtagcttgt tccgaatttc caagtgcttc
     7441 cccgtaacat ccgcatataa caaacagccc aacaacaaat acagcatcga gctcgagatg
     7501 gactataagg accacgacgg agactacaag gatcatgata ttgattacaa agacgatgac
     7561 gataagatgg ccccaaagaa gaagcggaag gtcggtatcc acggagtccc agcagccgac
     7621 aagaagtaca gcatcggcct ggacatcggc accaactctg tgggctgggc cgtgatcacc
     7681 gacgagtaca aggtgcccag caagaaattc aaggtgctgg gcaacaccga ccggcacagc
     7741 atcaagaaga acctgatcgg agccctgctg ttcgacagcg gcgaaacagc cgaggccacc
     7801 cggctgaaga gaaccgccag aagaagatac accagacgga agaaccggat ctgctatctg
     7861 caagagatct tcagcaacga gatggccaag gtggacgaca gcttcttcca cagactggaa
     7921 gagtccttcc tggtggaaga ggataagaag cacgagcggc accccatctt cggcaacatc
     7981 gtggacgagg tggcctacca cgagaagtac cccaccatct accacctgag aaagaaactg
     8041 gtggacagca ccgacaaggc cgacctgcgg ctgatctatc tggccctggc ccacatgatc
     8101 aagttccggg gccacttcct gatcgagggc gacctgaacc ccgacaacag cgacgtggac
     8161 aagctgttca tccagctggt gcagacctac aaccagctgt tcgaggaaaa ccccatcaac
     8221 gccagcggcg tggacgccaa ggccatcctg tctgccagac tgagcaagag cagacggctg
     8281 gaaaatctga tcgcccagct gcccggcgag aagaagaatg gcctgttcgg aaacctgatt
     8341 gccctgagcc tgggcctgac ccccaacttc aagagcaact tcgacctggc cgaggatgcc
     8401 aaactgcagc tgagcaagga cacctacgac gacgacctgg acaacctgct ggcccagatc
     8461 ggcgaccagt acgccgacct gtttctggcc gccaagaacc tgtccgacgc catcctgctg
     8521 agcgacatcc tgagagtgaa caccgagatc accaaggccc ccctgagcgc ctctatgatc
     8581 aagagatacg acgagcacca ccaggacctg accctgctga aagctctcgt gcggcagcag
     8641 ctgcctgaga agtacaaaga gattttcttc gaccagagca agaacggcta cgccggctac
     8701 attgacggcg gagccagcca ggaagagttc tacaagttca tcaagcccat cctggaaaag
     8761 atggacggca ccgaggaact gctcgtgaag ctgaacagag aggacctgct gcggaagcag
     8821 cggaccttcg acaacggcag catcccccac cagatccacc tgggagagct gcacgccatt
     8881 ctgcggcggc aggaagattt ttacccattc ctgaaggaca accgggaaaa gatcgagaag
     8941 atcctgacct tccgcatccc ctactacgtg ggccctctgg ccaggggaaa cagcagattc
     9001 gcctggatga ccagaaagag cgaggaaacc atcaccccct ggaacttcga ggaagtggtg
     9061 gacaagggcg cttccgccca gagcttcatc gagcggatga ccaacttcga taagaacctg
     9121 cccaacgaga aggtgctgcc caagcacagc ctgctgtacg agtacttcac cgtgtataac
     9181 gagctgacca aagtgaaata cgtgaccgag ggaatgagaa agcccgcctt cctgagcggc
     9241 gagcagaaaa aggccatcgt ggacctgctg ttcaagacca accggaaagt gaccgtgaag
     9301 cagctgaaag aggactactt caagaaaatc gagtgcttcg actccgtgga aatctccggc
     9361 gtggaagatc ggttcaacgc ctccctgggc acataccacg atctgctgaa aattatcaag
     9421 gacaaggact tcctggacaa tgaggaaaac gaggacattc tggaagatat cgtgctgacc
     9481 ctgacactgt ttgaggacag agagatgatc gaggaacggc tgaaaaccta tgcccacctg
     9541 ttcgacgaca aagtgatgaa gcagctgaag cggcggagat acaccggctg gggcaggctg
     9601 agccggaagc tgatcaacgg catccgggac aagcagtccg gcaagacaat cctggatttc
     9661 ctgaagtccg acggcttcgc caacagaaac ttcatgcagc tgatccacga cgacagcctg
     9721 acctttaaag aggacatcca gaaagcccag gtgtccggcc agggcgatag cctgcacgag
     9781 cacattgcca atctggccgg cagccccgcc attaagaagg gcatcctgca gacagtgaag
     9841 gtggtggacg agctcgtgaa agtgatgggc cggcacaagc ccgagaacat cgtgatcgaa
     9901 atggccagag agaaccagac cacccagaag ggacagaaga acagccgcga gagaatgaag
     9961 cggatcgaag agggcatcaa agagctgggc agccagatcc tgaaagaaca ccccgtggaa
    10021 aacacccagc tgcagaacga gaagctgtac ctgtactacc tgcagaatgg gcgggatatg
    10081 tacgtggacc aggaactgga catcaaccgg ctgtccgact acgatgtgga ccatatcgtg
    10141 cctcagagct ttctgaagga cgactccatc gacaacaagg tgctgaccag aagcgacaag
    10201 aaccggggca agagcgacaa cgtgccctcc gaagaggtcg tgaagaagat gaagaactac
    10261 tggcggcagc tgctgaacgc caagctgatt acccagagaa agttcgacaa tctgaccaag
    10321 gccgagagag gcggcctgag cgaactggat aaggccggct tcatcaagag acagctggtg
    10381 gaaacccggc agatcacaaa gcacgtggca cagatcctgg actcccggat gaacactaag
    10441 tacgacgaga atgacaagct gatccgggaa gtgaaagtga tcaccctgaa gtccaagctg
    10501 gtgtccgatt tccggaagga tttccagttt tacaaagtgc gcgagatcaa caactaccac
    10561 cacgcccacg acgcctacct gaacgccgtc gtgggaaccg ccctgatcaa aaagtaccct
    10621 aagctggaaa gcgagttcgt gtacggcgac tacaaggtgt acgacgtgcg gaagatgatc
    10681 gccaagagcg agcaggaaat cggcaaggct accgccaagt acttcttcta cagcaacatc
    10741 atgaactttt tcaagaccga gattaccctg gccaacggcg agatccggaa gcggcctctg
    10801 atcgagacaa acggcgaaac cggggagatc gtgtgggata agggccggga ttttgccacc
    10861 gtgcggaaag tgctgagcat gccccaagtg aatatcgtga aaaagaccga ggtgcagaca
    10921 ggcggcttca gcaaagagtc tatcctgccc aagaggaaca gcgataagct gatcgccaga
    10981 aagaaggact gggaccctaa gaagtacggc ggcttcgaca gccccaccgt ggcctattct
    11041 gtgctggtgg tggccaaagt ggaaaagggc aagtccaaga aactgaagag tgtgaaagag
    11101 ctgctgggga tcaccatcat ggaaagaagc agcttcgaga agaatcccat cgactttctg
    11161 gaagccaagg gctacaaaga agtgaaaaag gacctgatca tcaagctgcc taagtactcc
    11221 ctgttcgagc tggaaaacgg ccggaagaga atgctggcct ctgccggcga actgcagaag
    11281 ggaaacgaac tggccctgcc ctccaaatat gtgaacttcc tgtacctggc cagccactat
    11341 gagaagctga agggctcccc cgaggataat gagcagaaac agctgtttgt ggaacagcac
    11401 aagcactacc tggacgagat catcgagcag atcagcgagt tctccaagag agtgatcctg
    11461 gccgacgcta atctggacaa agtgctgtcc gcctacaaca agcaccggga taagcccatc
    11521 agagagcagg ccgagaatat catccacctg tttaccctga ccaatctggg agcccctgcc
    11581 gccttcaagt actttgacac caccatcgac cggaagaggt acaccagcac caaagaggtg
    11641 ctggacgcca ccctgatcca ccagagcatc accggcctgt acgagacacg gatcgacctg
    11701 tctcagctgg gaggcgacaa aaggccggcg gccacgaaaa aggccggcca ggcaaaaaag
    11761 aaaaagtaat taattaagag gacggcgaga agtaatcata tgtccgcatt ttgcgcaaac
    11821 caggcgctta gacaatttgc gcgtaagcac attcgaaatg tgcaaagctg aaagcagtgg
    11881 tttcgccagc ccgagttcag cgaaacggat tccttccaag tgtttgcatt cctggcggag
    11941 tattcctccc aaaatgcact caccctgcgt gcagtgccaa atcgtgagtt tcctaatttt
    12001 ttcatattgt ttattaccta ccaactaaag ttgttgttat atattgcgtt ttacgtacga
    12061 caaataagtt cgtattcaga aatatttgcg ataagagaga actcatttgc gatgaatctc
    12121 attgtattta gctaagtgcc ttgataagta agcggaacag caggaatatg acactccttg
    12181 ggaaatacat gtaagcgtct gtgaattaga tatatataca cgcaaccaaa tggtccatgg
    12241 ttgatttaag cactgcctgt tgtcgaacat tgctataagc aaaataaaga agcattcatt
    12301 aatctaaaat ttcttcaaag tgacttcaat gatgatctct aggctatagt gaaagctgaa
    12361 agcttattcg acaatgcaag ggaaagtgac gcacgtgtgt cgtatgggac cgcgcgcatt
    12421 tattctctca gctaattccc ctaatcatta gtaattgacg gcacgatttc tgcttcttac
    12481 ttccttttac tttggagctt ttcatcaata aaaccagtac catggccgta cgctcaacgg
    12541 aaaagcattc acaaaaaccc gcgttcctcg tgtgatttgt gggtgagtgg cgccatctat
    12601 tagagaatag ctgtactaca tctcgtggac gaaggggtca gagaagttga aagagagctt
    12661 gatcgactgc tatccaagct aggcgaggaa gggagatcgc tagagcaaaa gaaaaaaaat
    12721 aagcaaatat ctttttttat aacaaatcga cgttagcgaa atatgtttga atcgatttaa
    12781 cggttagaat tccctttggt tcgttcatta tgcgaggcgc gacgcAGCTG CAAACCAGCC
    12841 TCGCCACGAT CGAGCTGAAG CACTGGCGCA AGTTCGACCG GTTCGTACCG TACGCTAACG
    12901 CGCTGCCCCA GCCAGCCCAG CGGCTGGAGG TGCTGCGTGT GCTGCTGAGC CAGATCGGCG
    12961 ACCGGGAGAA GAAGACTTCC CACAAATATC TGGTAAAGGC GGCACGGCAG TTTGACATTT
    13021 GCGAGCAGTT TATCGGCCGG GGCAAGGTGG ACCAGGCGGT CAAGAAGCAG CTGGAGGAAC
    13081 TGCGCGGGAA GTTTGCAACG TTTGCGAAGG GCAAGAACTA CCAGCACTAT CTGAGCGAGT
    13141 CGCGCAAGTC GTCCGGCTAG GCGTCGGCGA CGGTCTAAAA GTGATTCAAA GCCCCGTAGG
    13201 CGAGGAACCC CCGTACGTGT CGTGTAGCGC AAACCCTTCA CCATCATAGC ACAGTGAAAT
    13261 AAAGCAAGAA TAAAAATTAA AAAAAAAAAC AATTGTAATC GAAATATTGA GCAACAGCAA
    13321 AGCAAAGCCT TTCCTTTGGA TCATCCGCAA TCGGCACGTT GCGAGTTGGG CGGCAGCGTT
    13381 GCTTGTGCAA TCGCTGGACC CATGCTGGAC AAACAGATGA GCACACTGGC CTTCCCAGCC
    13441 AGCTAATATT GACACTTCAT TCGAGCTGGA CCATTAATCA CGGCGTCTGT TTGCCTTTGG
    13501 GCCAAGTATA ACTAGCACCA CGCACCGAAC AGACGGGCTC TAGTTTTGTT TCTCTTGCTG
    13561 TGAGCGTCAG TTAATTAAAA AAACAAACCT ATGGATCACA GCAGAGTGTG GTGCTGTCTG
    13621 CTTAGCTTCA TCGCAATGGC CGTGCTGTGT AGCGCATCCA GTGCAGCATG CACGCTCCAG
    13681 GTCCCCGAAA CCATGATAAC GCGCCTGCTG CAAGCGGATC AGCCCGCCGG CGGCTGCGCA
    13741 GCTGCTTGGG ATAGTCTACT GTCTCGGCTG GAGCAAATGC GCCACAATCT TACGGGCTGC
    13801 AGCGAGCGCG AAACGACGAA CCCGGCACCA CCCGATGATA CCCGCACCAA CGCCCCCTGT
    13861 CAGCTGCTGC TGAACGAGCT GCAACGGAAA GGGAATCAAA CCCTGCTCCA GCTCGACAAA
    13921 GAAATGGCCA GCAAAGCACT GTACGAGCGC ATCGAGCTGG CCAAGGTGGT GCAGGACCAG
    13981 GACCGCATCC AACAAGCGAT CGAAACGATG CGCACCGAGC AGCGAAACCA CCGGCAGCAG
    14041 CTGCTACTCC TCTACCTCGA TGCGGCGGAC CTACGACGAG CCCTGCACCA GTATAAGCTG
    14101 CTCGTCGCGC AGGGCGATCG CCACCTTCCC CAGCAGATAG TGAAGTTCGT GTACGCCGCC
    14161 CCACGGCACG AGAACCGCCG GCTCGAGAAC CTGCTCGACC TGGTGCGGCA GCTGCCGGCG
    14221 CGCCAGGACC AGCGCACCCT GTACCAGCTG CTGCAGCCGG AAATTATGAA GCGCCCCACG
    14281 CAGAACCAGA GCACCCTCGC CATGCTGACA GCGCTCGAGA TGGGCCAGGT CGTGGAGGGG
    14341 AACGGCGAGC TGAAAAAGCA GCAGGACGCG ATGTACCAGC TGGTGCTGAA GCGCTGGATG
    14401 TTTCTGTGCC TCGCCGGCCA GTATCGCGAG ATAGTGCAGT TTGCCACGAA GCACCCGCGC
    14461 CTGTTCGAGC AGATCGGGGC GAAGATCGCC ACGATTCATC CGAAGTACTG GTGGAAGTTT
    14521 AGCTTCACCC AGTTCGTGAC GTACCCGAAC CTGTTGCCGC TGCCCGAGCA GCGGCTGGAA
    14581 GCGTTCCGCA CCATCATGAA GCAGCTGAAG CAGCGCAATG GGAAGTTTTT CGCCGACCAT
    14641 CTGGCGAAGC TGGCACCCCA GATCGAGCGC TGCGAGCAGT TTCTGCGGCA GCAGAAGAAG
    14701 GAGCTGGAGT GGAAGGACGA GCTGGTGAAG CTGAAGGGAC AGTTTGCCGA CTTTGACCgc
    14761 ttatcATGGG TGAGGTGGAG TACGCGCCCG GGGAGCCCAA GGGCACGCCC TGGCACCCGC
    14821 AgcttATCCC CTATAGTGAG TCGTATTACA TGGTCATAGC TGTTTCCTGG CAGCTCTGGC
    14881 CCGTGTCTCA AAATCTCTGA TGTTACATTG CACAAGATAA AAATATATCA TCATGAACAA
    14941 TAAAACTGTC TGCTTACATA AACAGTAATA CAAGGGGTGT TATGAGCCAT ATTCAACGGG
    15001 AAACGTCGAG GCCGCGATTA AATTCCAACA TGGATGCTGA TTTATATGGG TATAAATGGG
    15061 CTCGCGATAA TGTCGGGCAA TCAGGTGCGA CAATCTATCG CTTGTATGGG AAGCCCGATG
    15121 CGCCAGAGTT GTTTCTGAAA CATGGCAAAG GTAGCGTTGC CAATGATGTT ACAGATGAGA
    15181 TGGTCAGACT AAACTGGCTG ACGGAATTTA TGCCTCTTCC GACCATCAAG CATTTTATCC
    15241 GTACTCCTGA TGATGCATGG TTACTCACCA CTGCGATCCC CGGAAAAACA GCATTCCAGG
    15301 TATTAGAAGA ATATCCTGAT TCAGGTGAAA ATATTGTTGA TGCGCTGGCA GTGTTCCTGC
    15361 GCCGGTTGCA TTCGATTCCT GTTTGTAATT GTCCTTTTAA CAGCGATCGC GTATTTCGTC
    15421 TCGCTCAGGC GCAATCACGA ATGAATAACG GTTTGGTTGA TGCGAGTGAT TTTGATGACG
    15481 AGCGTAATGG CTGGCCTGTT GAACAAGTCT GGAAAGAAAT GCATAAACTT TTGCCATTCT
    15541 CACCGGATTC AGTCGTCACT CATGGTGATT TCTCACTTGA TAACCTTATT TTTGACGAGG
    15601 GGAAATTAAT AGGTTGTATT GATGTTGGAC GAGTCGGAAT CGCAGACCGA TACCAGGATC
    15661 TTGCCATCCT ATGGAACTGC CTCGGTGAGT TTTCTCCTTC ATTACAGAAA CGGCTTTTTC
    15721 AAAAATATGG TATTGATAAT CCTGATATGA ATAAATTGCA GTTTCATTTG ATGCTCGATG
    15781 AGTTTTTCTA ATCAGAATTG GTTAATTGGT TGTAACACTG GCAGAGCATT ACGCTGACTT
    15841 GACGGGACGG CGCAAGCTCA TGACCAAAAT CCCTTAACGT GAGTTACGCG TCGTTCCACT
    15901 GAGCGTCAGA CCCCGTAGAA AAGATCAAAG GATCTTCTTG AGATCCTTTT TTTCTGCGCG
    15961 TAATCTGCTG CTTGCAAACA AAAAAACCAC CGCTACCAGC GGTGGTTTGT TTGCCGGATC
    16021 AAGAGCTACC AACTCTTTTT CCGAAGGTAA CTGGCTTCAG CAGAGCGCAG ATACCAAATA
    16081 CTGTTCTTCT AGTGTAGCCG TAGTTAGGCC ACCACTTCAA GAACTCTGTA GCACCGCCTA
    16141 CATACCTCGC TCTGCTAATC CTGTTACCAG TGGCTGCTGC CAGTGGCGAT AAGTCGTGTC
    16201 TTACCGGGTT GGACTCAAGA CGATAGTTAC CGGATAAGGC GCAGCGGTCG GGCTGAACGG
    16261 GGGGTTCGTG CACACAGCCC AGCTTGGAGC GAACGACCTA CACCGAACTG AGATACCTAC
    16321 AGCGTGAGCT ATGAGAAAGC GCCACGCTTC CCGAAGGGAG AAAGGCGGAC AGGTATCCGG
    16381 TAAGCGGCAG GGTCGGAACA GGAGAGCGCA CGAGGGAGCT TCCAGGGGGA AACGCCTGGT
    16441 ATCTTTATAG TCCTGTCGGG TTTCGCCACC TCTGACTTGA GCGTCGATTT TTGTGATGCT
    16501 CGTCAGGGGG GCGGAGCCTA TGGAAAAACG CCAGCAACGC GGCCTTTTTA CGGTTCCTGG
    16561 CCTTTTGCTG GCCTTTTGCT CACATGTT
//
```

**Supplemental File 1E: *SagGD^vasa^***

```
LOCUS       SagGDvasa               17175 bp ds-DNA     circular
DEFINITION  synthetic circular DNA
ACCESSION   .
VERSION     .
KEYWORDS    .
SOURCE      synthetic DNA construct
  ORGANISM  synthetic DNA construct
REFERENCE   1  (bases 1 to 17175)
  AUTHORS   .
  TITLE     Direct Submission
  JOURNAL   Exported Wednesday, Dec 14, 2022 from SnapGene 5.2.5
            https://www.snapgene.com
FEATURES             Location/Qualifiers
     misc_feature    537..552
                     /label=M13F
                     /note="M13F"
     misc_feature    778..833
                     /label=attB 1
                     /note="attB 1"
     misc_feature    781..818
                     /label=minimal attb
                     /note="minimal attb"
     misc_feature    806..833
                     /label=EM1025
                     /note="EM1025"
     misc_feature    843..2587
                     /label=5' homology arm saglin
                     /note="5' homology arm saglin"
     misc_feature    2088..2587
                     /label=saglin ORF
                     /note="saglin ORF"
     misc_feature    2621..3073
                     /label=p609
                     /note="p609"
     misc_feature    complement(2621..2732)
                     /label=gRNA template
                     /note="gRNA template"
     misc_feature    2733..2752
                     /label="linker EM661/662"
                     /note="Saglin gRNA 3"
     misc_feature    complement(2753..3073)
                     /label=U6 promoter
                     /note="U6 promoter"
     misc_feature    3074..3526
                     /label=p610
                     /note="p610"
     misc_feature    complement(3074..3185)
                     /label=gRNA template
                     /note="gRNA template"
     misc_feature    3186..3205
                     /label="linker EM663/664"
                     /note="Saglin gRNA2"
     misc_feature    complement(3206..3526)
                     /label=U6 promoter
                     /note="U6 promoter"
     misc_feature    3527..3979
                     /label=p611
                     /note="p611"
     misc_feature    complement(3527..3634)
                     /label=gRNA template
                     /note="gRNA template"
     misc_feature    3635..3658
                     /label="Linker EM665/666"
                     /note="Saglin gRNA1"
     misc_feature    complement(3658..3979)
                     /label=U6 promoter
                     /note="U6 promoter"
     misc_feature    3980..3995
                     /label=EM723
                     /note="EM723"
     misc_feature    4014..4049
                     /label=Ag U6 terminator
                     /note="from Ag U6"
     misc_feature    4014..4042
                     /label=EM556
                     /note="EM556"
     misc_feature    complement(4047..4110)
                     /label=tracer
                     /note="tracer"
     misc_feature    complement(4050..4121)
                     /label=pX330 chimeric guide RNA scaffold
                     /note="pX330 chimeric guide RNA scaffold"
     misc_feature    complement(4115..4121)
                     /label=CRISPR  end
                     /note="CRISPR  end"
     misc_feature    complement(4122..4145)
                     /label=Lp sgRNA1 EM1096
                     /note="Lp gRNA3"
     misc_feature    complement(4146..4466)
                     /label=AGAP013557 U6 prom
                     /note="AGAP013557 U6 prom"
     misc_feature    complement(4169..4176)
                     /label=-30 TATA like
                     /note="-30 TATA like"
     misc_feature    complement(4198..4210)
                     /label=-65 conserved
                     /note="-65 conserved"
     misc_feature    4471..4502
                     /label=from Ag U6
                     /note="from Ag U6"
     misc_feature    complement(4500..4563)
                     /label=tracer
                     /note="tracer"
     misc_feature    complement(4503..4574)
                     /label=pX330 chimeric guide RNA scaffold
                     /note="pX330 chimeric guide RNA scaffold"
     misc_feature    complement(4568..4574)
                     /label=CRISPR  end
                     /note="CRISPR  end"
     misc_feature    4575..4598
                     /label=Lp sgRNA2 EM1098
                     /note="sgRNA2 EM1098"
     misc_feature    complement(4599..4919)
                     /label=AGAP013557 U6 prom
                     /note="AGAP013557 U6 prom"
     misc_feature    complement(4622..4629)
                     /label=-30 TATA like
                     /note="-30 TATA like"
     misc_feature    complement(4651..4663)
                     /label=-65 conserved
                     /note="-65 conserved"
     misc_feature    4924..4955
                     /label=Ag U6 terminator
                     /note="from Ag U6"
     misc_feature    complement(4953..5016)
                     /label=tracer
                     /note="tracer"
     misc_feature    complement(4956..5027)
                     /label=pX330 chimeric guide RNA scaffold
                     /note="pX330 chimeric guide RNA scaffold"
     misc_feature    complement(5021..5027)
                     /label=CRISPR  end
                     /note="CRISPR  end"
     misc_feature    5028..5051
                     /label=Lp sgRNA3 EM1100
                     /note="sgRNA3 EM1100"
     misc_feature    complement(5052..5372)
                     /label=AGAP013557 U6 prom
                     /note="AGAP013557 U6 prom"
     misc_feature    complement(5075..5082)
                     /label=-30 TATA like
                     /note="-30 TATA like"
     misc_feature    complement(5104..5116)
                     /label=-65 conserved
                     /note="-65 conserved"
     misc_feature    5396..6619
                     /label=3xP3 DsRedNLS sv40
                     /note="3xP3 DsRedNLS sv40"
     misc_feature    6623..8856
                     /label=vasa promoter
                     /note="vasa promoter"
     misc_feature    8872..8940
                     /label=3xFLAG
                     /note="3xFLAG"
     misc_feature    8941..8991
                     /label=NLS
                     /note="NLS"
     misc_feature    8992..13092
                     /label=Cas9
                     /note="Cas9"
     misc_feature    13093..13140
                     /label=NLS
                     /note="NLS"
     misc_feature    13150..13376
                     /label=SV40 term
                     /note="SV40 term"
     misc_feature    13409..15349
                     /label=Saglin 3' homology arm
                     /note="Saglin 3' homology arm"
     misc_feature    complement(15353..15408)
                     /label=attB2
                     /note="attB2"
     misc_feature    complement(15436..15454)
                     /label=M13R
                     /note="M13R"
     misc_feature    complement(15480..15498)
                     /label=pDONR-RP
                     /note="pDONR-RP"
     source          1..17175
                     /dnas_title=“knock-in plasmid”
ORIGIN
        1 ctttcctgcg ttatcccctg attctgtgga taaccgtatt accgcctttg agtgagctga
       61 taccgctcgc cgcagccgaa cgaccgagcg cagcgagtca gtgagcgagg aagcggaaga
      121 gcgcccaata cgcaaaccgc ctctccccgc gcgttggccg attcattaat gcagctggca
      181 cgacaggttt cccgactgga aagcgggcag tgagcgcaac gcaattaata cgcgtaccgc
      241 tagccaggaa gagtttgtag aaacgcaaaa aggccatccg tcaggatggc cttctgctta
      301 gtttgatgcc tggcagttta tggcgggcgt cctgcccgcc accctccggg ccgttgcttc
      361 acaacgttca aatccgctcc cggcggattt gtcctactca ggagagcgtt caccgacaaa
      421 caacagataa aacgaaaggc ccagtcttcc gactgagcct ttcgttttat ttgatgcctg
      481 gcagttccct actctcgcgt taacgctagc atggatgttt tcccagtcac gacgttgtaa
      541 aacgacggcc agtcttaagc tcgggcccct acaggtcact aataccatct aagtagttga
      601 ttcatagtga ctggatatgt tgtgttttac agtattatgt agtctgtttt ttatgcaaaa
      661 tctaatttaa tatattgata tttatatcat tttacgtttc tcgttcaact tttctataca
      721 aagttggtac cgggcccccc gctagcgtcg acggtatcga taagcttgat cggatcctgc
      781 gggtgccagg gcgtgccctt gggctccccg ggcgcgtact ccacctcacc cataggcctg
      841 ataacacact ggcacgacga tcatcccttt ctgctcgaaa cggtcgcgaa ggcgtacgag
      901 ccgctcaagg ccgcgctgga gtcggagcag gatgcgcatc gaagccttcg gctggtgccc
      961 gagctgcagc gcaagcagct catctacagc atcgaggcgg gccggctcga ggacgcccac
     1021 atcctgcaca tgacgctcaa gggccagtgg aagccggacc aaatcgtgtc ggccatccag
     1081 gacggctacc acgtcaaccc gacgatcatg gagcatctgc tcgagttcgt ccgcgccatc
     1141 ccggtgcgga aggagcgggc cgcctactac aaagcgatcg ggccggtcat tcgcaacttc
     1201 aagctgaccc tgacgtacgt gacgctcctg ttcgcgggcg atgcgaccgg cgtgtttgac
     1261 acgcgaaagg atcgcgacga ctacatctcg caaccgctca ccgtgttcgt gcacatgctg
     1321 cgctggcagc tggcgaacct cgagttcgag ttccacctgt cgctggcgga gcgcttcccc
     1381 cgctactact cgctgcacat cgagcaaatc ttcttctttg cgcccccgca ctggcagaag
     1441 gcggagaagc gccggctgtt cgagatcgcg agcctgttca aggcgaaggg gcaccggttt
     1501 gcggcgatcg agcagctgct gaagtgggcg caccagtacg caccgatgcg cggtggcaaa
     1561 gcccacctgg aggaaatact gcccacgctg gcgctcgaaa cggagaagct gcggcgcacg
     1621 gtcgagcaga cgggcaagag ctcagcggag ctggtgcggc tcaagaagct ggaggaaaag
     1681 ttcgtaagca aaaaggatcg ctggaacacg aacacctacc gcacgtacct gcacatgatc
     1741 aagacgaagg gtaagttcgg ccagcatagg atggaaatgg agcgcaaggg ttaaatagcg
     1801 ggaatggcac atgcgaggcg atggatgaaa tacaagtcgt ttggtgagcg gagtcaccaa
     1861 gcaagcataa ctttcatgcg tacaatgtgt atgatcactg atgacaaagc gcttgggtgg
     1921 ggcgcttgct gctgaagcgt ggctgacaaa tggggggggg gcttgaacaa atatttgcac
     1981 acacgataat gctcttgttt gccatgcgct gacggaagac aaatcgctgc aagcgacggc
     2041 tcaaacctgc tcaaaccagg gaaggtttgc ataaggttca cttataaatg agcgttcgtg
     2101 gctactcggg cgttcaagtc atctcgagca ggaagcaccg cagcatgtca cgtttgccca
     2161 ctgtactgtt gctcttagcg agcgccgcag tcctcgcggc tggcggccag gaagcaaccg
     2221 aagacccgtt cgcggacgaa accgaccagt gccagatcag cgttagtgcc gagacgatga
     2281 aatccctcca cggcgggtcg atgcagcccg acggcacgtg cgacaacctg tgggaaagct
     2341 ttctatctca gtttcaccaa gtcagggaaa acctgaccgc gtgccaggag cgggcagccg
     2401 ccggaccagc gcccgatcct tccagccagt tctgtcagca gctgctggac gatgcgcagc
     2461 ggcagatgga gcaggagcat cgccagtacg ccgccaccct ggaggagcag ctgcacgcag
     2521 cgcagcagga aacccagcag gagcaggaga tgaagaaggc gctgcagaag cagctcgacg
     2581 cgctcacgta agcgtcgacg gtatcgataa gcttgatcgg atccgatgaa aataagaaaa
     2641 acatttgaca aaaaaagcac cgactcggtg ccactttttc aagttgataa cggactagcc
     2701 ttattttaac ttgctatttc tagctctaaa accctgcttg aagttaccgt tcaaggacga
     2761 ggggaaaaaa ggttgtatat atactttgcg ctttcaatcc ttgctctagc gatgcataaa
     2821 ggatattcaa gaaggatttt ttgcctgagt gtgactgtta taaggttttg atcctgccgt
     2881 aactcacgct aatgtgatgt ttcataaact tttcaacatt tcttgtattg cttcatctct
     2941 tttttaatca aaagttacta ttcattgtaa tacgttgaaa tatcaaaaaa gaataaacag
     3001 ttttcctatg attaattcaa aaacagagtt ttctagaata accaagagcc aaaccgcttc
     3061 agtgtatatg tggggaagat gaaaataaga aaaacatttg acaaaaaaag caccgactcg
     3121 gtgccacttt ttcaagttga taacggacta gccttatttt aacttgctat ttctagctct
     3181 aaaaccaacg cgctgtacat cgatcaagga cgaggggaaa aaaggttgta tatatacttt
     3241 gcgctttcaa tccttgctct agcgatgcat aaaggatatt caagaaggat tttttgcctg
     3301 agtgtgactg ttataaggtt ttgatcctgc cgtaactcac gctaatgtga tgtttcataa
     3361 acttttcaac atttcttgta ttgcttcatc tcttttttaa tcaaaagtta ctattcattg
     3421 taatacgttg aaatatcaaa aaagaataaa cagttttcct atgattaatt caaaaacaga
     3481 gttttctaga ataaccaaga gccaaaccgc ttcagtgtat atgtggagag gatgaaaata
     3541 agaaaaacat ttgacaaaaa aagcaccgac tcggtgccac tttttcaagt tgataacgga
     3601 ctagccttat tttaacttgc tatttctagc tctaaaacct cgtactacaa cctgatgcaa
     3661 ggacgagggg aaaaaaggtt gtatatatac tttgcgcttt caatccttgc tctagcgatg
     3721 cataaaggat attcaagaag gattttttgc ctgagtgtga ctgttataag gttttgatcc
     3781 tgccgtaact cacgctaatg tgatgtttca taaacttttc aacatttctt gtattgcttc
     3841 atctcttttt taatcaaaag ttactattca ttgtaatacg ttgaaatatc aaaaaagaat
     3901 aaacagtttt cctatgatta attcaaaaac agagttttct agaataacca agagccaaac
     3961 cgcttcagtg tatatgtgga acacactggc acgacatcga taagcttgat cggatccgat
     4021 gaaaataaga aaaacatttg acaaaaaaag caccgactcg gtgccacttt ttcaagttga
     4081 taacggacta gccttatttt aacttgctat ttctagctct aaaacggtgt cgctagtgct
     4141 gattcaagga cgaggggaaa aaaggttgta tatatacttt gcgctttcaa tccttgctct
     4201 agcgatgcat aaaggatatt caagaaggat tttttgcctg agtgtgactg ttataaggtt
     4261 ttgatcctgc cgtaactcac gctaatgtga tgtttcataa acttttcaac atttcttgta
     4321 ttgcttcatc tcttttttaa tcaaaagtta ctattcattg taatacgttg aaatatcaaa
     4381 aaagaataaa cagttttcct atgattaatt caaaaacaga gttttctaga ataaccaaga
     4441 gccaaaccgc ttcagtgtat atgtggggaa gatgaaaata agaaaaacat ttgacaaaaa
     4501 aagcaccgac tcggtgccac tttttcaagt tgataacgga ctagccttat tttaacttgc
     4561 tatttctagc tctaaaactc cctgacgata tttcacgcaa ggacgagggg aaaaaaggtt
     4621 gtatatatac tttgcgcttt caatccttgc tctagcgatg cataaaggat attcaagaag
     4681 gattttttgc ctgagtgtga ctgttataag gttttgatcc tgccgtaact cacgctaatg
     4741 tgatgtttca taaacttttc aacatttctt gtattgcttc atctcttttt taatcaaaag
     4801 ttactattca ttgtaatacg ttgaaatatc aaaaaagaat aaacagtttt cctatgatta
     4861 attcaaaaac agagttttct agaataacca agagccaaac cgcttcagtg tatatgtgga
     4921 gaggatgaaa ataagaaaaa catttgacaa aaaaagcacc gactcggtgc cactttttca
     4981 agttgataac ggactagcct tattttaact tgctatttct agctctaaaa cccgaggacc
     5041 cacatcgttc caaggacgag gggaaaaaag gttgtatata tactttgcgc tttcaatcct
     5101 tgctctagcg atgcataaag gatattcaag aaggattttt tgcctgagtg tgactgttat
     5161 aaggttttga tcctgccgta actcacgcta atgtgatgtt tcataaactt ttcaacattt
     5221 cttgtattgc ttcatctctt ttttaatcaa aagttactat tcattgtaat acgttgaaat
     5281 atcaaaaaag aataaacagt tttcctatga ttaattcaaa aacagagttt tctagaataa
     5341 ccaagagcca aaccgcttca gtgtatatgt ggataaaccc acatcgttcc gtatctaatt
     5401 caattagaga ctaattcaat tagagctaat tcaattagga tccaagctta tcgatttcga
     5461 accctcgacc gccggagtat aaatagaggc gcttcgtcta cggagcgaca attcaattca
     5521 aacaagcaaa gtgaacacgt cgctaagcga aagctaagca aataaacaag cgcagctgaa
     5581 caagctaaac aatcggggta ccgctagagt cgacggtacc gcgggcccgg gatccaccgg
     5641 tcgccaccat ggcctcctcc gagaacgtca tcaccgagtt catgcgcttc aaggtgcgca
     5701 tggagggcac cgtgaacggc cacgagttcg agatcgaggg cgagggcgag ggccgcccct
     5761 acgagggcca caacaccgtg aagctgaagg tgaccaaggg cggccccctg cccttcgcct
     5821 gggacatcct gtccccccag ttccagtacg gctccaaggt gtacgtgaag caccccgccg
     5881 acatccccga ctacaagaag ctgtccttcc ccgagggctt caagtgggag cgcgtgatga
     5941 acttcgagga cggcggcgtg gcgaccgtga cccaggactc ctccctgcag gacggctgct
     6001 tcatctacaa ggtgaagttc atcggcgtga acttcccctc cgacggcccc gtgatgcaga
     6061 agaagaccat gggctgggag gcctccaccg agcgcctgta cccccgcgac ggcgtgctga
     6121 agggcgagac gcacaaggcc ctgaagctga aggacggcgg ccactacctg gtggagttca
     6181 agtccatcta catggccaag aagcccgtgc agctgcccgg ctactactac gtggacgcca
     6241 agctggacat cacctcccac aacgaggact acaccatcgt ggagcagtac gagcgcaccg
     6301 agggccgcca ccacctgttc ctgagatctc gacccaagaa aaagcggaag gtggaggacc
     6361 cgtaagatcc accggatcta gataactgat cataatcagc cataccacat ttgtagaggt
     6421 tttacttgct ttaaaaaacc tcccacacct ccccctgaac ctgaaacata aaatgaatgc
     6481 aattgttgtt gttaacttgt ttattgcagc ttataatggt tacaaataaa gcaatagcat
     6541 cacaaatttc acaaataaag catttttttc actgcattct agttgtggtt tgtccaaact
     6601 catcaatgta tcttattatc cgatcccgat gtagaacgcg agcaaattct tttccttcca
     6661 tgacagcagc agctacagtg ggaagccgaa cgtcagacgt gtttgacatg ccgaactggg
     6721 cgggaaaatt acagcgtgcg ctttgttttc aagcaaatca caactcgctg caaacaaaac
     6781 cgttgagaaa ttgattgttt tataatttgt attgtatttt atttgttata ataaactaaa
     6841 aagacatact ttttgcatat tttatacata aaaacataca tgcagcatta taaaacacat
     6901 ataaaccctc cctgtagagt cccgtatcga aatcttccat cctagttgca cagtacgacg
     6961 gacgagtagg ccgtgtccgt gcaaattcca gcttttagca gtcttttgct cggagcactc
     7021 gcggcgagtc ggaggtttct gctgaggtgc ttagcgctaa attagccaat tgcttttgca
     7081 agtgaaataa ccagccgaat agtacttcaa aactcaggta agtgaactag ttttatagaa
     7141 caaatgtttg tttgttagaa gttagtgaag tgtttgtgaa aaaaatctct catttcggca
     7201 aaactaacgt aactgatttc aaattgaatt attgttttgt gatgttatat tatttcatcc
     7261 agttgattag tattttctta gttatgttca aaatacagtt aaattaaatt tcatttcatt
     7321 tactcataaa ataatctctt ggcttattta atttttctcg aattcgcttg tattgttcag
     7381 tagcacgcgc cattcgccct ttgtttcatt ttgtacctgc tcccactaac acactggcag
     7441 tgcgaaacaa aagccttcgc acgcgttgct ggtattagag tgtgtgcgtg tgtgtgttga
     7501 gcgctctgtc aaaatcggct gttgccgccg gtaccgaaat tgcctgttcg cacgctgttc
     7561 gtaaacattc cgtggtgtgt atcgtgtgtt gtgcatgttg cgcgcctccc cccttttgat
     7621 agcaggctgc cgtggctgcc gtggtgtgtg gcgcagttga gtttttggat taattttcta
     7681 aggaaatggc acgagaagag cggtggcagt gtgttggttt gctctgtccc ttcctttctg
     7741 tgtgaagtgt tcttacagca cagcacgtat ccaccaccgc acacagagca ggcaaggaag
     7801 tggaagtgaa caagtgtgct gcgcatgcat gtgtgtgggg ggcattttag ctgagatcgt
     7861 cgttatttga gaagcggtat aggggccagt cggtgtcgac gtacggaagc ggtttagttt
     7921 taatccaagc gtatcccgtc gtggagtggt tgtgtggctc tgtgtgctct catatcagtt
     7981 ccagagtgag gttagtagaa tcacagtcct tggccttttt cgttacaaga tatccagaag
     8041 gatggcgtta tttccacagc ttaccatggt gctcttgttt gctcgaatca ggggagaaaa
     8101 acagtttcgt gtttcatgaa ccgcagttgg cactggagcg gattcaaaag tcttcgatat
     8161 gcaatagata agagagtcgt tggggcatag ttgggaagcc tttccgagat gtggagtttc
     8221 cgagaggaga aatggtgctt tcgtgcacgt tccgggacag cgggccccgc gaagagcatc
     8281 tcgttgtcgt tcatccggca ataattgatg cgaaaagcgc gcgcgccact ggcttagcgc
     8341 agtgtacaca gtgatattca cctacacaca cagaggcaca cgccttcaca cgcgcgcgtg
     8401 cttcaaaggc tacttcggtg gcggtgtgtg aggtcgcttg caatggacaa tgaaaatttc
     8461 gctggaaaat accatcgtct ctttaggttg caatgggtgc gggtagagcg gtggtcgtcg
     8521 atattggtgg tgtagtgtgt gtgtgtgtgt gtgtgtgtgt gtgtgtgtgt gtgtgtgtgt
     8581 gtgtgtgtgt gtgtgtgtgt gtgtgtgtgt gtgtgtgtgt gtgtgtgtgt gcaacggcaa
     8641 ttattttttg taatatttcg accatctttc tttctctctc tccacgtgct gctgctgttg
     8701 ctgctgctgc tgcattgcat gttccactat tcctctcggt ttgtgcctgc ggacgccatt
     8761 gctagtcgaa agagagtcgc cgttagtcgc gcttcgagca acggacacgt tttttggttg
     8821 aaaccaacag cttttttcat cttcgggaga cacacagatc ccggtgccac catggactat
     8881 aaggaccacg acggagacta caaggatcat gatattgatt acaaagacga tgacgataag
     8941 atggccccaa agaagaagcg gaaggtcggt atccacggag tcccagcagc cgacaagaag
     9001 tacagcatcg gcctggacat cggcaccaac tctgtgggct gggccgtgat caccgacgag
     9061 tacaaggtgc ccagcaagaa attcaaggtg ctgggcaaca ccgaccggca cagcatcaag
     9121 aagaacctga tcggagccct gctgttcgac agcggcgaaa cagccgaggc cacccggctg
     9181 aagagaaccg ccagaagaag atacaccaga cggaagaacc ggatctgcta tctgcaagag
     9241 atcttcagca acgagatggc caaggtggac gacagcttct tccacagact ggaagagtcc
     9301 ttcctggtgg aagaggataa gaagcacgag cggcacccca tcttcggcaa catcgtggac
     9361 gaggtggcct accacgagaa gtaccccacc atctaccacc tgagaaagaa actggtggac
     9421 agcaccgaca aggccgacct gcggctgatc tatctggccc tggcccacat gatcaagttc
     9481 cggggccact tcctgatcga gggcgacctg aaccccgaca acagcgacgt ggacaagctg
     9541 ttcatccagc tggtgcagac ctacaaccag ctgttcgagg aaaaccccat caacgccagc
     9601 ggcgtggacg ccaaggccat cctgtctgcc agactgagca agagcagacg gctggaaaat
     9661 ctgatcgccc agctgcccgg cgagaagaag aatggcctgt tcggaaacct gattgccctg
     9721 agcctgggcc tgacccccaa cttcaagagc aacttcgacc tggccgagga tgccaaactg
     9781 cagctgagca aggacaccta cgacgacgac ctggacaacc tgctggccca gatcggcgac
     9841 cagtacgccg acctgtttct ggccgccaag aacctgtccg acgccatcct gctgagcgac
     9901 atcctgagag tgaacaccga gatcaccaag gcccccctga gcgcctctat gatcaagaga
     9961 tacgacgagc accaccagga cctgaccctg ctgaaagctc tcgtgcggca gcagctgcct
    10021 gagaagtaca aagagatttt cttcgaccag agcaagaacg gctacgccgg ctacattgac
    10081 ggcggagcca gccaggaaga gttctacaag ttcatcaagc ccatcctgga aaagatggac
    10141 ggcaccgagg aactgctcgt gaagctgaac agagaggacc tgctgcggaa gcagcggacc
    10201 ttcgacaacg gcagcatccc ccaccagatc cacctgggag agctgcacgc cattctgcgg
    10261 cggcaggaag atttttaccc attcctgaag gacaaccggg aaaagatcga gaagatcctg
    10321 accttccgca tcccctacta cgtgggccct ctggccaggg gaaacagcag attcgcctgg
    10381 atgaccagaa agagcgagga aaccatcacc ccctggaact tcgaggaagt ggtggacaag
    10441 ggcgcttccg cccagagctt catcgagcgg atgaccaact tcgataagaa cctgcccaac
    10501 gagaaggtgc tgcccaagca cagcctgctg tacgagtact tcaccgtgta taacgagctg
    10561 accaaagtga aatacgtgac cgagggaatg agaaagcccg ccttcctgag cggcgagcag
    10621 aaaaaggcca tcgtggacct gctgttcaag accaaccgga aagtgaccgt gaagcagctg
    10681 aaagaggact acttcaagaa aatcgagtgc ttcgactccg tggaaatctc cggcgtggaa
    10741 gatcggttca acgcctccct gggcacatac cacgatctgc tgaaaattat caaggacaag
    10801 gacttcctgg acaatgagga aaacgaggac attctggaag atatcgtgct gaccctgaca
    10861 ctgtttgagg acagagagat gatcgaggaa cggctgaaaa cctatgccca cctgttcgac
    10921 gacaaagtga tgaagcagct gaagcggcgg agatacaccg gctggggcag gctgagccgg
    10981 aagctgatca acggcatccg ggacaagcag tccggcaaga caatcctgga tttcctgaag
    11041 tccgacggct tcgccaacag aaacttcatg cagctgatcc acgacgacag cctgaccttt
    11101 aaagaggaca tccagaaagc ccaggtgtcc ggccagggcg atagcctgca cgagcacatt
    11161 gccaatctgg ccggcagccc cgccattaag aagggcatcc tgcagacagt gaaggtggtg
    11221 gacgagctcg tgaaagtgat gggccggcac aagcccgaga acatcgtgat cgaaatggcc
    11281 agagagaacc agaccaccca gaagggacag aagaacagcc gcgagagaat gaagcggatc
    11341 gaagagggca tcaaagagct gggcagccag atcctgaaag aacaccccgt ggaaaacacc
    11401 cagctgcaga acgagaagct gtacctgtac tacctgcaga atgggcggga tatgtacgtg
    11461 gaccaggaac tggacatcaa ccggctgtcc gactacgatg tggaccatat cgtgcctcag
    11521 agctttctga aggacgactc catcgacaac aaggtgctga ccagaagcga caagaaccgg
    11581 ggcaagagcg acaacgtgcc ctccgaagag gtcgtgaaga agatgaagaa ctactggcgg
    11641 cagctgctga acgccaagct gattacccag agaaagttcg acaatctgac caaggccgag
    11701 agaggcggcc tgagcgaact ggataaggcc ggcttcatca agagacagct ggtggaaacc
    11761 cggcagatca caaagcacgt ggcacagatc ctggactccc ggatgaacac taagtacgac
    11821 gagaatgaca agctgatccg ggaagtgaaa gtgatcaccc tgaagtccaa gctggtgtcc
    11881 gatttccgga aggatttcca gttttacaaa gtgcgcgaga tcaacaacta ccaccacgcc
    11941 cacgacgcct acctgaacgc cgtcgtggga accgccctga tcaaaaagta ccctaagctg
    12001 gaaagcgagt tcgtgtacgg cgactacaag gtgtacgacg tgcggaagat gatcgccaag
    12061 agcgagcagg aaatcggcaa ggctaccgcc aagtacttct tctacagcaa catcatgaac
    12121 tttttcaaga ccgagattac cctggccaac ggcgagatcc ggaagcggcc tctgatcgag
    12181 acaaacggcg aaaccgggga gatcgtgtgg gataagggcc gggattttgc caccgtgcgg
    12241 aaagtgctga gcatgcccca agtgaatatc gtgaaaaaga ccgaggtgca gacaggcggc
    12301 ttcagcaaag agtctatcct gcccaagagg aacagcgata agctgatcgc cagaaagaag
    12361 gactgggacc ctaagaagta cggcggcttc gacagcccca ccgtggccta ttctgtgctg
    12421 gtggtggcca aagtggaaaa gggcaagtcc aagaaactga agagtgtgaa agagctgctg
    12481 gggatcacca tcatggaaag aagcagcttc gagaagaatc ccatcgactt tctggaagcc
    12541 aagggctaca aagaagtgaa aaaggacctg atcatcaagc tgcctaagta ctccctgttc
    12601 gagctggaaa acggccggaa gagaatgctg gcctctgccg gcgaactgca gaagggaaac
    12661 gaactggccc tgccctccaa atatgtgaac ttcctgtacc tggccagcca ctatgagaag
    12721 ctgaagggct cccccgagga taatgagcag aaacagctgt ttgtggaaca gcacaagcac
    12781 tacctggacg agatcatcga gcagatcagc gagttctcca agagagtgat cctggccgac
    12841 gctaatctgg acaaagtgct gtccgcctac aacaagcacc gggataagcc catcagagag
    12901 caggccgaga atatcatcca cctgtttacc ctgaccaatc tgggagcccc tgccgccttc
    12961 aagtactttg acaccaccat cgaccggaag aggtacacca gcaccaaaga ggtgctggac
    13021 gccaccctga tccaccagag catcaccggc ctgtacgaga cacggatcga cctgtctcag
    13081 ctgggaggcg acaaaaggcc ggcggccacg aaaaaggccg gccaggcaaa aaagaaaaag
    13141 taagaattct agacataatc agccatacca catttgtaga ggttttactt gctttaaaaa
    13201 acctcccaca cctccccctg aacctgaaac ataaaatgaa tgcaattgtt gttgttaact
    13261 tgtttattgc agcttataat ggttacaaat aaagcaatag catcacaaat ttcacaaata
    13321 aagcattttt cttcactgca ttctagttgt ggtttgtcca aactcatcaa tgtatctcga
    13381 cgatgtaggt cacagtctcg aagccgcgac gcagctgcaa accagcctcg ccacgatcga
    13441 gctgaagcac tggcgcaagt tcgaccggtt cgtaccgtac gctaacgcgc tgccccagcc
    13501 agcccagcgg ctggaggtgc tgcgtgtgct gctgagccag atcggcgacc gggagaagaa
    13561 gacttcccac aaatatctgg taaaggcggc acggcagttt gacatttgcg agcagtttat
    13621 cggccggggc aaggtggacc aggcggtcaa gaagcagctg gaggaactgc gcgggaagtt
    13681 tgcaacgttt gcgaagggca agaactacca gcactatctg agcgagtcgc gcaagtcgtc
    13741 cggctaggcg tcggcgacgg tctaaaagtg attcaaagcc ccgtaggcga ggaacccccg
    13801 tacgtgtcgt gtagcgcaaa cccttcacca tcatagcaca gtgaaataaa gcaagaataa
    13861 aaattaaaaa aaaaaacaat tgtaatcgaa atattgagca acagcaaagc aaagcctttc
    13921 ctttggatca tccgcaatcg gcacgttgcg agttgggcgg cagcgttgct tgtgcaatcg
    13981 ctggacccat gctggacaaa cagatgagca cactggcctt cccagccagc taatattgac
    14041 acttcattcg agctggacca ttaatcacgg cgtctgtttg cctttgggcc aagtataact
    14101 agcaccacgc accgaacaga cgggctctag ttttgtttct cttgctgtga gcgtcagtta
    14161 attaaaaaaa caaacctatg gatcacagca gagtgtggtg ctgtctgctt agcttcatcg
    14221 caatggccgt gctgtgtagc gcatccagtg cagcatgcac gctccaggtc cccgaaacca
    14281 tgataacgcg cctgctgcaa gcggatcagc ccgccggcgg ctgcgcagct gcttgggata
    14341 gtctactgtc tcggctggag caaatgcgcc acaatcttac gggctgcagc gagcgcgaaa
    14401 cgacgaaccc ggcaccaccc gatgataccc gcaccaacgc cccctgtcag ctgctgctga
    14461 acgagctgca acggaaaggg aatcaaaccc tgctccagct cgacaaagaa atggccagca
    14521 aagcactgta cgagcgcatc gagctggcca aggtggtgca ggaccaggac cgcatccaac
    14581 aagcgatcga aacgatgcgc accgagcagc gaaaccaccg gcagcagctg ctactcctct
    14641 acctcgatgc ggcggaccta cgacgagccc tgcaccagta taagctgctc gtcgcgcagg
    14701 gcgatcgcca ccttccccag cagatagtga agttcgtgta cgccgcccca cggcacgaga
    14761 accgccggct cgagaacctg ctcgacctgg tgcggcagct gccggcgcgc caggaccagc
    14821 gcaccctgta ccagctgctg cagccggaaa ttatgaagcg ccccacgcag aaccagagca
    14881 ccctcgccat gctgacagcg ctcgagatgg gccaggtcgt ggaggggaac ggcgagctga
    14941 aaaagcagca ggacgcgatg taccagctgg tgctgaagcg ctggatgttt ctgtgcctcg
    15001 ccggccagta tcgcgagata gtgcagtttg ccacgaagca cccgcgcctg ttcgagcaga
    15061 tcggggcgaa gatcgccacg attcatccga agtactggtg gaagtttagc ttcacccagt
    15121 tcgtgacgta cccgaacctg ttgccgctgc ccgagcagcg gctggaagcg ttccgcacca
    15181 tcatgaagca gctgaagcag cgcaatggga agtttttcgc cgaccatctg gcgaagctgg
    15241 caccccagat cgagcgctgc gagcagtttc tgcggcagca gaagaaggag ctggagtgga
    15301 aggacgagct ggtgaagctg aagggacagt ttgccgactt tgaccgctta tcatgggtga
    15361 ggtggagtac gcgcccgggg agcccaaggg cacgccctgg cacccgcagc ttatccccta
    15421 tagtgagtcg tattacatgg tcatagctgt ttcctggcag ctctggcccg tgtctcaaaa
    15481 tctctgatgt tacattgcac aagataaaaa tatatcatca tgaacaataa aactgtctgc
    15541 ttacataaac agtaatacaa ggggtgttat gagccatatt caacgggaaa cgtcgaggcc
    15601 gcgattaaat tccaacatgg atgctgattt atatgggtat aaatgggctc gcgataatgt
    15661 cgggcaatca ggtgcgacaa tctatcgctt gtatgggaag cccgatgcgc cagagttgtt
    15721 tctgaaacat ggcaaaggta gcgttgccaa tgatgttaca gatgagatgg tcagactaaa
    15781 ctggctgacg gaatttatgc ctcttccgac catcaagcat tttatccgta ctcctgatga
    15841 tgcatggtta ctcaccactg cgatccccgg aaaaacagca ttccaggtat tagaagaata
    15901 tcctgattca ggtgaaaata ttgttgatgc gctggcagtg ttcctgcgcc ggttgcattc
    15961 gattcctgtt tgtaattgtc cttttaacag cgatcgcgta tttcgtctcg ctcaggcgca
    16021 atcacgaatg aataacggtt tggttgatgc gagtgatttt gatgacgagc gtaatggctg
    16081 gcctgttgaa caagtctgga aagaaatgca taaacttttg ccattctcac cggattcagt
    16141 cgtcactcat ggtgatttct cacttgataa ccttattttt gacgagggga aattaatagg
    16201 ttgtattgat gttggacgag tcggaatcgc agaccgatac caggatcttg ccatcctatg
    16261 gaactgcctc ggtgagtttt ctccttcatt acagaaacgg ctttttcaaa aatatggtat
    16321 tgataatcct gatatgaata aattgcagtt tcatttgatg ctcgatgagt ttttctaatc
    16381 agaattggtt aattggttgt aacactggca gagcattacg ctgacttgac gggacggcgc
    16441 aagctcatga ccaaaatccc ttaacgtgag ttacgcgtcg ttccactgag cgtcagaccc
    16501 cgtagaaaag atcaaaggat cttcttgaga tccttttttt ctgcgcgtaa tctgctgctt
    16561 gcaaacaaaa aaaccaccgc taccagcggt ggtttgtttg ccggatcaag agctaccaac
    16621 tctttttccg aaggtaactg gcttcagcag agcgcagata ccaaatactg ttcttctagt
    16681 gtagccgtag ttaggccacc acttcaagaa ctctgtagca ccgcctacat acctcgctct
    16741 gctaatcctg ttaccagtgg ctgctgccag tggcgataag tcgtgtctta ccgggttgga
    16801 ctcaagacga tagttaccgg ataaggcgca gcggtcgggc tgaacggggg gttcgtgcac
    16861 acagcccagc ttggagcgaa cgacctacac cgaactgaga tacctacagc gtgagctatg
    16921 agaaagcgcc acgcttcccg aagggagaaa ggcggacagg tatccggtaa gcggcagggt
    16981 cggaacagga gagcgcacga gggagcttcc agggggaaac gcctggtatc tttatagtcc
    17041 tgtcgggttt cgccacctct gacttgagcg tcgatttttg tgatgctcgt caggggggcg
    17101 gagcctatgg aaaaacgcca gcaacgcggc ctttttacgg ttcctggcct tttgctggcc
    17161 ttttgctcac atgtt
//
```
