## Supplemental File 2 for "A population modification gene drive targeting both *Saglin* and *Lipophorin* impairs *Plasmodium* transmission in *Anopheles* mosquitoes"

### Inheritance dynamics of the two transgenes *SagGDvasa* (DsRed) and *Lp::Sc2A10* (GFP) in 8 mosquito populations, tracked by COPAS flow cytometry

#### Typical COPAS analysis :

Objects separated according to size and opacity

Objects from first gate separated according to fluorescence

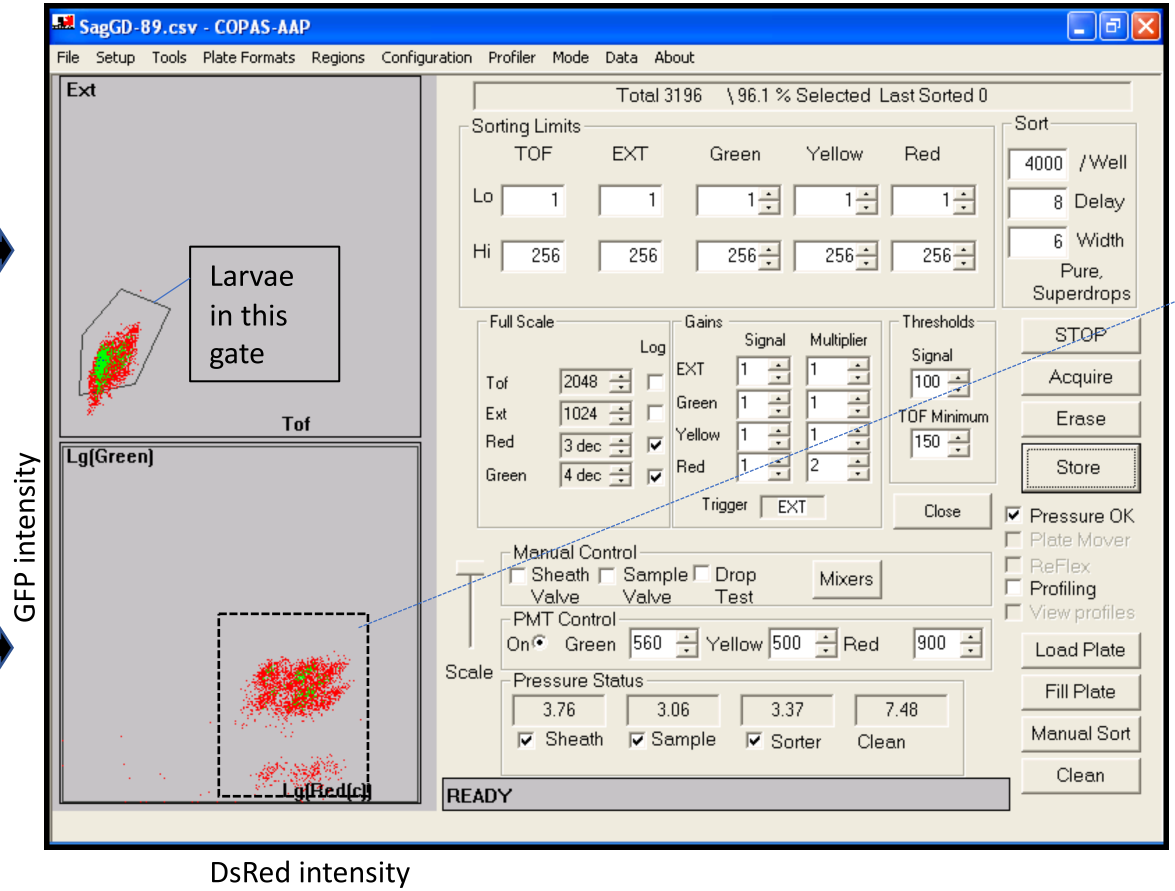


Extract screenshots of region with larvae

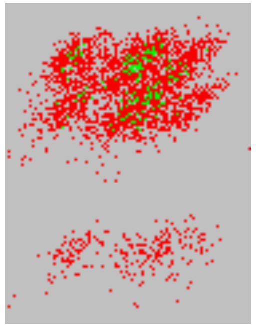

Interpretation of fluorescence :

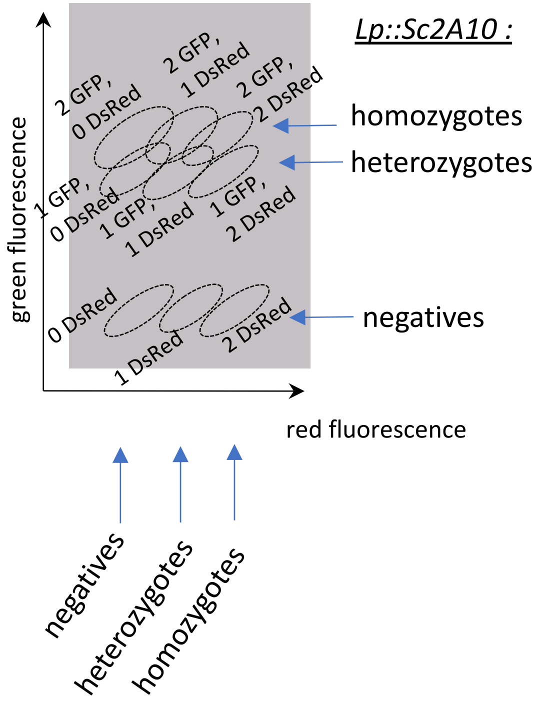

Population 1 : G0 = ♂ $\frac{SagGD^{vasa}}{Y}$ ; $\frac{Lp::Sc2A10}{+}$ x ♀ WT

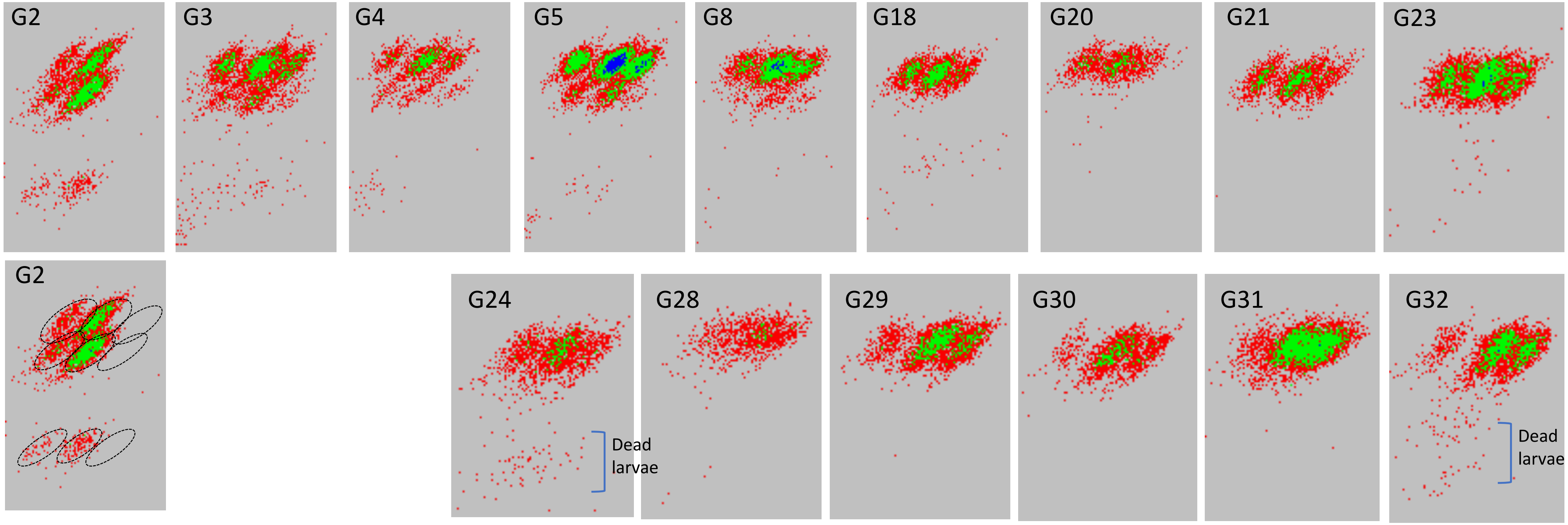

Population 2 : G0 = ♂ $\frac{SagGD^{vasa}}{Y}$ ; $\frac{Lp::Sc2A10}{+}$ x ♀ WT

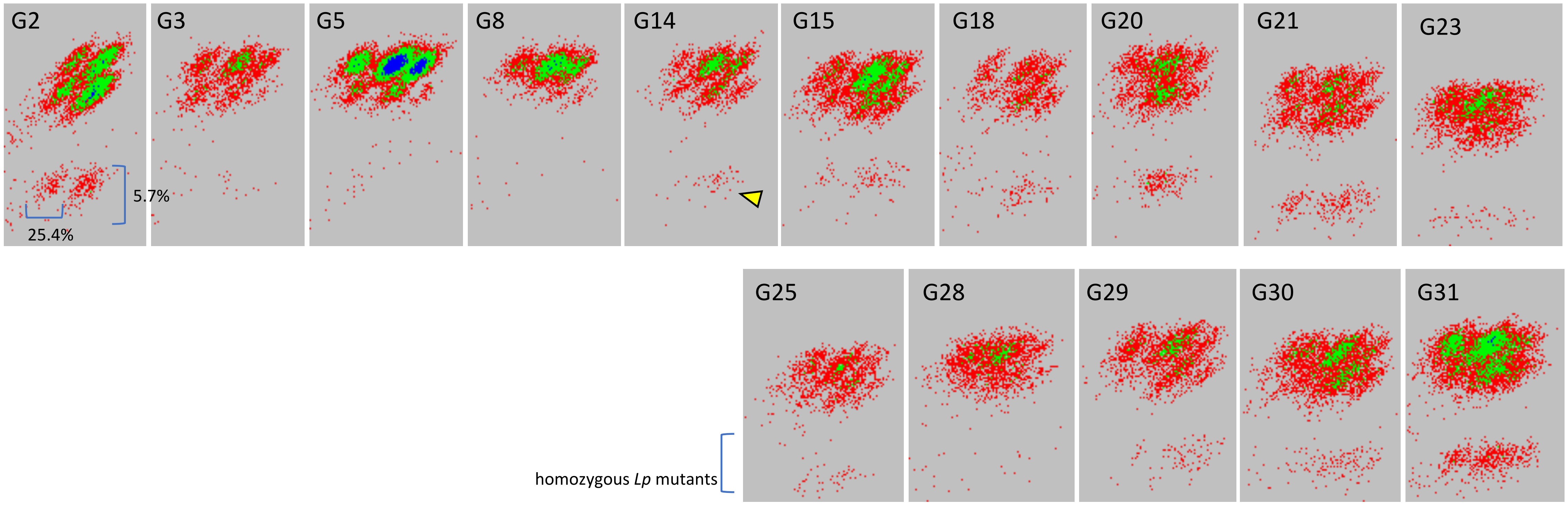

Population 3 : G0 = ~ 1:3 mix of G10 individuals from Population 1 : WT

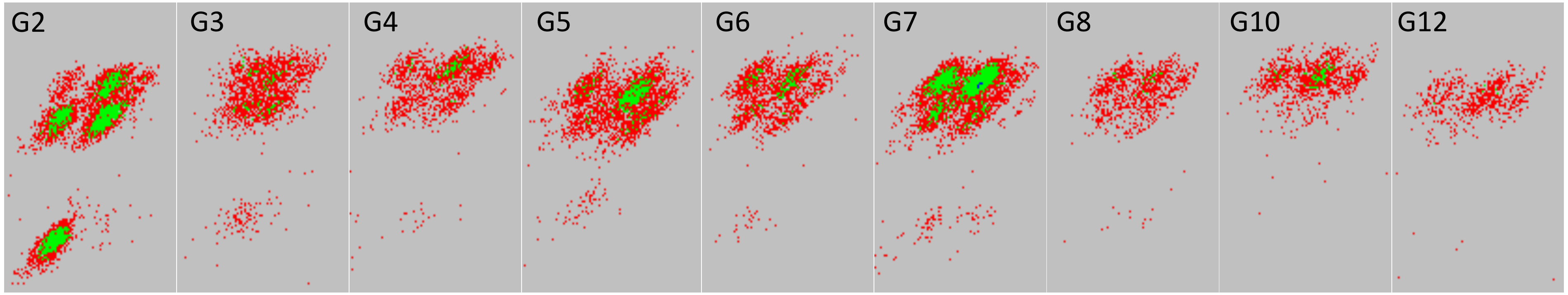


Population 4 :  G0 = ~ 1:2 mix G10 individuals from Population 1 : WT

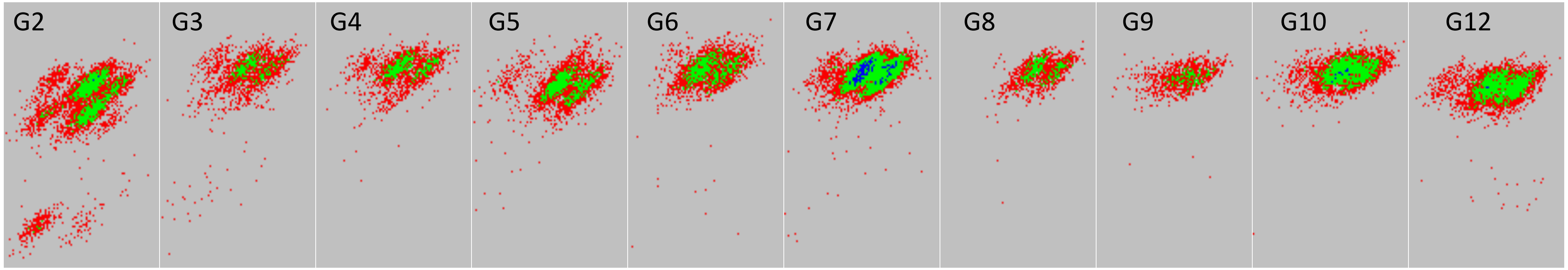

Population 5 : G0 = ♂ $\frac{SagGD^{vasa}}{Y}$ ; $\frac{Lp::Sc2A10}{Lp::Sc2A10}$ x ♀ WT

(from G16 of Population 1)

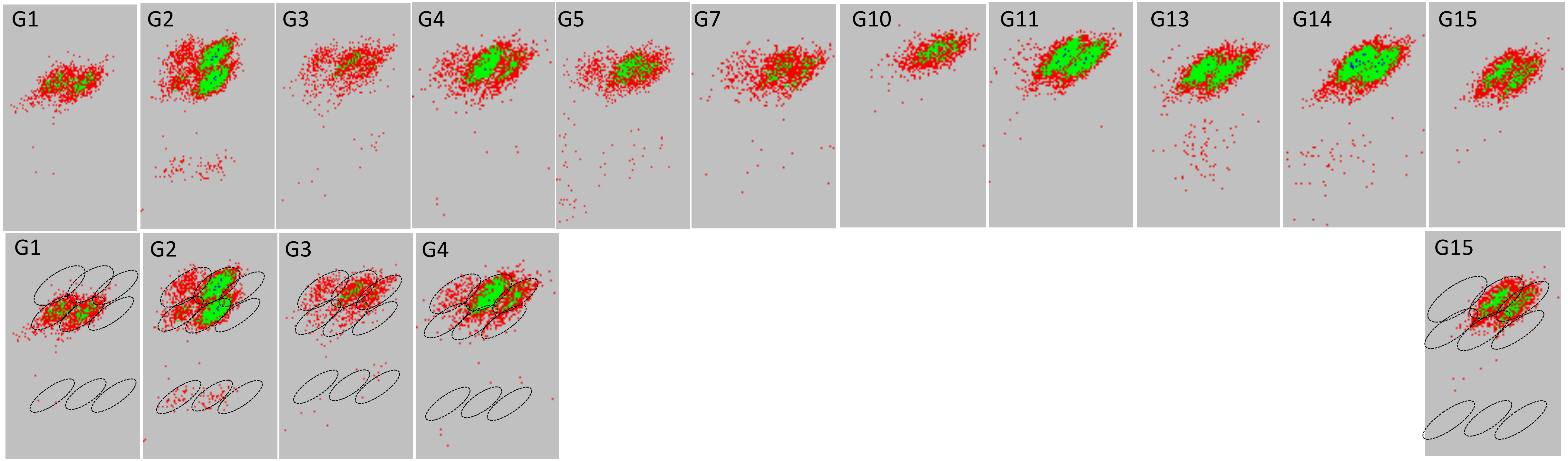

Population 6 : G0 = ♀ $\frac{SagGD^{vasa}}{SagGD^{vasa}}$ ; $\frac{Lp::Sc2A10}{Lp::Sc2A10}$ x ♂ WT

(from G16 of Population 1)

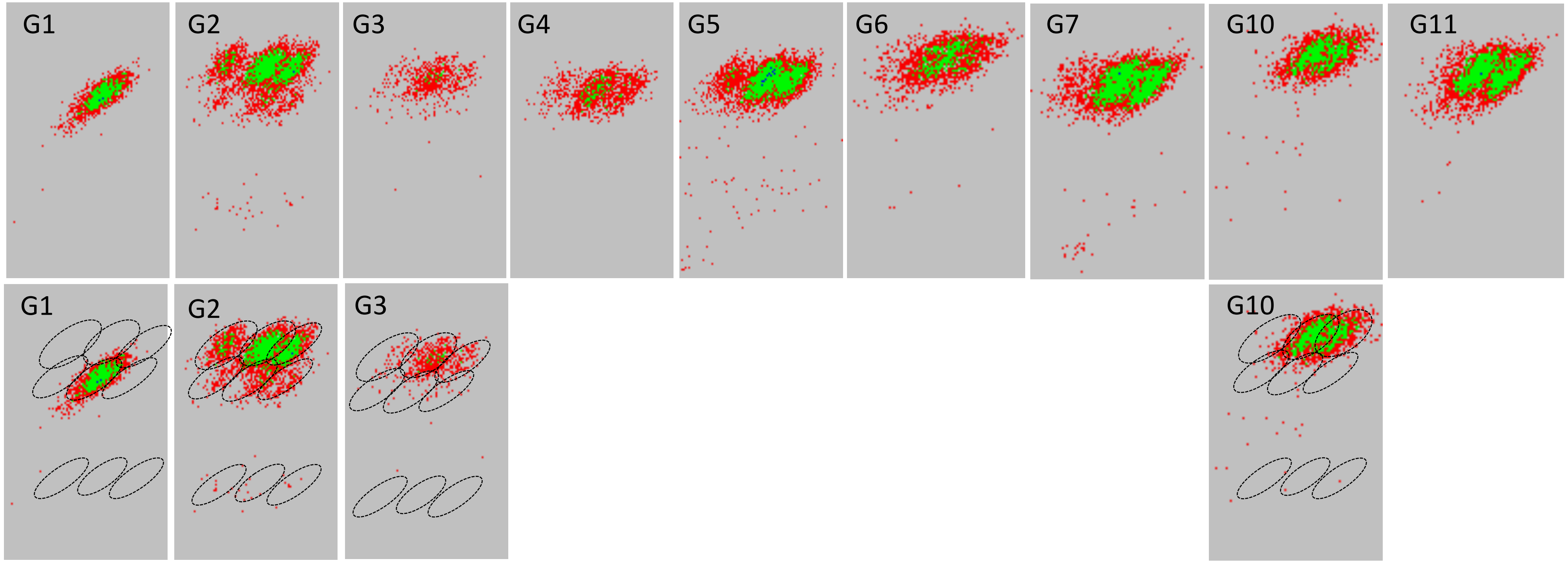

Population 7 : G0 = ♂ $\frac{SagGD^{vasa}}{Y}$ ; $\frac{Lp::Sc2A10}{+}$ x ♀ WT

(from G1 of Population 5)

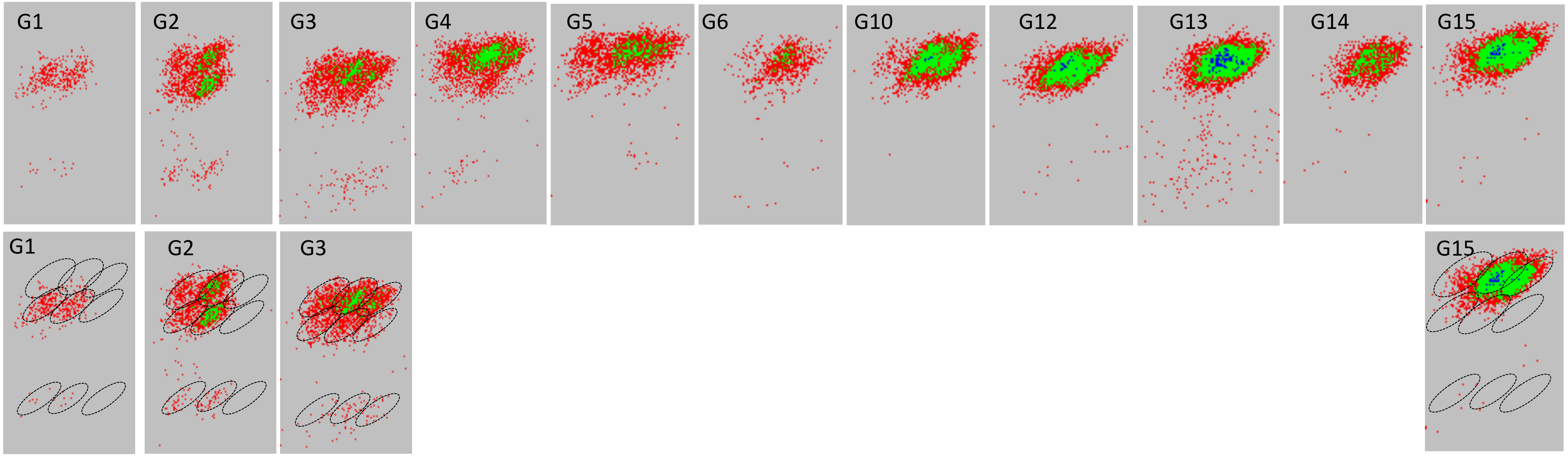

Population 8 : G0 = ♀ $\frac{SagGD^{vasa}}{+}$ ; $\frac{Lp::Sc2A10}{+}$ x ♂ WT

(from G1 of Population 5)

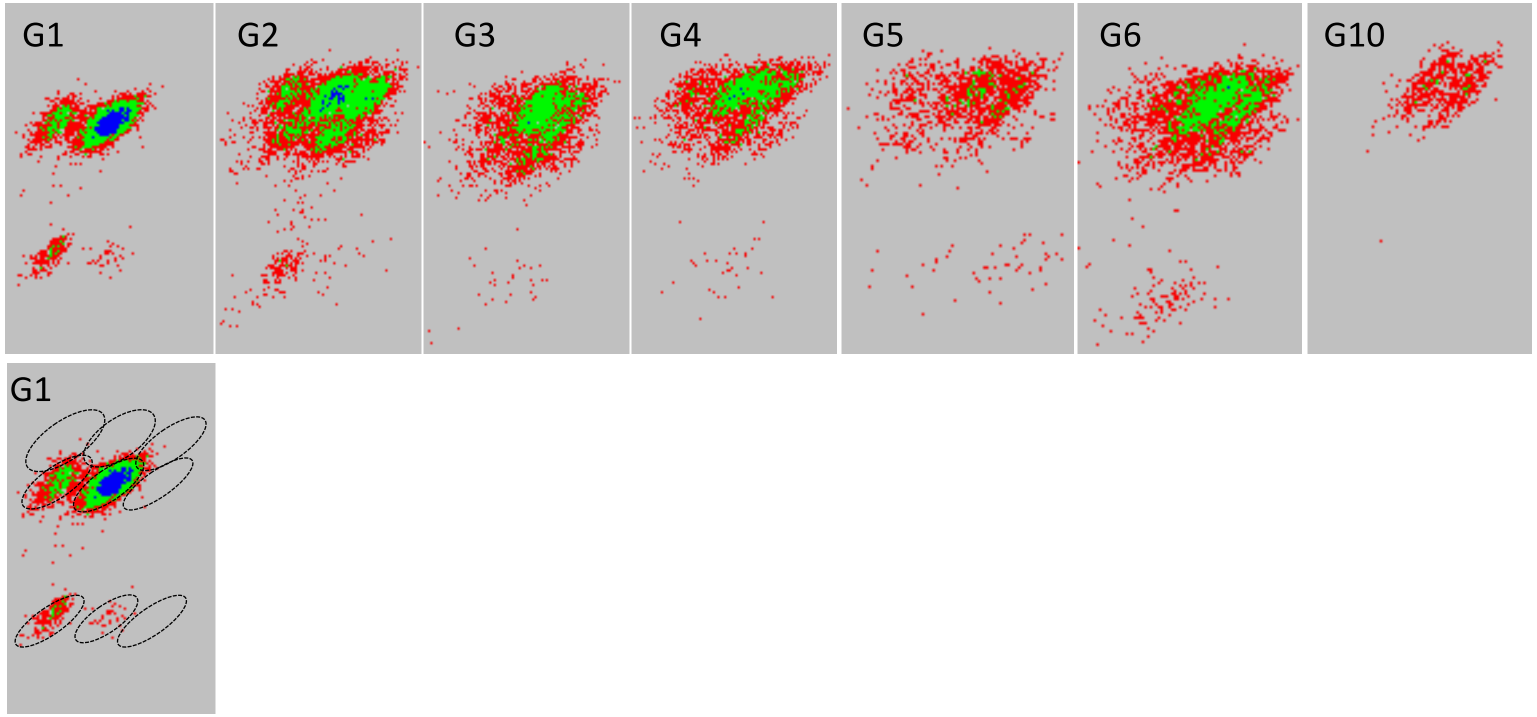
