## Supplemental Table 1 for "A population modification gene drive targeting both *Saglin* and *Lipophorin* impairs *Plasmodium* transmission in *Anopheles* mosquitoes"

| Generation | 1 | 2 | 3 | 4 | 7 | 10 | 11 | 12 | 16 | 17 |
| --- | --- | --- | --- | --- | --- | --- | --- | --- | --- | --- |
| N= | 3653 | 1714 | 1222 | 1975 | 2273 | 6852 | 9153 | 2810 | 522 | 4335 |
| % Homozygotes | 24,5 | 19,0 | 22,4 | 20,4 | 14,1 | 7,4 | 7,8 | 5,4 | 0,7 | 1,1 |
| % Heterozygotes | 49,1 | 53,9 | 46,4 | 46,2 | 47,8 | 42,9 | 39,3 | 39,5 | 21,2 | 20,5 |
| % Negatives | 26,4 | 27,1 | 31,2 | 33,4 | 38,1 | 49,7 | 53,0 | 55,1 | 78,1 | 78,3 |
| Tsg frequency | 0,49 | 0,46 | 0,46 | 0,44 | 0,38 | 0,29 | 0,27 | 0,25 | 0,11 | 0,11 |

**Supplemental Table 1A: Tracking the evolution dynamics of *Lp::Sc2A10* transgene frequency**. A parental cage was assembled containing only heterozygous transgenic mosquitoes (G0, transgene frequency =50%). Neonate larvae of subsequent generations except G5, 6, 8, 9 (N indicates the number of larvae analysed) were analyzed using COPAS flow cytometry. Gates were drawn on COPAS diagrams around clouds of larvae corresponding to homozygous, heterozygous and negative individuals according to the intensity of GFP fluorescence, and the corresponding percentage of objects in each gate was recorded. Percentages were corrected to exclude objects not corresponding to larvae. Transgene frequency per 100 chromosomes dropped from 50% in G0 to 11.35% in G17, an average loss of 2.3% per generation.

| Replicate | Number of tsg ♀ + WT ♀ in cage | Number of WT ♂ | Number of GFP^+^ progeny | Number of GFP^-^ progeny |
| --- | --- | --- | --- | --- |
| 1a | 66 + 66 | 100 | 426 (23.5 %) | 1385 |
| 1b | 66 + 66 | 100 | 378 (26.8%) | 1031 |
| 2a | 53 + 53 | 94 | 240 (10.4 %) | 2060 |
| 2b | 53 + 53 | 94 | 621 (29.6%) | 1471 |
| 3a | 140+140 | 200 | 4170 (39.7%) | 6323 |
| 3b | 140+140 | 200 | 910 (45.6%) | 1084 |
| 4a | 71+71 | 132 | 2378 (39.1 %) | 3693 |
| 4b | 71+71 | 132 | 62 (8.9 %) | 700 |

**Supplemental Table 1B: fertility tests comparing the number of progeny produced by homozygous *Lp::Sc2A10* female versus WT female mosquitoes.** Indicated identical numbers of virgin transgenic and WT females were mixed in cages with WT males. After blood feeding, neonate larvae produced by each cage were analyzed by flow cytometry (COPAS) and the numbers of GFP fluorescent and negative larvae were counted using WinMDI software on COPAS files. Identical fertility of the two categories of females would produce 50% of GFP positive progeny. a, b after the replicate number indicate the first and second egg batch, respectively, from the same mosquitoes. Replicate 1 was composed of smaller mosquitoes, due to higher density larval rearing. All other replicates were performed with mosquitoes of standard size. Replicate 4b was performed with older mosquitoes, with >50% of females already dead.
