## Supplemental Table 2 for "A population modification gene drive targeting both *Saglin* and *Lipophorin* impairs *Plasmodium* transmission in *Anopheles* mosquitoes"

**MOUSE PARASITEMIA FOLLOWING BITE-BACK EXPERIMENTS**

A. Mice bitten by homozygous *Lp::Sc2A10* mosquitoes and their wild-type sibling controls

Parasitemia in bitten mouse

| Expt. 1 (day 18) | Mosquito genotype | | day 4 | day 5 | day 9 | day 10 |
|---|---|---|---|---|---|---|
| June 29, 2019 | control | | 0.01 % | 0.04 % | 6.7 % | |
| | control | | 0.03 % | 0.24 % | | |
| | control | | 0.07 % | 0.62 % | | |
| | hmz tsg | | 0 % | 0 % | 0 % | 0 % |
| | hmz tsg | | 0 % | 0 % | 0 % | 0 % |
| | hmz tsg | | 0 % | 0 % | 0 % | 0 % |

| Expt. 2 (day 16) | Mosquito genotype | | day 4 | day 5 | day 6 | day 10 | day 14 |
|---|---|---|---|---|---|---|---|
| July 12, 2019 | control | | 0.1 % | 0.7% | 2.7 % | | |
| | hmz tsg | | 0 % | 0 % | 0 % | 0 % | 0 % |
| | hmz tsg | | 0 % | 0 % | 0 % | 0 % | 0 % |

| Expt. 3 (day 16) | Mosquito genotype | | day 5 | day 6 | day 7 | day 8 | day 9 | day 12 | day 14 |
|---|---|---|---|---|---|---|---|---|---|
| July 17, 2019 | control | | 0 % | 0 % | 0 % | 0 % | 0 % | 0 % | 0 % |
| | control | | 0.02 % | 0.15 % | 0.98% | 3.17% | 4,62% | | |
| | control | | 0.16 % | 1.01 % | 3.20% | | | | |
| | control | | 0.01 % | 0.14 % | 0.93% | 3.76% | 7,46% | | |
| | hmz tsg | | 0 % | 0 % | 0 % | 0 % | 0 % | 0 % | 0 % |
| | hmz tsg | | 0 % | 0 % | 0 % | 0 % | 0 % | 0 % | 0 % |
| | hmz tsg | D | 0 % | 0.02 % | 0.18 % | 1.14 % | 3.20 % | | |
| | hmz tsg | | 0 % | 0.01 % | 0 % | 0 % | 0 % | 0 % | 0 % |

| Expt. 4 (day 18) | Mosquito genotype | | day 4 | day 5 | day 6 | day 7 | day 10 | day 11 |
|---|---|---|---|---|---|---|---|---|
| July 19, 2019 | control | | 0 % | 0.04% | 0.30% | 2% | | |
| | control | | 0 % | 0.01% | 0.05% | 0.42% | | |
| | control | | 0 % | 0.03% | 0.32% | 1.76% | | |
| | hmz tsg | | 0.01% | 0.02 % | 0.16 % | 1.46 % | | |
| | hmz tsg | | 0 % | 0 % | 0 % | 0 % | 0 % | 0 % |
| | hmz tsg | | 0 % | 0 % | 0 % | 0 % | 0 % | 0 % |

| Expt. 5 (day 20) | Mosquito genotype | | day 4 | day 7 | day 8 | day 9 | day 10 |
|---|---|---|---|---|---|---|---|
| August 5, 2019 | control | | 0 % | 1.5 % | 1.3 % | 1.3 % | 7.8 % |
| | control | | 0 % | 1.3% | 3.3 % | 2.8 % | 4.2 % |
| | hmz tsg | | 0 % | 0 % | 0 % | 0 % | 0 % |

| Expt. 9 (day 16) | Mosquito genotype | | day 4 | day 5 | day 6 | day 7 | day 10 | day 11 | day 12 |
|---|---|---|---|---|---|---|---|---|---|
| October 4, 2019 | control | | 0.04 % | 0.4 % | 2.2 % | 5.2 % | | | |
| | control | | 0.06 % | 0.5 % | 2.5 % | 5.5 % | | | |
| | control | | 0 % | 0.1 % | 0.5 % | 2.1 % | 4.64 % | | |
| | control | | 0.02 % | 0.2 % | 1.2 % | 3.6 % | | | |
| | control | | 0.11 % | 0.8 % | 3.3 % | 4.7 % | | | |
| | hmz tsg | D | 0 % | 0 % | 0.3 % | 1.4 % | 4.66 % | | |
| | hmz tsg | D | 0 % | 0 % | 0.2 % | 0.3 % | 4.22 % | | |
| | hmz tsg | D | 0 % | 0 % | 0.2 % | 2.1 % | 6.03 % | | |
| | hmz tsg | | 0 % | 0 % | 0.1 % | 0 % | 0.04 % | 0 % | 0 % |
| | hmz tsg | | 0 % | 0 % | 0.1 % | 0.1 % | 0.01 % | 0 % | 0 % |

| Expt. 10 (day 20) | Mosquito genotype | | day 6 | day 7 | day 8 | day 9 |
|---|---|---|---|---|---|---|
| October 10, 2019 | control | | 4.54 % | | | |
| | control | | 1.18 % | 3.43% | | |
| | control | | 0.42 % | 2 % | 4.25 % | |
| | control | | 0.22 % | 1.14 % | 2.93 % | |
| | control | | 0.69 % | 2.69 % | 5/58 % | |
| | hmz tsg | | n.d. | 0 % | 0 % | 0 % |
| | hmz tsg | | n.d. | 1.17 % | 3.57 % | |
| | hmz tsg | | n.d. | 0.01 % | 0.04 % | 0 % |
| | hmz tsg | | n.d. | 0 % | 0 % | 0 % |
| | hmz tsg | | n.d. | 2.29 % | 6.1 % | |
| | hmz tsg | | n.d. | 0 % | 0.07% | 0 % |

| Expt. 11 (day 17) | | | day 6 | day 7 | day 9 | day 12 |
|---|---|---|---|---|---|---|
| June 25, 2021 | control | | 0.16 % | 1.28 % | | |
| | control | | 0.18% | 1.04% | | |
| | hmz tsg | | 0 % | 0 % | 0 % | 0 % |
| | hmz tsg | | 0.03% | 0.26 % | | |
| | hmz tsg | | 0.43% | 1.63 % | | |
| | hmz tsg | | 0 % | 0 % | 0 % | 0 % |

| Expt. 12 (day 18) | | | day 4 | day 8 | day 9 | day 12 |
|---|---|---|---|---|---|---|
| October 25, 2021 | control | 4 mosq bit | 0 % | 1.96 % | | |
| | control | | 0 % | 3.47 % | | |
| | control | | 0.06 % | 2.56 % | | |
| | hmz tsg | 5 mosq bit | 0 % | 0 % | 0 % | 0 % |

| Expt. 13 (day 18) | | | day 4 | day 5 | day 6 | day 7 | day 10 |
|---|---|---|---|---|---|---|---|
| October 29, 2021 | control | | 0.02 % | 0.11 % | 0.89 % | 2.69 % | |
| | control | | 0.02 % | 0.21 % | 1.22 % | 3.39 % | |
| | control | | 0.02 % | 0.18 % | 1.28 % | 2.94 % | |
| | hmz tsg | | 0 % | 0 % | 0 % | 0 % | 0 % |
| | hmz tsg | | 0 % | 0 % | 0 % | 0 % | 0 % |
| | hmz tsg | D | 0 % | 0 % | 0 % | 0.05 % | 8 % |
| | hmz tsg | D | 0.01 % | 0.08 % | 0.7 % | 2.32 % | |

B. Mice bitten by heterozygous *Lp::Sc2A10* mosquitoes and their wild-type sibling controls

| Expt. 6 (day 17) | | day 4 | day 5 | day 6 | day 9 | day 11 | day 12 |
|---|---|---|---|---|---|---|---|
| September 6, 2019 | control | 0 % | 0 % | 0 % | 0 % | 0 % | 0 % |
| | control | 0 % | 0.01 % | 0.06 % | 6.65 % | | |
| | control | 0.03 % | 0.22 % | 1.88% | | | |
| | het tsg | 0 % | 0 % | 0 % | 0.03 % | 0 % | 0 % |
| | het tsg | 0 % | 0 % | 0 % | 0 % | 0.03 % | 0 % |
| | het tsg | 0 % | 0 % | 0 % | 0 % | 0.03 % | 0 % |

| Expt. 7 (day 20) | | day 4 | day 7 | day 8 | day 9 | day 11 | day 12 |
|---|---|---|---|---|---|---|---|
| September 9, 2019 | control | 0 % | 0.09 % | 0.81% | 4.51% | | |
| | control | 0.02 % | 3.68 % | | | | |
| | het tsg | 0 % | 0.02 % | 0 % | 0 % | 0 % | 0 % |
| | het tsg | 0 % | 0 % | 0 % | 0 % | 0 % | 0 % |

| Expt. 12 (day 18) | | day 4 | day 8 | day 9 | day 12 |
|---|---|---|---|---|---|
| October 25, 2021 | control | 0 % | 3.57 % | | |
| | control | 0 % | 3.47 % | | |
| | control | 0.01% | 4.46 % | | |
| | het tsg | 0 % | 2.93 % | | |
| | het tsg | 0 % | 0 % | 0 % | 0 % |
| | het tsg | 0 % | 0 % | 0 % | 0 % |

| Expt. 13 (day 18) | | day 4 | day 5 | day 6 | day 7 |
|---|---|---|---|---|---|
| October 29, 2021 | control | 0.02 % | 0.16% | 0.92 % | 2.9 % |
| | control | 0.03 % | 0.32 % | 1.98 % | 3.93 % |
| | control | 0% | 0.03 % | 0.17 % | 0.84 % |
| | het tsg | 0.01 % | 0.08 % | 0.58 % | 2.59 % |
| | het tsg | 0 % | 0.03 % | 0.31 % | 1.58 % |
| | het tsg | 0.03 % | 0.18 % | 1.23 % | 3.58 % |

| Expt. 14 (day 18) | | day 5 | day 7 | day 10 | day 12 |
|---|---|---|---|---|---|
| November 19, 202 | control | 0.15 % | 3.4 % | | |
| | het tsg | 0 % | 0 % | 0 % | 0 % |

C. Mice bitten by homozygous *Lp::Sc2A10* mosquitoes and their wild-type sibling controls infected either with wild-type *P.berghei* or with *P. berghei* expressing *P. falciparum CSP*

| Expt. 8 (day 18) | | day 4 | day 7 | day 8 | day 10 | day 11 |
|---|---|---|---|---|---|---|
| September 16, 201 | control | 0 % | 0.54 % | 2.10 % | | |

| | | | | | | |
|---|---|---|---|---|---|---|
| with *P.berghei-PfCSP* | control | 0.09 % | | | | |
| | control | 0.1 % | | | | |
| | hmz tsg | 0.01 % | 0.07 % * | 0 % | 0 % | 0 % |
| | hmz tsg | 0 % | 0.03 % * | 0 % | 0 % | 0 % |
| | hmz tsg | 0 % | 0.03 % * | 0 % | 0 % | 0 % |
| | hmz tsg | 0.01 % | 0.08 % * | 0 % | 0 % | 0 % |
| with *P. berghei- WT* | control | 0.11 % | | | | |
| | control | 0.03 % | 5.58 % | | | |
| | control | 0 % | 0.32 % | 1.32 % | | |
| | control | 0.04% | 2.32 % | | | |
| | hmz tsg | 0.02 | 3.12 % | | | |
| | hmz tsg | 0.01 % | 3.76 % | | | |
| | hmz tsg | 0.01% | 2.38 % | | | |
| | hmz tsg | 0 % | 1.08 % | | | |
| | hmz tsg | 0 % | 1 % | | | |

* overerestimation due to a technical problem with the flow cytometer

D. Mice bitten by homozygous *SagGD, Lp::Sc2A10* mosquitoes and wild-type controls infected with *P. berghei* expressing *P. falciparum* CSP

| Expt. 15 (day 18) February 11, 2022 | | day 4 | day 6 | day 7 | day 10 | day 13 |
|---|---|---|---|---|---|---|
| | control | 0.02 % | 0.86% | 2.7 % | | |
| | control | 0 % | 0.23 % | 1.41 % | | |
| | control | 0 % | 0.24 % | 1.36 % | | |
| | control | 0 % | 0.08 % | 0.62 % | | |
| | hmz tsg | 0 % | 0 % | 0 % | 0 % | 0 % |
| | hmz tsg | 0 % | 0 % | 0 % | 0 % | 0 % |
| | hmz tsg | 0 % | 0 % | 0 % | 0 % | 0 % |
| | hmz tsg | 0 % | 0 % | 0 % | 0 % | 0 % |
| | hmz tsg | 0 % | 0 % | 0 % | 0 % | 0 % |
| | hmz tsg | 0 % | 0 % | 0 % | 0 % | 0 % |

E. Same as (D) but *SagGD, Lp::Sc2A10* mosquitoes blindly taken from generation 4 of population 6 (*SagGD* $^{vasa}$ not 100% homozygous)

| Exp. 16 (day 19) June 7, 2022 | | day 6 | day 7 | day 9 | day 13 |
|---|---|---|---|---|---|
| | tsg | 0 % | 0 % | 0 % | 0 % |
| | tsg | 0 % | 0 % | 0 % | 0 % |
| | tsg | 0.29 % | 1.42 % | | |
| | tsg | 0.28 % | 1.75 % | | |
| | tsg | 0 % | 0 % | 0 % | 0 % |
| | tsg | 0 % | 0 % | 0 % | 0 % |
